# Using summary data to detect and quantify ascertainment in biobanks

**DOI:** 10.64898/2026.08.02.742371

**Authors:** Babatunde S. Olasege, Adrian I. Campos, Julia Sidorenko, Tian Lin, Ciarrah-Jane S. Barry, Genona T. Maseras, Bjarni J. Vilhjálmsson, The Australian Genetics of Depression Study – Research Team, Naomi R. Wray, Valentin Hivert, Loic Yengo

## Abstract

Non-random participation in genetic studies can bias associations between genetic variants and outcomes. Existing methods to detect ascertainment bias often require individual-level data, thus limiting their broad applicability. Here, we introduce a summary-statistics-based method to detect and quantify ascertainment bias in large-scale genetic studies. Our method estimates a parameter, *θ*, which captures deviations in the mean polygenic score (PGS) of an ascertained sample relative to its expectation across non-ascertained or differentially ascertained references. We show through extensive simulations that our method is robust to population stratification and reference misspecification unlike naive mean PGS comparison. When applied to 21 traits across 11 large-scale biobanks, our method recapitulates known patterns of ascertainment and detects new evidence of ascertainment on genetic susceptibility to depression, height and blood pressure in many biobanks. Overall, our framework enables systematic assessment of ascertainment directly from summary statistics and provides a scalable tool for evaluating representativeness in large scale genetic studies.

## INTRODUCTION

Over the past 20 years, large-scale biobanks have emerged with unprecedented depth of data available in their participants. Findings from biobank studies are now shaping knowledge across various disciplines, from epidemiology to behavioural psychology.^1^ However, these discoveries can be biased by non-random participation in research studies, often driven by traits predisposing individuals to volunteer. For example, in the UK Biobank (UKB),^2^ individuals with higher genetic susceptibility to schizophrenia and Alzheimer’s disease were less likely to participate^3^ while participation in the Taiwan biobank (TWB)^4^ was skewed towards older and highly educated individuals living in urban areas.^5^ Such an ascertainment suggests that discoveries made from biobanks may not generalise to the rest of the population.

Previous studies have addressed ascertainment biases using a range of approaches.^6-8^ These approaches can be broadly classified as phenotype-based or genotype-based (**Supplementary Fig. 1**) methods. Phenotype-based methods typically infer ascertainment by comparing phenotypic distributions between study participants and a representative census-derived population. Using this approach, Fry, et al. ^7^ showed that participation in the UKB was biased towards highly-educated, leaner and taller individuals. In contrast, genotype-based methods encompass a broader set of approaches and have primarily been used to detect differential ascertainment across subgroups within a cohort. For example, Pirastu, et al. ^9^ leveraged sex differences in allele frequencies to demonstrate that participation-related traits (for example, body mass index) in the UKB differed between males and females. More recently, Benonisdottir and Kong ^6^ developed a new method to detect ascertainment by comparing trait-associated allele frequencies between sets of related and unrelated individuals in a biobank. Genotype-based methods are particularly advantageous when phenotypic data in study participants or census records are limited as genomic predictors of unmeasured traits can be leveraged instead. However, their power is also constrained by the accuracy of these predictors.

Overall, methods for detecting ascertainment often rely on individual-level data, which may be difficult to access due to privacy constraints. By contrast, summary-level data are often more accessible and, thus, provide a practical and effective alternative.^10^ Here, we introduce a summary-statistics-based method to detect and quantify ascertainment by comparing the distribution of allele frequencies between a target sample and multiple reference samples. Building on liability threshold model theory,^11^ we derive the expected allele frequency change in an ascertained sample. Through extensive simulations, we demonstrate that our method is robust to population stratification and misspecification of the reference. We applied our method to 21 complex traits and diseases in 11 large-scale biobanks and detected both expected and novel signals of ascertainment bias.

## RESULTS

### Theory and overview of methods

We model ascertainment using a liability threshold model,^11^ which has also been done in a recent study by Song and colleages.^12^ We assume an unobserved liability (*ℓ*) for participation in a study, influenced by many heritable traits, such that individuals are included in the study if their liability exceed a threshold *t*. We assume *ℓ* to be normally distributed with mean 0 and variance 1. If a trait *y* influences the probability to join the study, then any genetic predictor of that trait should be correlated with *ℓ*. We denote *R* the correlation between *ℓ* and a polygenic score (PGS) of *y*, hereafter denoted *g* for conciseness.

Overall, our strategy to detect ascertainment relative to trait *y* consists of testing if the mean of *g* in a target sample deviates from its expectation in a non-ascertained reference sample. Specifically, we seek to estimate *θ* = (E[*g*|*ℓ* > *t*] − E[*g*])/σ[*g*], where E[*g*] and σ[*g*] denote the expectation and standard deviation of *g* in a non-ascertained reference sample, respectively.

We consider two estimators of *θ*, which are both weighted averages of differences in trait-increasing allele (TIA) frequencies between a target and a reference sample. The first estimator, 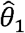, corresponds to a naïve difference in mean PGS between samples. The second estimator, 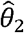, is obtained by regressing differences in TIA frequencies between the target sample and a reference onto effect sizes of TIAs (**Methods**). Importantly, 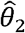 is conceptually analogous to LD score regression^13^ and, therefore, is expected to be more robust to confounding. We show through theory (**Supplementary Note 1**) and simulations (**Supplementary Fig. 2**) that both estimators have the same expectation *θ*, which can be expressed as *θ* =*iR*, where *i* = *ϕ* (*t*)/[1 − Φ(*t*)] with *ϕ* and Φ denoting the probability density function (PDF) and cumulative distribution function (CDF) of a standard Gaussian distribution, respectively. Note that parameter *i* is classically referred to as *selection intensity*.^14,15^

In practice, different combinations of *R* and *t* (and therefore, *i*) can yield the same values for *θ* which makes each parameter unidentifiable. However, given that *i* is finite, *θ*significantly different from 0 implies that *R* must also be distinct from 0. Moreover, *θ* and *R* have the same sign because *i* > 0 , such that knowing the sign of *θ* informs the direction of ascertainment. Thus, a positive (negative) value of *θ* indicates that study participants have, on average, higher (lower) genetic liability for the target trait relative to the reference sample.

In subsequent sections of the manuscript, we present and discuss the results of simulations comparing the statistical power of 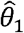 and 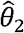, as well as their robustness to confounding and misspecification of the reference sample.

### Simulations comparing the statistical power of 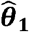 and 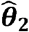

We first performed a set of simulations to determine the level of ascertainment detectable with 80% statistical power at a significant level *α* = 5% (two-sided test). We evaluated different scenarios reflecting variation in sample sizes across both the target biobank (*N*_*s*_) (i.e., the cohort where we wish to test for ascertainment) and the reference cohort (*N*_*r*_) (which provide allele frequencies from a gold standard non-ascertained cohort). For the target sample, we considered *N*_*s*_ =1,000, 15,000, and 350,000, corresponding to the approximate sample sizes of the Sweden Population Biobank (SweGen),^16^ the Australian Genomics of Depression Study (AGDS) cohort,^17^ and unrelated European ancestry individuals in the UK Biobank (UKB),^2^ respectively. For the reference sample, we varied *N*_*r*_ between 100, 500 and 1,000. Note that *N*_*r*_ = 100 is comparable to the per-population sample size in the 1000 Genome Project (1KGP) dataset.^18^

Across combinations of *N*_*s*_ and *N*_*r*_, we simulated varying strengths of ascertainment using the liability threshold model and quantified the empirical statistical power of both estimators. We found 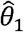 to consistently achieve higher power than 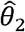, especially when the reference panel is small (**Supplementary Fig. 3**). When *N*_*r*_ = 100, a signal of approximately *θ* ≈ 0.3 was detectable with 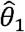, compared to *θ* ≈ 0.4 for 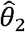. As the reference sample size increased (*N*_*r*_ ≥ 500), the performance gap narrowed and both estimators could detect signals as small as *θ* ≈ 0.15 when *N*_*r*_ = 1,000. Finally, these results also show that the size of the reference panel (as opposed to the biobank sample size, which is generally large) is the key limiting factor to detect ascertainment signal.

### Simulation to assess the impact of population stratification

#### Standard approach

Next, we sought to assess how much population structure in the target and reference samples may bias 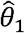 and 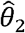 and inflate the type I error to reject the null hypothesis that *θ* = 0. Here, we simulated a scenario without any ascertainment such that any deviation of 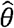 from zero may only arises because of population structure. For convenience, we hereafter refer to the type I error of 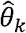 (*k* = 1 or 2), whenever 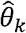 is used for hypothesis testing. In this simulation, we assumed that both target and reference are composed of two discrete sub-populations. We denote *π*_*s*_ and *π*_*r*_ as the proportions of individuals from sub-population 1 in target sample and reference, respectively. We simulated non-directional genetic differentiation between sub-populations using a Balding-Nichols model,^19^ with a fixation index *F*_ST_ ={0.001,0.011} (**Methods**). We fixed *π*_*s*_ = 0.5 and varied *π*_*r*_ between 0.1, 0.3, 0.5, 0.7 and 0.9 to simulate varying patterns of population stratification between the target sample and the reference. Importantly, *π*_*s*_ and *π*_*r*_ were only used to simulate the data and were treated as unknown in all analyses. Details about the simulations of effect sizes and allele frequencies are provided in the **Methods** section.

Across 10,000 simulation replicates, we found that both 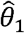 and 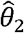 were unbiased under the null model of genetic drift, meaning that their 95% confidence intervals included zero (**Methods**; **Fig. 1a**). However, the type 1 error of 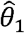 increased approximately linearly with |*π*_*s*_ = *π*_*r*_| and the degree of inflation also increased with *F*_ST_ (**Fig. 1b**). By contrast, the type 1 error of 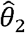 remained stable around its expected value *α* = 5% across all levels of stratification (**Fig. 1b**). A more extreme stratification scenario (**Methods**), where we artificially induced directional allele frequency differences between the two sub-samples, also resulted in a well-controlled type 1 error of 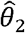 (**Fig. 1c-d**). Interestingly, the calculation of 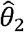 produces a secondary intercept parameter (denoted as *I*_2_, which captures residual stratification similarly to the LD score regression intercept (**Methods**). As expected, the average estimate of *I*_2_ across simulation replicates was 0 under simulated genetic drift, while it closely tracked the simulated directional effect under our more extreme model (**Supplementary Fig. 4**). Similar conclusions were obtained under scenarios with genuine ascertainment (**Supplementary Fig. 5**). Whereas 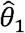 became increasingly biased in the presence of population structure, 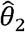 was more robust and *I*_2_ continued to tracked the simulated directional effect under our more extreme model.

**Fig 1.**
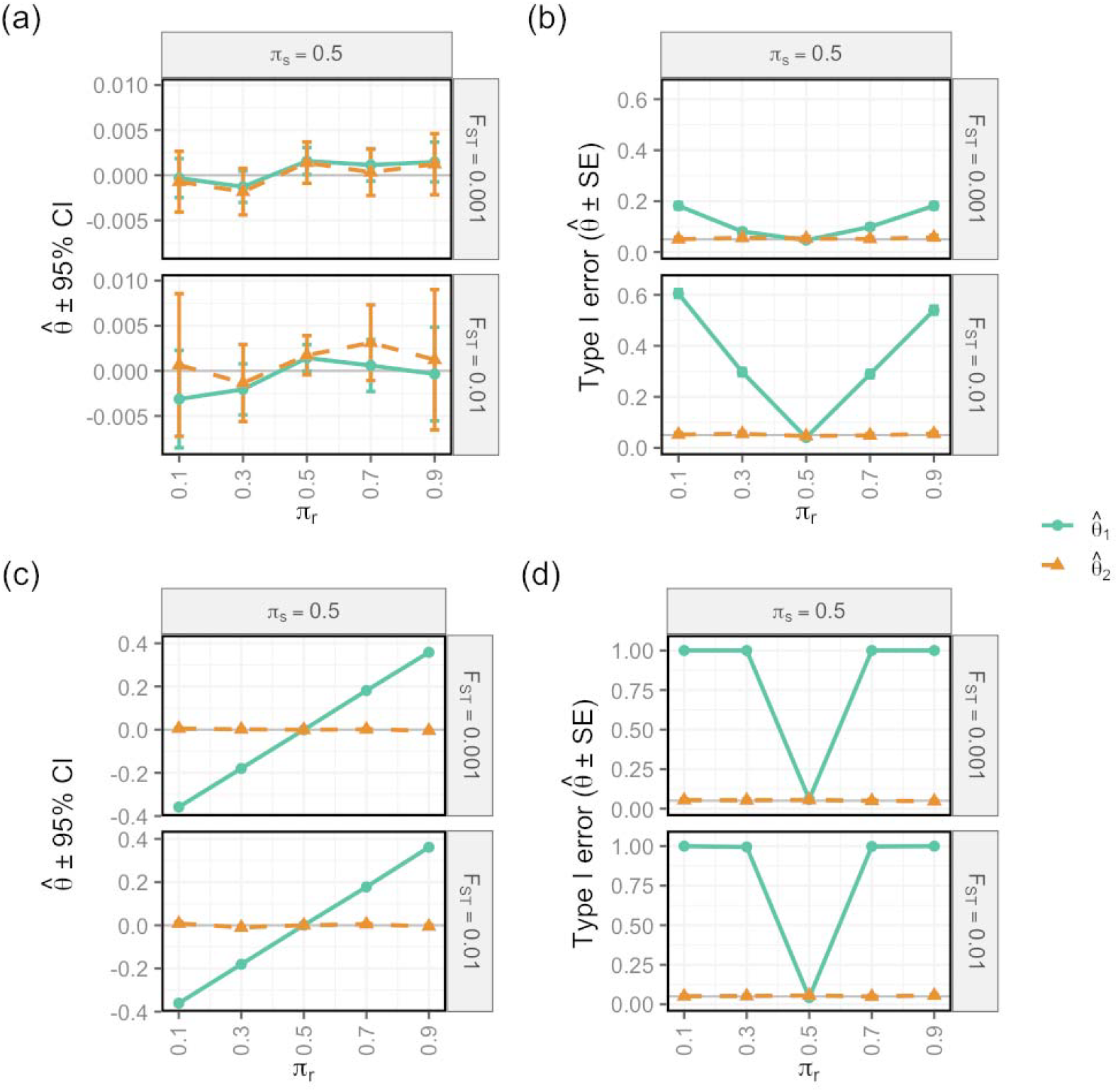
Performance 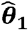 and 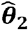 under drift (top row) and extreme (bottom row) model of population stratification across varying *F*_ST_. Panels show results from simulations where both target and reference samples consist of two sub-populations with allele frequency differences determined by *F*_ST_ ∈ {0.001, 0.01} (**Methods**). The target sample was modelled with equal ancestral loadings (*π*_*S*_ = 0.5), whereas the reference sample was simulated with either matched (*π*_*r*_ = 0.5) or skewed ancestral loadings (*π*_*r*_∈ {0.1, 0.3, 0.7, 0.9}). **Panel (a, c)** shows the mean estimates of 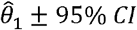 (green) and 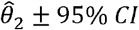 (orange) under drift (top) and extreme (bottom) model. **Panel (b, d)** shows the type I error rate of 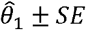 (green) and 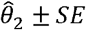 (orange) under drift (top) and extreme (bottom) model. CI represent the confidence interval estimated as 1.96±s.e.m. Type I error was calculated as the fraction of *P*-values below 0.05. SE represent standard error estimated as 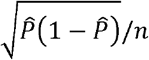, where 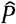 denotes the empirical type 1 error rate and *n* represent the number of replicates. Simulation was performed across 10,000 replicates.

#### Ancestry-deconvolution-based approach

The results above clearly established that inference using 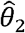 is more robust to population stratification both under the null hypothesis of no ascertainment and in the presence of genuine ascertainment signals. However, given that the statistical power to detect ascertainment is higher using a naïve PGS difference with 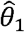, we further explored approaches that may reduce the inflated type 1 error of 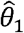 with (potentially) little impact on its statistical power. Specifically, we used an ancestry deconvolution approach previously proposed by Arriaga-MacKenzie, et al. ^20^ and Privé ^21^. Overall, this approach aims to estimate proportions of genetic ancestry from multiple references (here, *π*_*s*_) and leverages that information to build an optimal reference for calculating our parameter of interest. Intuitively, if *π*_*s*_ is known and estimated allele frequencies from the two sub-populations (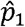 and 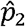) are available, then the best reference to benchmark our target sample against should have allele frequencies equal to 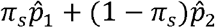 instead of 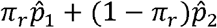 used above. Details about our implementation of that ancestry deconvolution approach are given in the **Methods** section.

Across 10,000 simulation replicates of a non-directional stratification scenario, ancestry-deconvolution substantially reduces the type I error of 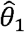, from approximately 60% to about 6% (**Fig. 2**). This improvement confirms that much of the inflation observed for 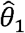 arises from mismatched ancestry proportions between the target and reference sample, and correcting for these differences can largely restore valid inference. However, a small residual inflation remained. By contrast, inference based on 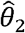 was already well-controlled and remained stable across all simulation scenarios, including both non-directional and extreme stratification scenario (**Fig. 2**). Notably, under the extreme stratification scenario, the intercept parameter *I*_2_ estimated alongside 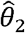 deviated systemically from zero, thus capturing the presence and the direction of residual population structure driving the inflation in 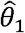 (**Supplementary Fig. 6**). A comparison of the statistical power of 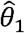 and 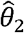 (**Supplementary Fig. 7**) still showed higher power for 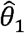, which could partly reflect its residual inflation in type I error. Overall, these additional simulations demonstrate that while ancestry deconvolution can substantially reduce the bias in 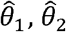 provide a more robust alternative by maintaining correct calibration and explicitly diagnosing residual stratification through *I*_2_.

**Fig 2.**
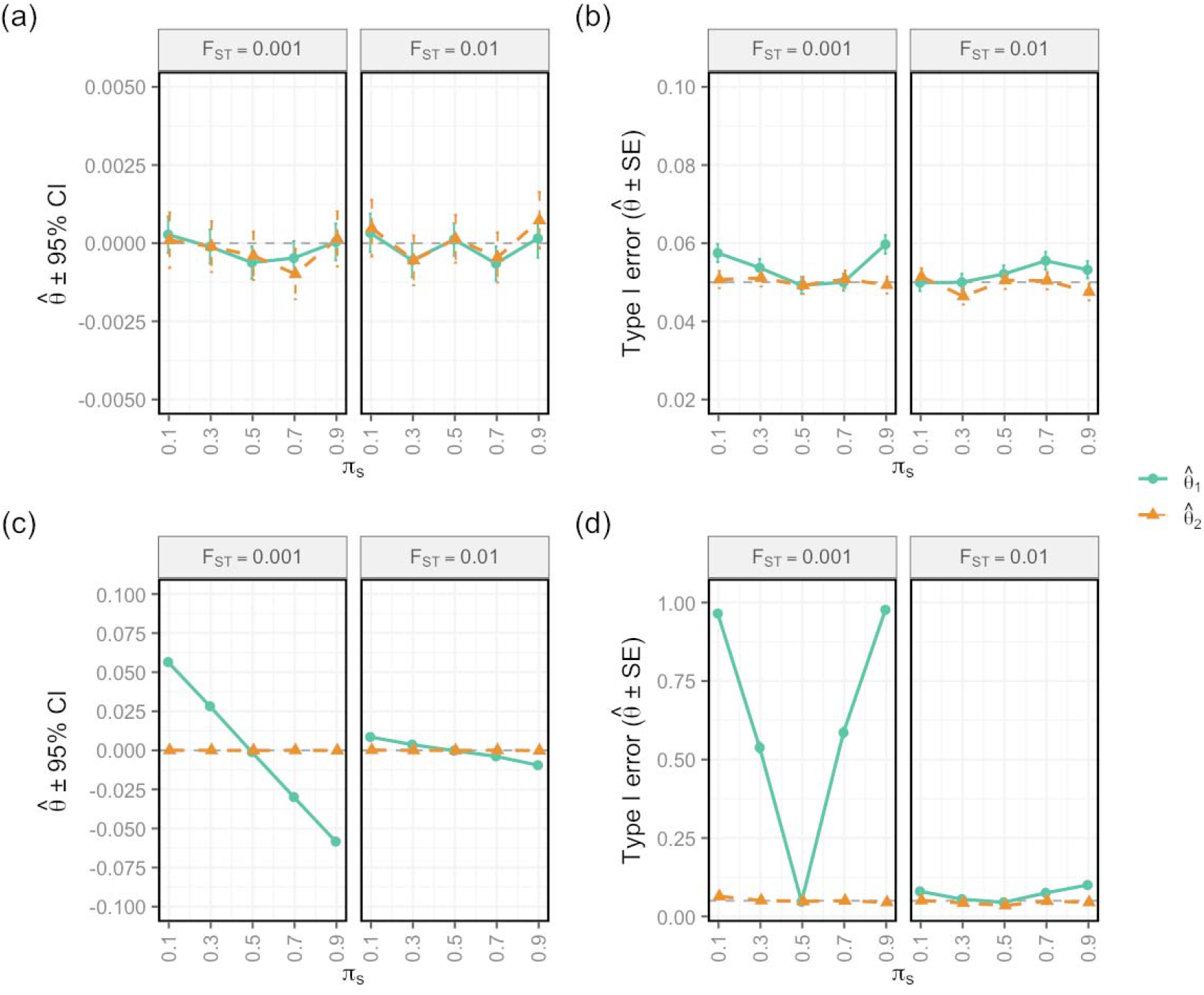
Performance of 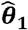 and 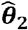 after applying the deconvolution approach under drift (top row) and extreme (bottom row) model of stratification. Panels show results from simulations where the target sample was modelled as convex linear combination of allele frequencies from two regional reference ancestries 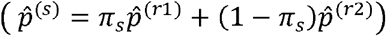 with allele frequency differences determined by *F*_ST_ ∈ {0.001, 0.01}. For each simulation settings, the ancestral loadings 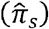 were re-estimated from sets of independent background SNPs and the estimates were used reconstruct a weighted (synthetic) reference panel (**Methods**). **Panel (a, c)** shows the mean estimates of 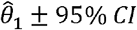 (green) and 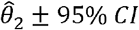 (orange) under drift (top) and extreme (bottom) model. **Panel (b, d)** shows the type I error rate of 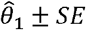 (green) and 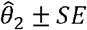 (orange) under drift (top) and extreme (bottom) model. CI represent the confidence interval estimated as 1.96±s.e.m. Type I error was calculated as the fraction of *P* values below 0.05. SE represent standard error estimated as 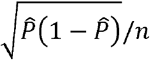, where 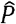 denotes the empirical type 1 error rate and *n* represent the number of replicates. Simulation was performed across 10,000 replicates.

### Simulations assessing misspecification of the reference and attenuation bias

We ran a complementary simulation study (**Methods and Supplementary Methods**) to evaluate the robustness of 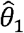 and 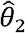 under two practical challenges. First, we assessed how each estimator performs when ancestry-deconvolution relies on a reference panel that lacks one of the ancestry groups present in the target sample. This scenario induces systematic distortions in *π*_*s*_, allowing us to assess the robustness of 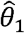 and 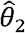 under misspecified reference. Second, we examined how sampling error in the reference biases 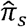 and how this attenuation bias propagates to 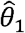 and 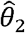.

In the first scenario, missing a reference component biased 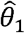 and inflated its type I error (**Supplementary Fig. 8**). In contrast, 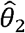 remained robust and unbiased under the same conditions. Importantly, the intercept *I*_2_ effectively absorbed the bias observed with 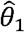, thus providing a diagnostic indicator for missing reference (**Supplementary Fig. 9**). In the second scenario, we found that sampling error in the reference biases 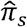, leading to misspecification of the composite reference used to benchmark the target sample, particularly when the sample size is small and *F*_ST_ is low (**Supplementary Fig. 10-11**). This bias increased with the contribution of the under-sampled reference to the target composition, resulting in substantial inflation of the type I error rate for 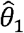 (**Supplementary Fig. 10-11**). Applying an attenuation-correction strategy mitigated this bias while maintaining the type I error of 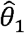 close to 5%, even when one reference was severely under-sampled (**Supplementary Fig. 10-11**). By contrast, 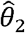 was robust to attenuation bias, maintaining well-controlled type I error rates (**Supplementary Fig. 10-11**).

### Simulations assessing impact of sample overlap

In practice, GWAS summary statistics may partially overlap with the target cohorts under investigation. This is particularly relevant for some large biobanks which may have contributed to the discovery GWAS used to construct PGSs. We therefore evaluated the impact of overlap between the GWAS discovery sample, target cohort, and reference panel. Across simulation scenarios, sample overlap did not generate spurious ascertainment signals under the null hypothesis of no ascertainment (*θ* = 0; **Supplementary Fig.12**). Under genuine ascertainment (*θ* > 0), overlap involving the target sample attenuated estimates of 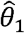 and 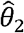, with the magnitude of attenuation increasing as the proportion of overlap increased (**Supplementary Fig. 12**). The largest attenuation was observed when overlap involved both the target and reference samples, whereas overlap restricted to the reference sample had little effect on either estimator. These findings indicate that sample overlap does not inflate ascertainment signals and, if present, is more likely to bias estimates towards the null, making our inference of ascertainment conservative.

### Application to real data

We next applied our framework to GWAS summary statistics of 21 complex traits and diseases. For each phenotype, we analysed sets of independently associated SNPs (P < 0.005; **Methods)**, restricting analyses to variants with minor allele frequency (MAF) >5% in each population. To account for ancestry-specific linkage disequilibrium (LD) structure, independent SNPs were selected separately within ancestry groups using LD clumping based on the corresponding 1KGP super-population reference panels. Given that our simulation results have shown that 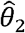 is more robust to population stratification, missing reference ancestries, and attenuation bias than 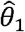, we focused our real data analyses on 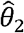 and only report results based on 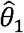 in **Supplementary Note 4** for comparison. The biobanks considered in this study span a range of recruitment strategies, including disease-specific cohorts and population-based samples, providing a diverse set of ascertainment scenarios. A structured overview of study design, recruitment, and expected ascertainment for each cohort is provided in **Supplementary Table S1**. Details of cohort recruitment, genotyping, and quality control are provided in the **Supplementary Methods**.

Importantly, interpretation of ascertainment signals is relative to the chosen reference population. We therefore first evaluated the framework in a setting where a census-representative reference is available, before extending the analysis to relative ascertainment across biobanks.

#### Validation using a census-representative reference (iPSYCH)

We evaluated the framework using the iPSYCH cohorts (iPSYCH2012: *N*_*s*_ = 40,295; iPSYCH2015: *N*_*s*_ = 30,015), which include *N*_*r*_ = 38,049 individuals randomly sampled from the population at the time of recruitment. The number of SNPs analysed for each trait is shown in **Supplementary Table S2**. Statistical significance for these validation analyses was assessed using a Bonferroni-corrected threshold of *α* = 0.05/21 = 0.0024.

Consistent with the study design, we observed strong positive associations between participation in case cohorts and genetic liability to all the psychiatric disorders studied. In particular, genetic liability to attention-deficit/hyperactivity disorder 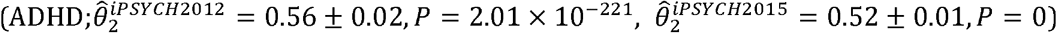 and major depressive disorder 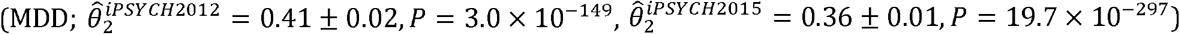 was enriched in iPSYCH cohorts relative to the reference population (**Fig. 3a**). We also observed enrichment for genetically correlated traits, including problematic alcohol use (PAU), consistent with shared genetic architecture across psychiatric disorders. These signals were consistent across the two genotyping waves, with similar effect sizes observed despite differences in statistical significance. Overall, the results demonstrate that our method captures ascertainment when the reference population is appropriately specified, providing a direct validation of the proposed framework in a near-ground-truth setting.

**Fig. 3.**
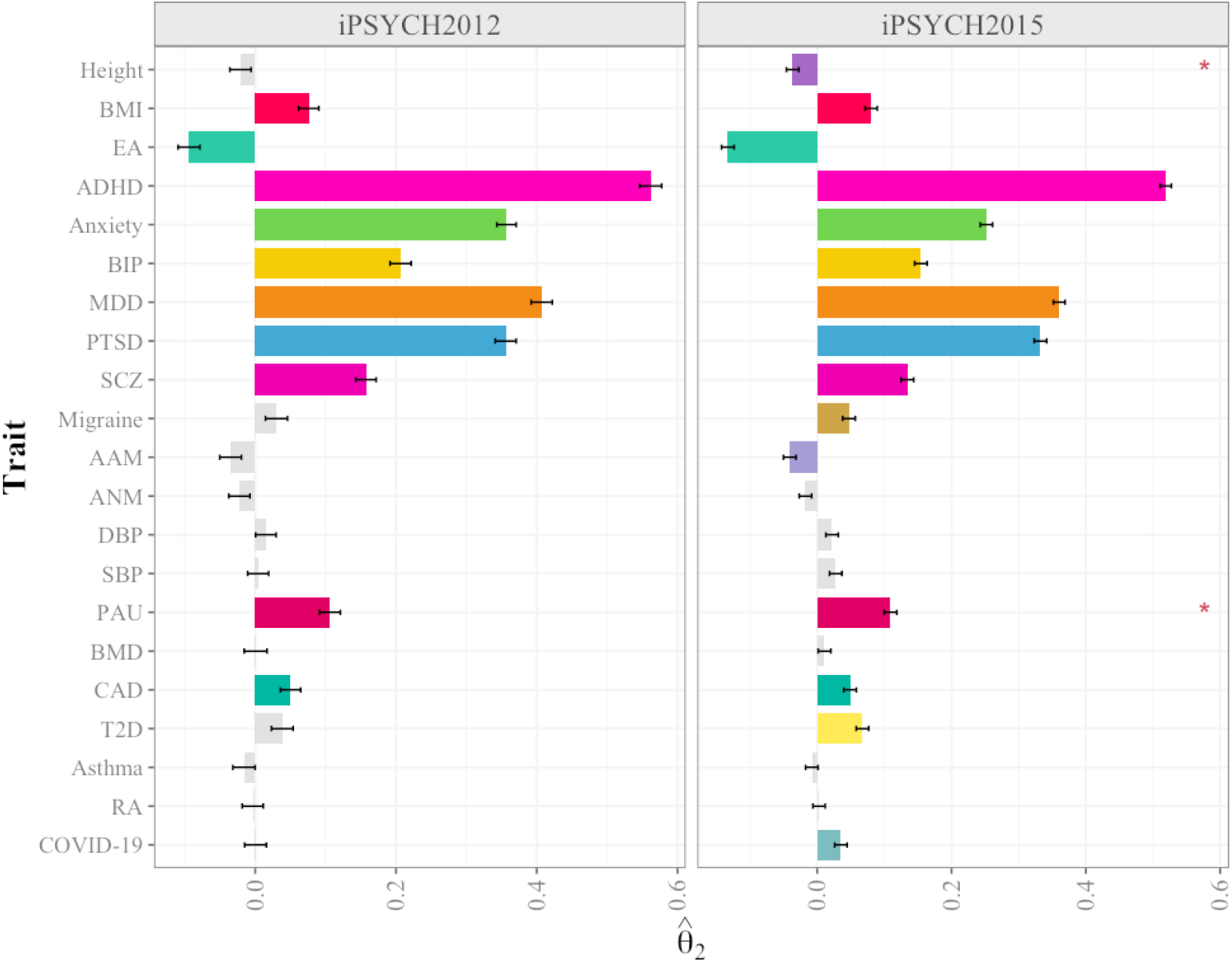
Estimates of ascertainment signals in iPSYCH cohorts using a census-representative reference. Panel shows estimates of ascertainment signals across 21 complex traits and diseases in the 2012 and 2015 waves of the Integrative Psychiatric Research (iPSYCH2012 and iPSYCH2015) biobank using a census-representative reference constructed from population controls. Bars represent point estimates of 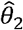 ± standard error. Grey bars denote non-significant results, while coloured bars highlight statistically significant estimates after multiple-testing correction (*P* values < 0.0024). Asterisks indicate estimates with significant intercept (*I*_*2*_; *P* values < 0.0024). Positive (negative) and statistically significant value of 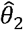 implies that the sample is enriched for individuals with higher (lower) genetic propensity or liability for the target trait compared to the reference. Only traits with ≥10 SNPs are shown.

#### Relative ascertainment across biobanks using ancestry-matched references

Given that population-representative allele frequencies were unavailable for most cohorts, we therefore investigated relative ascertainment across biobanks by comparing allele frequencies between cohorts within the same broadly defined ancestry group. Such analyses can thus inform if a biobank or cohort is differentially ascertained for a particular trait.

We analysed nine biobanks spanning diverse recruitment and sampling strategies: the Australian Genomics Depression Study (AGDS; N=14,252), Biobank Japan (BBJ; N=178,726), the Estonian biobank (EstBB; N=204,747), the Mexican biobank (MXB; N=5,663), the Mexican city prospective cohort study (MCPS; N=136,401), the children cohort (age 8) from the Norwegian Mother and Child Cohort Study (MoBa_child8; N=3,862), the Swedish population biobank (SweGen; N=1,000), the Taiwan biobank (TWB; N=92,615), and the UK Biobank (UKB; N=348,502). The cohorts were grouped according to their average genetic ancestry inferred using genetic similarity with broad ancestry groups in the 1KGP (**Supplementary Fig. 13**). We restricted cross-cohort comparisons within ancestry groups to ensure that detected signals reflect ascertainment patterns rather than other forms of selection that may exist between ancestry groups. The number of SNPs analysed for each trait across within-ancestry comparisons is provided in **Supplementary Table 3**. Statistical significance was assessed using a Bonferroni-corrected threshold accounting for all within-ancestry pairwise comparisons across the 21 investigated traits.

Within European ancestry cohorts, the strongest and most consistent ascertainment signals were observed in iPSYCH and AGDS relative to other cohorts, consistent with their recruitment strategies (**Fig. 4a-b**). Both iPSYCH2012 and iPSYCH2015 showed strong positive associations between participation and genetic liability to psychiatric traits, particularly for ADHD, MDD, anxiety and post-traumatic stress disorder (PTSD), relative to population-based European cohorts, reflecting the psychiatric case-control design of iPSYCH (**Fig. 4a**). Similarly, participation in AGDS was associated with higher genetic liability for MDD, together with enrichment for genetically correlated psychiatric phenotypes including PTSD, anxiety and ADHD, consistent with the substantial shared genetic architecture across psychiatric disorders. We did not detect significant differential ascertainment between AGDS and iPSYCH.

**Fig. 4a.**
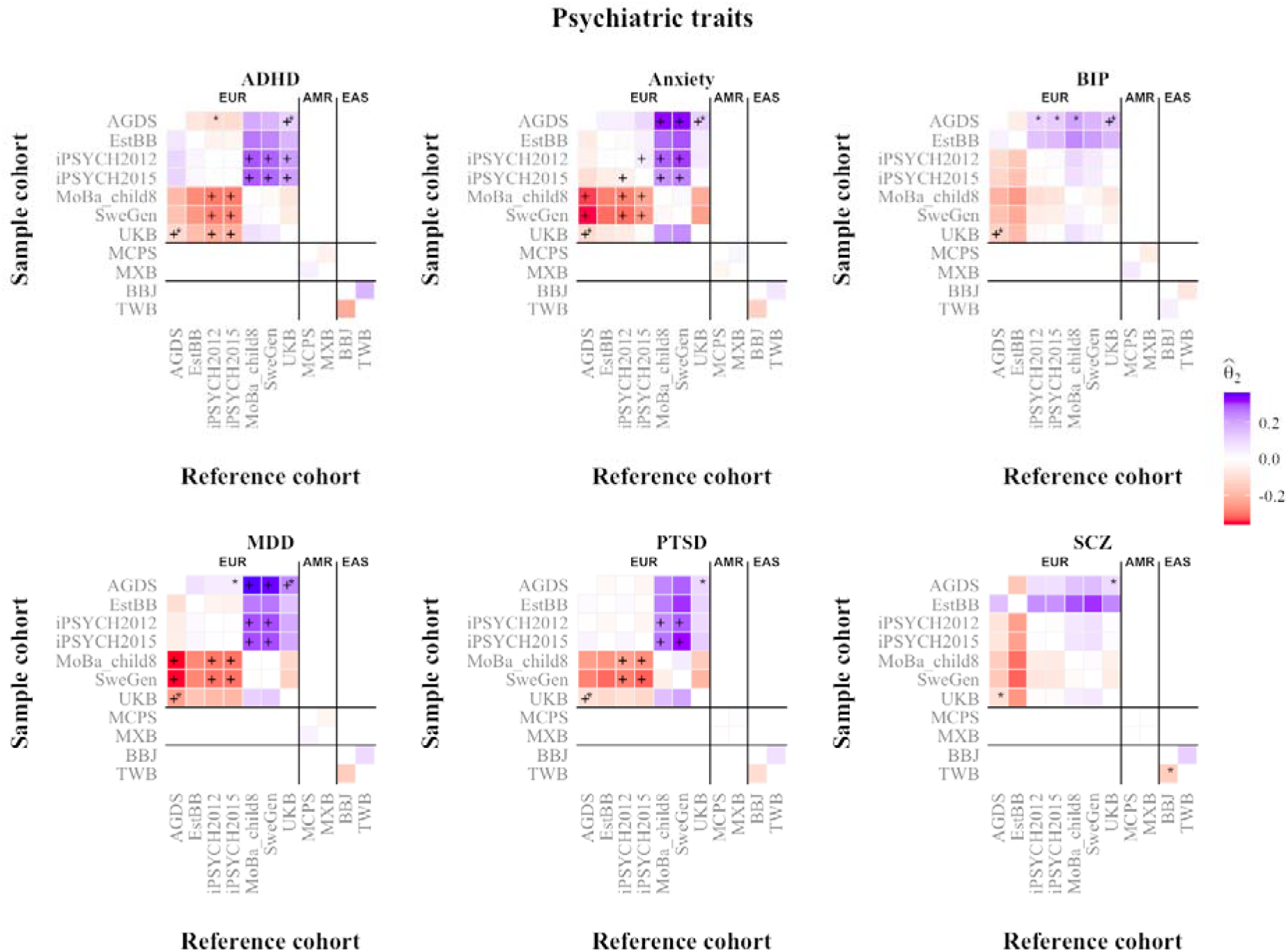
Relative ascertainment for psychiatric traits across within-ancestry biobank comparisons. Heatmaps show estimates of 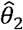 for attention deficit hyperactivity disorder (ADHD), anxiety, bipolar disorder (BIP), major depressive disorder (MDD), post-traumatic stress disorder (PTSD), and schizophrenia (SCZ) across pairwise biobank comparisons within European (EUR), admixed American (AMR), and East Asian (EAS) ancestry groups. The biobanks includes the Australian Genomics Depression Study (AGDS), Biobank Japan (BBJ), the Estonian biobank (EstBB), the 2012 and 2015 waves of the Integrative Psychiatric Research biobank (iPSYCH2012 and iPSYCH2015), the Mexican biobank (MXB), the children cohort (age 8) from the Norwegian Mother and Child Cohort Study (MoBa_child8), the Mexican city prospective cohort study (MCPS), the Swedish population biobank (SweGen), the Taiwan biobank (TWB), and the UK Biobank (UKB)). Each tile represents the estimated relative ascertainment signal between a sample cohort (rows) and a reference cohort (columns), using ancestry-matched sets of independently associated SNPs selected through LD clumping based on the corresponding 1000 Genomes Project (1KGP) super-population reference panel. Positive (negative) and statistically significant value of 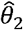 implies that the sample cohort is enriched for individuals with higher (lower) genetic propensity or liability for the target trait compared to the reference. The Plus sign (+) indicates significant 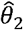 estimates (*P* values < 0.0001) after Bonferroni correction across all within-ancestry pairwise comparisons and investigated traits. Asterisks (*) indicate estimates with significant intercept (*I*_*2*_; *P* values < 0.0001) after Bonferroni correction. +* indicate when both are significant. Comparisons were restricted to cohorts belonging to the same broad ancestry group.

On the other hand, UKB, EstBB and SweGen exhibited weaker or non-significant ascertainment signals across psychiatric traits. Nevertheless, significant relative ascertainment was observed between cohorts for several non-psychiatric phenotypes, including height, systolic blood pressure (SBP) and diastolic blood pressure (DBP) (**Fig. 4b)**. Results for the remaining traits are shown in **Supplementary Fig. 14a–c**. Outside European ancestry cohorts, ascertainment signals were generally weaker and less consistent, likely reflecting the reduced predictive performance of predominantly European-derived PGSs in non-European ancestry populations.

We further evaluated the robustness and generalisability of our findings through a series of sensitivity analyses based on iPSYCH data. These additional analyses are described in **Supplementary Note 4** and (**Supplementary Fig. 15–20** and are largely concordant findings shown here.

Together, these findings suggest that cohort-specific ascertainment leaves detectable signatures on allele frequency distributions that can be recovered using summary-statistics-based approaches.

**Fig. 4b.**
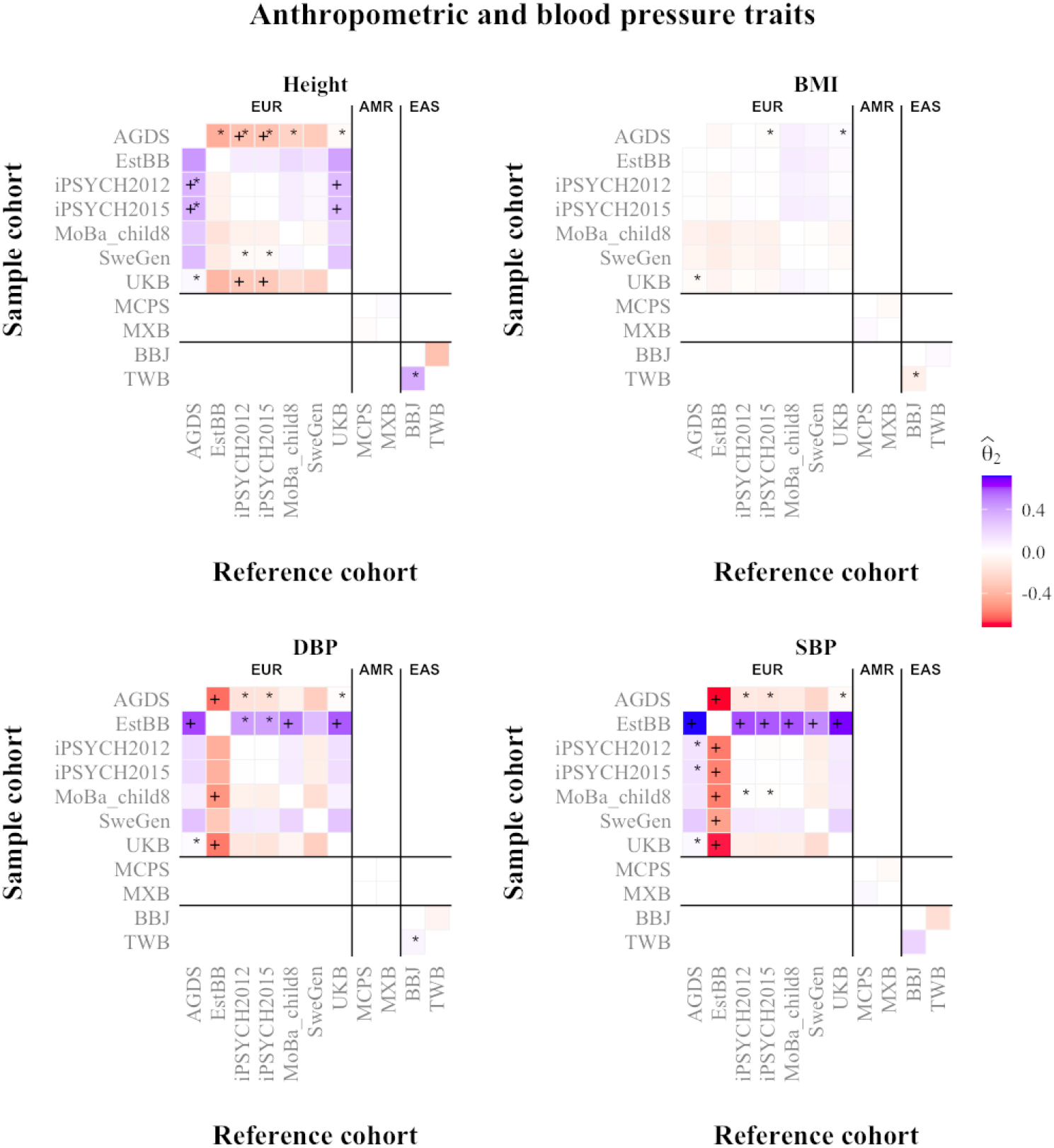
Relative ascertainment for anthropometric and blood pressure traits across within-ancestry biobank comparisons. Heatmaps show estimates of 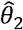 for height, body mass index (BMI), diastolic blood pressure (DBP) and systolic blood pressure (SBP) across pairwise biobank comparisons within European (EUR), admixed American (AMR), and East Asian (EAS) ancestry groups. The biobanks includes the Australian Genomics Depression Study (AGDS), Biobank Japan (BBJ), the Estonian biobank (EstBB), the 2012 and 2015 waves of the Integrative Psychiatric Research biobank (iPSYCH2012 and iPSYCH2015), the Mexican biobank (MXB), the children cohort (age 8) from the Norwegian Mother and Child Cohort Study (MoBa_child8), the Mexican city prospective cohort study (MCPS), the Swedish population biobank (SweGen), the Taiwan biobank (TWB), and the UK Biobank (UKB)). Each tile represents the estimated relative ascertainment signal between a sample cohort (rows) and a reference cohort (columns), using ancestry-matched sets of independently associated SNPs selected through LD clumping based on the corresponding 1000 Genomes Project (1KGP) super-population reference panel. Positive (negative) and statistically significant value of 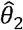 implies that the sample cohort is enriched for individuals with higher (lower) genetic propensity or liability for the target trait compared to the reference. The Plus sign (+) indicates significant 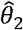 estimates (*P* values < 0.0001) after Bonferroni correction across all within-ancestry pairwise comparisons and investigated traits. Asterisks (*) indicate estimates with significant intercept (*I*_*2*_; *P* values < 0.0001) after Bonferroni correction. +* indicate when both are significant. Comparisons were restricted to cohorts belonging to the same broad ancestry group.

## DISCUSSION

This study introduces a new summary statistics-based method for detecting and quantifying trait-specific ascertainment by comparing PGS distributions between samples. Through theory and extensive simulations, we showed that a naïve comparison of mean PGS between ancestry -matched samples may still be biased by subtle population stratification within-ancestry group and proposed an alternative regression-based approach, which is robust to those artefacts. By relying solely on summary data, our approach circumvents the need for individual-level data, overcoming privacy and data sharing restrictions common to modern biobanks. More broadly, this framework opens new opportunities to systematically assess how recruitment and study design shaped the observed genetic architecture across large-scale biobanks

Using a census-representative reference in the iPSYCH cohorts, we demonstrate that ascertainment signals can be accurately recovered when the reference population is appropriately specified. In this setting, ascertainment signals closely matched expected patterns induced by psychiatric case-control sampling, while also revealing selection for additional genetically correlated traits beyond primary psychiatric phenotypes. We further recapitulated known patterns of ascertainment in iPYSCH, which was enriched with individuals with psychiatric disorders ^22,23^ or AGDS, which targeted individuals at high risk for depression ^17,24^. We additionally observed significant relative ascertainment signals for several non-psychiatric phenotypes, including height and blood pressure traits, consistent with known participation and recruitment biases in large volunteer-based cohorts.

Inference of natural selection based on mean PGS differences (i.e., 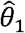) was previously criticised because of its sensitivity to population stratification ^25^. Although applied in a different context (that is, selection into a study as opposed to natural selection), our simulations and real-data analyses reinforce these previous concerns while also providing a potential solution through our second estimator, 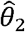. In particular, across several within-ancestry biobank comparisons, often 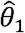 produced inflated signals that were absent under the regression-based framework 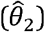 (**Supplementary Fig. 21-22**). These discrepancies were accompanied by non-zero *I*_2_ estimates, indicating systematic directional shifts in mean PGS comparisons that are consistent with either residual stratification or reference misspecification. Nevertheless, 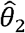 remained highly concordant with 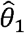 overall (**Supplementary Table S5**), suggesting that attenuation bias and residual population stratification were generally limited in the analyses considered here. Larger discrepancies between 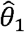 and 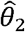 were observed when comparing cohorts from different ancestries (slope of regression of 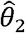 onto 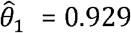; **Supplementary Table S5**). Although not the focus of this study, these observations reinforce the notion that mean PGS comparisons between populations can be misleading when subtle population structure within ancestry groups are not adequately accounted for.

One potential strategy to mitigate reference misspecification in cohorts lacking national-scale population allele frequencies like iPSYCH is to use publicly available reference resources, such as the 1KGP, together with ancestry deconvolution approaches to construct synthetic references approximating the ancestry composition of the target cohort. However, we observed evidence that the 1KGP itself carries ascertainment signatures for several complex traits. For example, educational attainment showed systematic differences across some 1KGP ancestry groups, in some cases exceeding those observed for known ascertained cohorts such as UKB (**Supplementary Fig. 23**). These findings suggest that although resources such as 1KGP are highly effective for capturing broad-scale genetic ancestry, they may not necessarily represent unbiased population baselines for analyses based on PGSs. This distinction may be particularly important for studies attempting to infer polygenic adaptation or cross-population differences in complex traits from allele-frequency contrasts alone. Therefore, our proposed estimator, 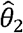, may have applications beyond detecting ascertainment, including the detection of polygenic adaptation. In fact, a previous study have attempted to use LD score regression to produce robust inference of selection by estimating the genetic correlation between a given complex trait and allele frequency differences between populations.^26^ However, these studies have been largely inconclusive because of large standard errors of genetic correlation estimates. While analogous to LD score regression, our proposed regression approach gains substantial power by focusing on variants marginally associated with the focal trait. Therefore, future research may consider extending this framework to improve the robustness of existing approaches for detecting polygenic adaptation.

Our study has several limitations. First, our framework is specifically designed to detect ascertainment when the PGS and the liability (*ℓ*) to join a study are linearly related. relationships between PGS and *ℓ*, which do not induce a shift in the mean of the PGS Therefore, our methods cannot detect ascertainment induced by nonlinear distribution. For example, a study that recruits both individuals at high genetic risk of a disease and healthy individuals with low genetic risk could enrich both tails of the PGS distribution. This would alter the distribution of PGS values among participants without necessarily changing its mean. In such cases, participation biases would primarily affect higher-order moments of the PGS distribution, and alternative methods would be required. Importantly, the absence of a directional signal in our framework should not be interpreted as evidence of random participation, but rather as an indication that ascertainment operates through non-directional mechanisms, which are beyond the scope of the present model. Second, we sometimes used allele frequencies from a subset of individuals in the biobank (e.g., Moba_child8) to reflect the general pattern of recruitment in the biobank. However, subgroups in the biobanks may exhibit different ascertainment patterns, which may limit the generalisability of some of our findings. Third, the power of our method is inherently dependent on the prediction accuracy of PGS included in our analyses. Therefore, future studies may detect patterns of ascertainment within the biobanks analysed herein, which our current analyses have missed because of low statistical power. Other factors that can influence statistical power include the fact that publicly available GWAS data can be truncated to minimize the risk to identify their participants, which increases the sampling variance of our estimates. Likewise, overlap between the GWAS discovery cohort and the target sample attenuates ascertainment estimates, biasing them towards the null rather than generating spurious ascertainment signals. Consequently, sample overlap is expected to make our inference conservative rather than inflate false-positive findings. Finally, our method does not provide a straightforward solution for correcting biases in estimates of exposure-outcome relationships within ascertained biobanks. Despite this limitation, we discuss in **Supplementary Note 5** and (**Supplementary Fig. 24-25**) how estimates of *θ* can help predict collider biases when the response rate to participation calls is known.

Overall, our study contributes to expanding the set of genome-based tools for detecting and quantifying ascertainment bias and highlights unrecognized patterns of ascertainment bias in existing publicly available datasets. As biobanks continue to expand and diversify, such tools will become essential for assessing representativeness, guiding data integration, and ensuring that genomic discoveries remain both accurate and equitable.

## METHODS

### Model Overview

Theoretically, ascertainment is expected to alter the genotype distribution of a sample relative to a non-ascertained (census-representative) reference. We adopt the classical liability threshold model ^11^ to model ascertainment. Let *ℓ*∼*N*(0,1) denote unobserved liability in the general population that is influenced by many heritable traits. We assume that individuals are included in a study when their liability exceeds some threshold *t*. Let *g* denote the PGS for a given trait *y* constructed as 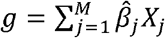, where *X*_*j*_ ∈ {0,1,2} is the trait-increasing allele (TIA) count for SNP *j* , and 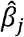 is the magnitude of effect of the TIA for SNP *j* directly observed in GWAS summary statistics, and *M* is the number of independent trait-associated SNPs included in the PGS calculation. If a trait *y* influences the probability of participation, then the PGS *g* must be correlated with *ℓ*.

We propose two equivalent equations to model the correlation between *ℓ* and *g*:

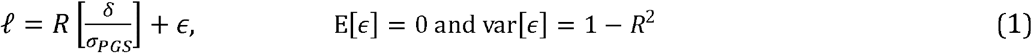

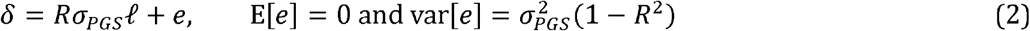

where *δ* = *g* - µ*PGS* and µ*PGS* is the mean PGS in the non-ascertained reference sample such that E[*δ*] = 0 and 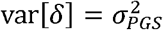.

Our strategy to detect ascertainment relative to trait *y* consists of testing if the mean of *g* in an ascertained sample deviates from *µ*_*PGS*_ Here we assume that *µ*_*PGS*_ is known. In brief, we aim to estimate *θ* = (E[*g*|*ℓ* > *t*] − E[*g*])/*σ*[*g*], where E[*g*]and *σ*[*g*]denote the expectation and standard deviation of *g* in a non-ascertained reference sample, respectively. In **Supplementary Note 1**, we show that *θ* = *iR*, where *i* = *ϕ(t)* /[1− Φ(*t*)] with *ϕ* and Φ denoting the PDF and CDF of a standard Gaussian distribution, respectively. We proposed two estimators of *θ*, which are both weighted averages of differences in TIA frequencies between a target and a reference sample.

The first estimator denoted as 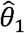 quantifies ascertainment as the standardised difference in mean PGS between samples:

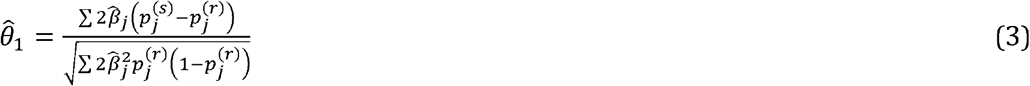

where 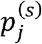 and 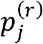 denote the TIA frequencies in the target and the reference sample, respectively. The numerator corresponds to the difference in mean PGS between the two samples, while the denominator, 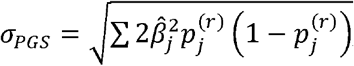, is the standard deviation of the PGS in the reference population under Hardy-Weinberg equilibrium. Standardising by *σ*_*PGS*_ yields a dimensionless measure of ascertainment that is directly comparable across traits, GWAS effect-size scales, and biobanks. Importantly, this construct ensures that 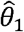 estimates the theoretical quantity *θ* = (E[*g*|*ℓ*>*t*] − E[*g*])/*σ*[*g*], derived under the liability threshold model.

The second estimator, 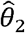, is a regression-based estimator obtained by regressing TIA frequency difference between the target and reference sample onto the effect size weighted by the heterozygosity of the reference sample. For each SNP *j*, we define the response as 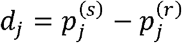 and the predictor as 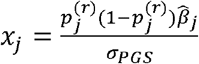 . We estimate *θ* as the slope of the regression,

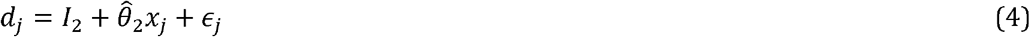

where *I*_2_ is the regression intercept. To account for heteroskedastic sampling error, each SNP is weighted by the inverse sampling variance of 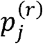 i.e., 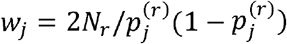, with *Nr* denoting the size of the reference sample. When *I*_2_ is constrained to zero, the estimator follows a closed-form expression 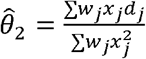. The intercept *I*_2_ captures confounding similarly to the LD score regression intercept. Consequently, 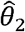 is expected to be more robust to confounding than the naïve estimator 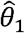.

We derived 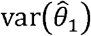 using delta approximation for the variance of a ratio (**Supplementary Note 1)** to obtain the standard error. For 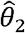, standard errors were obtained from the fitted inverse-variance-weighted regression. We then assessed the significance of the *θ* using⍰P-values from Wald tests, with the null hypothesis ‘*H*_0_ : *θ* = 0’ and the alterative hypothesis ‘‘*H*_1_ : *θ*’ ≠ 0’. Thus, a positive (resp. negative) and statistically significant value of 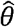 implies that the sample is enriched for individuals with higher (resp. lower) genetic liability for the target trait compared to the reference. Finally, we show in **Supplementary Note 1** that 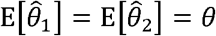 (**Supplementary Fig. 2**).

### Simulations

#### Validation of model

We conducted a simulation study to validate our proposed model and evaluate the 540 theoretical sampling variance of its estimates. First, we generated a diploid base population of *N* individuals, each genotyped at *M* =1,000 unlinked causal loci. For each SNP *j*, the TIA frequency was drawn independently as 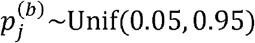 and the corresponding trait-increasing effect size as 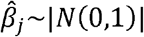. Next, we simulated the liability of each individual in the base population using the Eqn. (1), with a range of 545 different values of R. To create an ascertained sample, we selected individuals whose *ℓ*>*t*, yielding sample size of approximate *N*_*s*_ ∈ {1*K*, 15*K*, 350*K*}. The reference samples were obtained by randomly sampling individuals from the base population with varying size *N*_*r*_ ∈ {100,500,1000}. We then estimated allele frequencies 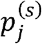 and 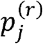 in the ascertained and reference populations, respectively. Finally, we computed the theoretical (*θ*) and empirical (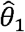 and 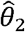) estimates and variance of our estimators across 1000 replicates to evaluate the robustness and precision of our model.

#### Impact of population stratification

Intuitively, estimates of *θ* can be confounded by population stratification when there is a systematic allele frequency difference between the target and the reference sample that are unrelated to the trait or the ascertainment process. If *θ* is computed using a reference that does not match the target sample in terms of population structure pattern, then the observed deviations may reflect ancestry differences rather than genuine ascertainment. We ran a simulation (each with 1000 replicates) under a two-way population mixture model to investigate the impact of population stratifications on 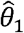 and 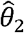. Two scenarios were considered: (i) a drift model representing non-directional stratification, and (ii) an extreme model representing directional stratification.

For the drift model, we simulated the ancestral allele frequency *p*^(0)^for *M* = 1,000 independent SNPs from a Beta (0.8, 0.8) distribution and then rescaled *p*^(0)^ so that the frequencies lie between 0.05 and 0.95. Next, we generated the allele frequencies for the two genetically drifted sub-populations for the sample and reference using a Balding– Nichols model ^19^ with *F*_ST_ = {0.001,0.01}:

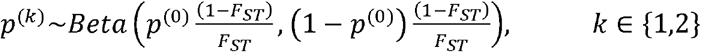

The target and reference sample were then modelled as two-way mixture of these drifted sub-populations. For each SNP, we defined

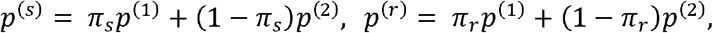

where *π*_*s*_ and *π*_*r*_ is the proportions of individuals from sub-population 1 in target sample and reference, respectively, and *p*(^1^) and *p*(^2^) represent the allele frequencies in sub-population 1 and 2. In practice, *p*(^1^) and *p*(^2^) are unknown and must be estimated from finite samples. To incorporate the sampling variance, we sampled estimates of allele frequencies for each subpopulation *k* as:

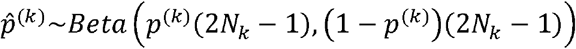

so that 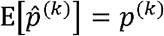 and 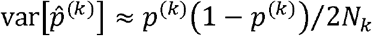. We defined 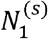 and 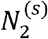 as the sample sizes of the two sub-populations in the target sample, and 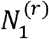 and 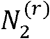 as those in the reference. The observed mixture of the allele frequencies were thus:

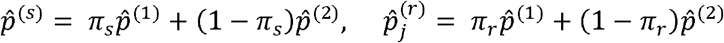

For the simulation parameters, we set total target and reference sample size to *N*_*s*_ = 5000 and *N*_*r*_ = 2500, respectively. For given mixture proportions *π*_*s*_ and *π*_*r*_, the numbers of individuals drawn from sub-populations 1 and 2 were 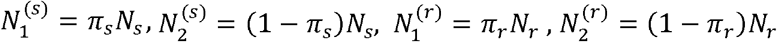. We then modelled 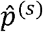 as a 50:50 mixture of two sub-populations (*π*_*s*_ = 0.5) and varied 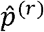 to either be a balance proportion (*π*_*r*_ = 0.5) or skewed *π*_*r*_ ∈ {0.1,0.3, 0.7,0.9}. Thus, *π*_*s*_ = *π*_*r*_ yields matched sample and reference in terms of ancestral proportion, whereas *π*_*s*_ ≠ *π*_*r*_ induce a systematic difference, 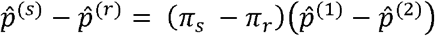, whose magnitude scale proportional with the mismatch proportions. Notably, *π*_*s*_ and *π*_*r*_ were used only in the simulation of the data and were treated as unknown parameters in all downstream analyses.

For the extreme model, we enforced a systematic directional difference for each SNP between *p*^(1)^ and *p*^(2)^ to challenge our model with a worst-case scenario of genetic structure. We simulate the two sub-populations allele frequencies as:

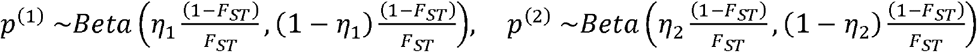

Where 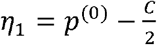 and 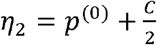. This construct guarantees a directional shift between 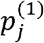 and 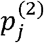 such that 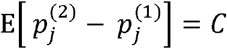. We set *C* =0.01 and perform all subsequent steps similar to the drift model: form the target and reference mixtures using *π*_*s*_ and *π*_*r*_ ; introduce finite-sample estimation by drawing 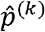 from Beta distributions parameterized by 2*N*_*k*_; and compute the observed mixture frequencies 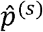 and 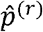. This setup yields a worst-case, directional, ancestry-driven difference in allele frequencies that inflates genetic differentiation between the target and reference samples when *π*_*s*_ ≠ *π*_*r*_.

Finally, we simulate TIA effect sizes under a polygenic model in which all variants influence the trait in the same direction. For each SNP *j*, effect sizes were drawn from a half-normal distribution 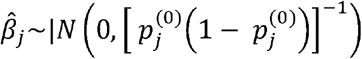 so that variants with lower MAFs exhibit larger expected variance in effect size. We then evaluated how the simulated genetic structure influences the performance of our estimators. Across 10,000 replicates, we quantified the bias and type-1 error (*α* = 0.05) on 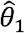 and 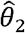 as well as *I*_2_.

#### Deconvolution to mitigate bias due to population stratification

In principle, an ancestry-matched reference should reproduce the allele frequencies mixture of the target sample. If the target ancestry proportion *π*_*s*_ were known and the allele frequencies of the two sub-populations were available, the ideal reference to benchmark the target sample would have allele frequencies 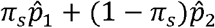 rather than the mismatched mixture 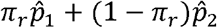. To approximate this ideal reference in practice, we implemented an ancestry deconvolution strategy ^20,21^ to reconstruct a matched reference whose allele frequencies composition reflects the ancestry mixture of the target sample. We modelled the observed allele frequencies in the target sample as a convex linear combination of allele frequencies from *K* regional reference ancestries: 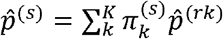, where 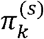 denotes the ancestral loadings of reference of reference ancestry *k* in the target sample in the, such that 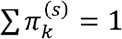 and 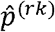 is the observed allele frequencies for ancestry k in the reference panel. In our simulation scenario settings (i.e., two subs-populations), this reduces to *k*=2 with 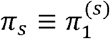 and 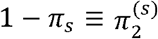.

Consider a large set of *L* independent SNPs that are sample dependent but not associated with any trait (i.e., background SNPs). To avoid confusion with trait-associated SNPs, we denote such SNPs with subscript *i*. Let 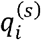 and 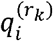 denote the estimated allele frequency of the *i*-th background SNP in the target and *k*-th reference sample, respectively. The ancestral loadings 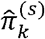 can then be estimated from 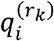 with these set of SNPs by minimizing the non-negative least square: 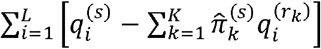. The resulting 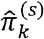 can then be used to reconstruct a new ancestry-matched reference, 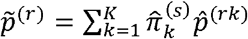. Replacing 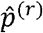 with 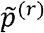 in Eqn. (3) and Eqn. (4) yields and ancestry-corrected estimate of 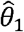 and 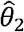. As shown in **Supplementary Note 2**, this approach is principled to mitigate the bias due to regional population structure provided that 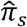 is an unbiased estimate of *π*_*s*_.

To evaluate how accurately the deconvolution strategy correct for ancestral mismatch, we ran a simulation using the drift and extreme scenario of stratification described above. In this setting, we did not simulate separate allele frequencies for the target and reference samples. Instead, we constructed the observed allele frequencies of the target sample as a mixture of two reference ancestries: 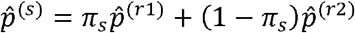. We set 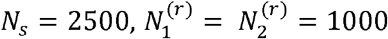, and varied *π*_*s*_ ∈ {0.1,0.3,0.5,0.7,0.9}. Next, we re-estimated 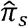 form *L* = 10,000 independent background SNPs by regressing 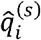 on the *k*-*th* reference allele frequencies 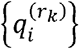. The estimated ancestral loadings were then used to reconstruct a new reference 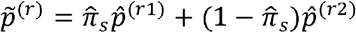 that matched the sample composition. Finally, we sampled TIA effect sizes 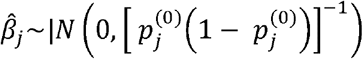 and evaluated the bias and type-1 error (*α* = 0.05) of 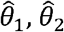 (and therefore *I*_2_).

#### Robustness of 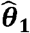 *and* 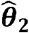 to missing reference

We further assessed the robustness of our estimators when the deconvolution approach is performed using an incomplete reference panel i.e., when one of the ancestries present in sample is missing from the reference. To mimic this situation, we simulated the target sample as a two-way mixture as above. However, in the reference panel, only 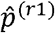 was available, mimicking a scenario with a missing ancestry. We set *N*_*s*_ = 2500, 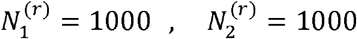, *π*_*s*_ ∈ {0.1,0.3,0.5,0.7,0.9} and sampled 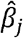 from 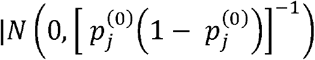. The ancestry loadings 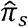 were re-estimated from *L* = 10,000 independent background SNPs by regression 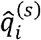 on the available reference frequencies 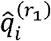. Because only one ancestry was present in the reference panel, the reconstructed reference reduced to 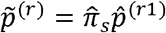. Finally, we sample 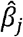 and then quantify the extent to which 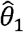, and 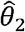 remain unbiased under the drift (non-directional) and extreme (directional) model across 10,000 replicates.

#### Robustness of 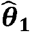 and 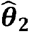 to sampling error

We next investigated the sensitivity of the estimators to sampling error in the reference allele frequencies, which could attenuate the estimated ancestry loadings 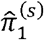 towards zero and bias the reconstructed reference during deconvolution. This is because least-squares regression assumes that the independent variables (i.e., the allele frequencies from the reference sample) are measured without error. In practice, allele frequencies are estimated from finite samples, which could introduce noise to the estimates and therefore attenuate the ancestral loadings during deconvolution. To assess the impact, we simulated the target sample as a two-way mixture: 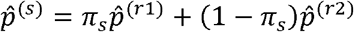. We set *N*_*s*_ = 2500, 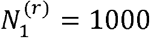, varied 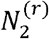 across five levels: 1000, 500, 200, 100 to represent increasing sampling noise in the second ancestry. The ancestral loading of the target sample was varied as *π*_*s*_ ∈ {0.1,0.3,0.5,0.7,0.9}. We then re-estimated 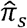 from *L* = 10,000 independent background SNPs by regressing 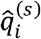 on the reference allele frequencies 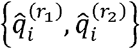 and reconstructed the new synthetic reference as 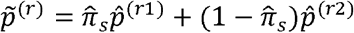. Finally, we sampled TIA effect sizes 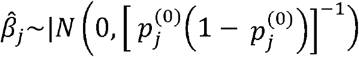 and evaluated the bias and type-1 error (*α* = 0.05) of 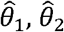 and the intercept term *I*_2_.

#### Simulations quantifying the impact of sample overlap

We performed simulations to quantify the impact of overlap between the GWAS discovery cohort and the target sample, the reference sample, or both on our methods to detect ascertainment. We simulated *M* = 1,000 unlinked causal variants with population TIA frequencies drawn from a Beta distribution and true TIA effect sizes from a normal distribution. Target and reference genotypes were sampled from the same underlying population. Individual liabilities were generated according to Eqn. (1), and individuals with *ℓ* > *t* were selected into the target sample, yielding approximately *N*_*s*_ = 5,000 ascertained individuals. The reference sample size was fixed at *N*_*r*_ = 2,500. An independent discovery cohort of 50K individuals were sampled independently of the target and reference populations. SNP effect sizes were estimated by regressing the simulated phenotype of the independent cohort on each SNP. The true causal effects used to generate the phenotype were retained as a benchmark. We then introduced overlap between the discovery cohort and the target sample, the reference sample, or both. Overlap proportions of 0%, 25%, 50%, 75%, and 100% were considered, where 0% corresponds to no overlap and 100% to complete overlap. For each configuration, randomly selected individuals from the target and/or reference samples were added to the discovery cohort and SNP effect sizes were re-estimated. Finally, we estimated 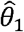 and 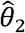 using three sets of SNP effects: the true causal effects, effects estimated from the independent discovery cohort, and effects estimated after introducing overlap. Bias due to sample overlap was quantified by comparing estimates obtained from the overlapping and non-overlapping discovery cohorts.

### Real data analyses

We applied our framework to detect and quantify ascertainment bias across multiple biobanks using two complementary strategies: (i) validation using a census-representative reference population and (ii) relative ascertainment analyses using ancestry-matched cohort comparisons.

#### Census-representative reference (iPSYCH)

To evaluate the performance of the framework under ideal conditions, we first considered a setting where a representative reference sample is available. We used allele frequencies derived from control individuals in the iPSYCH cohort, combining samples from the 2012 and 2015 genotyping waves (*N*_*r*_ = 38,049), to approximate the underlying Danish population. Ascertainment was then estimated in case samples the iPSYCH2012 (*N*_*s*_ = 40,295) and iPSYCH2015 (*N*_*s*_ = 30,015) cohorts. Using a shared from reference across cohorts ensures that differences in inferred ascertainment reflect cohort-specific recruitment rather than variation in the reference population.

#### Relative ascertainment across biobanks

We next considered a more common setting where census-representative allele frequencies are not available. In this case, we investigated relative ascertainment by directly comparing allele frequencies between cohorts within the same broad ancestry group. We analysed nine additional biobanks spanning diverse recruitment and sampling strategies: AGDS, BBJ, EstBB, MCPS, MXB, MoBa_child8, SweGen, TWB and UKB. Cohorts were grouped according to their corresponding 1KGP super-population ancestry, and analyses were restricted within ancestry groups to minimize confounding arising from broad-scale demographic differences. Allele frequencies were obtained from publicly available resources, except for iPSYCH2012, iPSYCH2015, AGDS and UKB, which were derived in-house. Detailed descriptions of cohort recruitment, genotyping and quality control are provided in the Supplementary Methods, together with a summary of study designs and expected ascertainment patterns in Supplementary Table S3.

#### GWAS summary statistics and SNP selection

We obtained GWAS summary statistics for 21 complex traits from the largest available studies. The URLs and references for all GWAS summary statistics are provided in **Supplementary Table 4**. These include GWASs on traits such as anthropometric (height and body mass index (BMI)), cognitive (educational attainment (EA)), psychiatric disorders (attention deficit hyperactivity disorder (ADHD), anxiety, bipolar disorder (BIP), major depressive disorder (MDD), and post-traumatic stress disorder (PTSD), schizophrenia (SCZ)), neurological (migraine), reproductive timing (age at menarche (AAM) and age at natural menopause (ANM)), blood pressure (diastolic blood pressure (DBP) and systolic blood pressure (SBP)), behavioural (problematic alcohol use (PAU)), skeletal (bone mineral density (BMD)) and other common diseases (coronary artery disease (CAD), type 2 diabetes (T2D), asthma, rheumatoid arthritis (RA), and COVID-19). Whenever possible, we used GWAS summary statistics that excluded the target cohort under investigation. However, complete exclusion was not always possible. In particular, some psychiatric GWAS included iPSYCH participants, while GWAS of major depressive disorder included AGDS participants. To assess the impact of such overlap, we performed simulations varying the degree of overlap between the discovery GWAS, target sample, and reference panel (**Supplementary Fig. 12**). For each trait, we selected approximately independent SNPs from the corresponding GWAS summary statistics using the LD clumping algorithm implemented in PLINK (LD squared correlation (r^2^) <⍰0.01 for SNPs⍰< ⍰1⍰Mb apart and association *P*⍰value⍰< ⍰5⍰× ⍰10^-3^). To account for ancestry-specific LD structure, SNPs were clumped separately within ancestry groups using ancestry-matched 1KGP super-population reference panels. The selected SNPs and corresponding effect sizes were subsequently subset within each cohort dataset. All datasets were harmonized prior to analysis to ensure consistent SNP matching and allele alignment across effect-size and allele-frequency files. Effect sizes were aligned to the TIA, and allele frequencies were expressed with respect to the same allele across all analyses. Standard quality-control procedures, including MAF filtering (MAF > 5%), were applied throughout (**Supplementary Methods**).

### Sensitivity analyses

We performed a series of sensitivity analyses to evaluate the robustness of our estimators using the iPSYCH dataset. First, we repeated the main analyses across the 21 traits using the marginally associated SNPs (*P* < 5 × 10^-3^) with 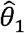 and investigated the relationship between 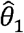 and 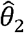. Second, we evaluated sensitive of 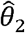 to different SNP inclusion threshold across all the studied traits. To do this, we repeated all the analyses across multiple SNP inclusion thresholds (5 × 10^-04^ up to 5 ×10^-08^), applying LD clumping at each threshold using PLINK (r^2^ < 0.01, window size = 1 Mb) with the local LD reference panel. Lastly, we evaluated the robustness of our estimators to the choice of GWAS effect sizes by repeating analyses using non-European ancestry GWAS summary statistics for representative traits, while preserving the same SNP selection framework used in the main analyses.

## DATA AVAILABILITY

AGDS: Data are available upon reasonable request (Please refer to Byrne, et al. ^24^).

BBJ: https://humandbs.biosciencedbc.jp/files/hum0197/hum0197.v3.BBJ.Hei.v1.zip

EstBB: https://www.ebi.ac.uk/gwas/studies/GCST90624701

iPYSCH: https://www.ebi.ac.uk/gwas/studies/GCST90162563

MCPS: https://www.ebi.ac.uk/gwas/studies/GCST90435342

MoBa_child8: https://www.fhi.no/en/ch/studies/moba/for-forskere-artikler/gwas-data-from-moba

MXB: https://doi.org/10.5281/zenodo.7420254

QGP: https://www.ebi.ac.uk/gwas/studies/GCST90013304

SweGen: Frequencies were obtained from the SweGen project (release 20240627), generated by Science for Life Laboratory: https://doi.org/10.17044/NBIS/G000003 ^16^

TWB: https://www.ebi.ac.uk/gwas/studies/GCST90278637

UKB: Individual-level data of UKB participants can be accessed upon application to the UKB (http://www.ukbiobank.ac.uk).

Individual level to genetic and register iPSYCH data are not publicly available due to Danish data legislation and restrictions governing access. Relevant Danish authorities may provide approval to approved research collaborations.

## CODE AVAILABILITY

Analyses were performed using publicly available software. The source code (R scripts) used to run simulations and estimate ascertainment signals can be publicly downloaded at: https://github.com/optimist0372/ascertainment-bias

## CONSORTIUM MEMBERS

We list below the members of The Australian Genetics of Depression Study (AGDS) research team and highlight in bold font those listed as authors.

**AGDS Research Team** (contributing authors on this manuscript are highlight in bold font) Enda M. Byrne (Child Health Research Centre; The University of Queensland) Jacob J. Crouse (Brain and Mind Centre; The University of Sydney) Ian B. Hickie (Brain and Mind Centre; The University of Sydney) Penelope A. Lind (Psychiatric Genetics Department, QIMR Berghofer Medical Research Institute) Nicholas G. Martin (Genetic Epidemiology Department, QIMR Berghofer Medical Research Institute) Sarah E. Medland (Psychiatric Genetics Department, QIMR Berghofer Medical Research Institute) Brittany L. Mitchell (Brain and Mental Health Program, QIMR Berghofer Medical Research Institute) Richard Parker (Brain and Mental Health Program, QIMR Berghofer Medical Research Institute) **Naomi R. Wray** (Department of Psychiatry, University of Oxford)

## ACKNOWLEGEMENTS

We thank the participants of all biobanks whose data have enabled this study. UK Biobank data was accessed under approved project ID 12505. The Australian Genetics of Depression Study (AGDS) was primarily funded by the National Health and Medical Research Council (NHMRC) of Australia (grant no. 1086683). L.Y. is supported by the Australian Research Council (grant no. FT220100069), the Snow Medical Research Foundation, and NHMRC IDEAS grant 2002694. NRW, BJV and CSB acknowledge Danish National Research Foundation, Denmark, grant number P4 supporting the Pioneer Center for Statistical and computational Methods for Advanced Research to Transform Biomedicine (SMARTbiomed). We thank Peter Visscher for critical comments and suggestions on the manuscript.

## AUTHOR CONTRIBUTIONS

L.Y. designed the study and supervised the work with critical contributions from N.R.W. B.S.O. and A.I.C. performed statistical analyses with the assistance from J.S, V.H., and T.L. B.S.O., T.L. and J.S. performed quality control of data. B.S.O. and L.Y. wrote the paper with contributions from all authors.

## COMPETING INTERESTS

A.I.C. is an employee of Regeneron Pharmaceuticals, Inc. The other authors declare no competing interests.

## REFERENCES

1. Gallagher, C.S., Ginsburg, G.S. & Musick, A. Biobanking with genetics shapes precision medicine and global health. Nature Reviews Genetics 26, 191–202 (2025).

2. Bycroft, C. et al. The UK Biobank resource with deep phenotyping and genomic data. Nature 562, 203–209 (2018).

3. Tyrrell, J. et al. Genetic predictors of participation in optional components of UK Biobank. Nature communications 12, 886 (2021).

4. Feng, Y. et al. Taiwan Biobank: A rich biomedical research database of the Taiwanese population. Cell Genom. 2, 100197. (2022).

5. Wu, C.-S. et al. Comparison of demographic and clinical characteristics of Taiwan biobank participants with nonparticipants. Journal of Epidemiology 35, 206–211 (2025).

6. Benonisdottir, S. & Kong, A. Studying the genetics of participation using footprints left on the ascertained genotypes. Nature Genetics 55, 1413–1420 (2023).

7. Fry, A. et al. Comparison of sociodemographic and health-related characteristics of UK Biobank participants with those of the general population. American journal of epidemiology 186, 1026–1034 (2017).

8. Shapland, C.Y., Gkatzionis, A., Hemani, G. & Tilling, K. Use of genetic correlations to examine selection bias. Genetic Epidemiology 49, e22584 (2025).

9. Pirastu, N. et al. Genetic analyses identify widespread sex-differential participation bias. Nat Genet 53, 663–671 (2021).

10. Pasaniuc, B. & Price, A.L. Dissecting the genetics of complex traits using summary association statistics. Nature reviews genetics 18, 117–127 (2017).

11. Falconer, D.S. The inheritance of liability to certain diseases, estimated from the incidence among relatives. Annals of human genetics 29, 51–76 (1965).

12. Song, S., Benonisdottir, S., Liu, J.S. & Kong, A. Participation bias in the estimation of heritability and genetic correlation. Proceedings of the National Academy of Sciences 122, e2425530122 (2025).

13. Bulik-Sullivan, B.K. et al. LD Score regression distinguishes confounding from polygenicity in genome-wide association studies. Nat Genet 47, 291–5 (2015).

14. Falconer, D.S. Introduction to quantitative genetics, (Pearson Education India, 1996).

15. Lynch, M. & Walsh, B. Genetics and analysis of quantitative traits, (Sinauer Sunderland, MA, 1998).

16. Ameur, A. et al. SweGen: a whole-genome data resource of genetic variability in a cross-section of the Swedish population. European Journal of Human Genetics 25, 1253–1260 (2017).

17. Mitchell, B.L. et al. The Australian Genetics of Depression Study: new risk loci and dissecting heterogeneity between subtypes. Biological Psychiatry 92, 227–235 (2022).

18. Consortium, G.P. A global reference for human genetic variation. Nature 526, 68 (2015).

19. Balding, D.J. & Nichols, R.A. A method for quantifying differentiation between populations at multi-allelic loci and its implications for investigating identity and paternity. Genetica 96, 3–12 (1995).

20. Arriaga-MacKenzie, I.S. et al. Summix: a method for detecting and adjusting for population structure in genetic summary data. The American Journal of Human Genetics 108, 1270–1282 (2021).

21. Privé, F. Using the UK Biobank as a global reference of worldwide populations: application to measuring ancestry diversity from GWAS summary statistics. Bioinformatics 38, 3477–3480 (2022).

22. Bybjerg-Grauholm, J. et al. The iPSYCH2015 Case-Cohort sample: updated directions for unravelling genetic and environmental architectures of severe mental disorders. MedRxiv, 2020.11. 30.20237768 (2020).

23. Pedersen, C.B. et al. The iPSYCH2012 case–cohort sample: new directions for unravelling genetic and environmental architectures of severe mental disorders. Molecular psychiatry 23, 6–14 (2018).

24. Byrne, E.M. et al. Cohort profile: the Australian genetics of depression study. BMJ open 10, e032580 (2020).

25. Sohail, M. et al. Polygenic adaptation on height is overestimated due to uncorrected stratification in genome-wide association studies. Elife 8(2019).

26. Berg, J.J. et al. Reduced signal for polygenic adaptation of height in UK Biobank. Elife 8, e39725 (2019).

