## Supplementary Information for "Using summary data to detect and quantify ascertainment in biobanks"

### **SUPPLEMENTARY NOTES**

#### **Supplementary Notes 1: Theoretical framework and derivation of ascertainment model**

##### 1.1. Model description and notations

We model ascertainment using a liability threshold model. We consider an unobserved liability ($\mathcal{l}$), which is determined by many heritable traits and such that individuals in the study have their underlying liability values larger than a threshold $t$. We assume that $\mathcal{l}$ is normally distributed with mean 0 and variance 1. If a trait $y$ influences the probability to join the study, then any genetic predictor of that trait should be correlated with $\mathcal{l}$. We denote $R$ as the correlation between $\mathcal{l}$ and a polygenic score (hereafter denoted $g$) of $y$.

We propose two equivalent equations to model the correlation between $\mathcal{l}$ and $g$:

$$\left( 1.1 \right)\mathcal{l=}R\left[ \frac{g-\mu_{PGS}}{\sigma_{PGS}} \right]+\epsilon$$

where $\epsilon$ is independent of $g$ such that $E\left[ \epsilon\right]=0$ and $\mathrm{var}\left[ \epsilon\right]=1-R^{2}$, and

$$\left( 1.2 \right) \delta=\left( g-\mu_{PGS} \right)=R\sigma_{PGS}\mathcal{l+}e$$

where $e$ independent of $\mathcal{l}$, $E\left[ e \right]=0$ and $\mathrm{var}\left[ e \right]=\sigma_{PGS}^{2}\left( 1-R^{2} \right)$, consistent with $E\left[ \delta\right]=0$ and $\mathrm{var}\left[ \delta\right]=\sigma_{PGS}^{2}$.

Importantly, all expectations and variances mentioned above are defined relative to a randomly mating and unascertained population. Our strategy to detect ascertainment relative to trait $y$ consists of testing if the mean PGS of $y$ in an ascertained sample deviate from $\mu_{PGS}$. Here we assume that $\mu_{PGS}$ is known. We describe further below how $\mu_{PGS}$ is obtained from a reference sample.

In the next sub-sections derive the expectations and variances of $\delta$ in an ascertained sample.

##### 1.2. Mean and variance of PGS in ascertained samples

Eqn. (1.2) implies that

$$\left( 1.3 \right) E\left[ \delta| \mathcal{l>}t \right]=R\sigma_{PGS}E\left[ \mathcal{l} | \mathcal{l>}t \right]+E\left[ e | \mathcal{l>}t \right]$$

$$\left( 1.3 \right) E\left[ \delta| \mathcal{l>}t \right]=\boldsymbol{iR}\boldsymbol{\sigma}_{\boldsymbol{PGS}}$$

where $i=E\left[ \mathcal{l} | \mathcal{l>}t \right]$, which is also referred to as *selection intensity*.

Similarly, using truncated normal distribution theory, we can write that

$$\left( 1.4 \right) \mathrm{var}\left[ \delta| \mathcal{l>}t \right]=R^{2}\sigma_{PGS}^{2}\mathrm{var}\left[ \mathcal{l} | \mathcal{l>}t \right]+\mathrm{var}\left[ e | \mathcal{l>}t \right]=R^{2}\sigma_{PGS}^{2}\left[ 1-i\left( i-t \right) \right]+\sigma_{PGS}^{2}\left( 1-R^{2} \right)$$

$$\left( 1.4 \right) \mathrm{var}\left[ \delta| \mathcal{l>}t \right]=\boldsymbol{\sigma}_{\boldsymbol{PGS}}^{\boldsymbol{2}}\left[ \boldsymbol{1-}\boldsymbol{R}^{\boldsymbol{2}}\boldsymbol{i}\left( \boldsymbol{i-t} \right) \right]$$

##### 1.3. Changes in allele and genotype frequencies in ascertained samples

We now assume that the PGS $g$ is calculated from a set of $M$ independent SNPs. We denote $p_{j}$ and $\hat{\beta}_{j}$ as the frequency and estimated effect size of the trait-increasing allele at SNP $j$, respectively.

Therefore, we can express $g$ as

$$\left( 1.5 \right) g=\sum_{j=1}^{M} \hat{\beta}_{j}X_{j}$$

and rewrite Eqn. (1.1) as

$$\left( 1.6 \right)\mathcal{l=}\frac{R}{\sigma_{PGS}}\left[ \sum_{j=1}^{M} \hat{\beta}_{j}(X_{j}-2p_{j}) \right]+\epsilon$$

where $\mu_{PGS}=\sum_{j=1}^{M} 2p_{j}\hat{\beta}_{j}$ and $\sigma_{PGS}^{2}=\sum_{j=1}^{M} 2p_{j}(1-p_{j})\hat{\beta}_{j}^{2}$

For simplicity we denote $h_{j}= 2p_{j}(1-p_{j})$ and $q_{j}^{2}={h_{j}\hat{\beta}}_{j}^{2}R^{2}/\sigma_{PGS}^{2}$.

We can rewrite $\mathcal{l}$ as

$$\left( 1.7 \right)\mathcal{l=}\frac{R}{\sigma_{PGS}}\left[ \hat{\beta}_{j}\left( X_{j}-2p_{j} \right) \right]+\mathcal{l}_{-j} \Longleftrightarrow\delta_{j}=\hat{\beta}_{j}\left( X_{j}-2p_{j} \right)=\frac{\sigma_{PGS}}{R}\left[ \mathcal{l-}\mathcal{l}_{-j} \right]$$

where $E\left[ \mathcal{l}_{-j} \right]=0$ and $\mathrm{var}\left[ \mathcal{l}_{-j} \right]=1-q_{j}^{2}$ and $\mathrm{corr}\left( \mathcal{l,}\mathcal{l}_{-j} \right)=\sqrt{1-q_{j}^{2}}$

Eqn. (1.7) implies that

$$\left( 1.8 \right) E[\delta_{j}\mathcal{|l>}t]=\frac{\sigma_{PGS}}{R}\left[ E\mathcal{[l|l>}t]-E[\mathcal{l}_{-j}\mathcal{|l>}t] \right]$$

We can express $\mathcal{l}_{-j}$ as a function of $\mathcal{l}$ as

$$\left( 1.9 \right) \mathcal{l}_{-j}=\frac{\mathrm{cov}\left( \mathcal{l,}\mathcal{l}_{-j} \right)}{\mathrm{var}\left[ \mathcal{l} \right]}\mathcal{l+}e_{j}^{'}=\left( 1-q_{j}^{2} \right)\mathcal{l+}e_{j}^{'}$$

where $E\left[ e_{j}^{'} \right]=0$ and $\mathrm{var}\left[ e_{j}^{'} \right]=\mathrm{var}\left[ \mathcal{l}_{-j} \right]-\left( 1-q_{j}^{2} \right)^{2}\mathrm{var}\left[ \mathcal{l} \right]=\left( 1-q_{j}^{2} \right)-\left( 1-q_{j}^{2} \right)^{2}=q_{j}^{2}\left( 1-q_{j}^{2} \right)$ and $\mathrm{cov}\left( \mathcal{l,}e_{j}^{'} \right)=0$ by design. Therefore, $E\left[ \mathcal{l}_{-j} | \mathcal{l>}t \right]=\left( 1-q_{j}^{2} \right)E\mathcal{[l|l>}t]$, which further implies that

$$\left( 1.10 \right) E[\delta_{j}\mathcal{|l>}t]=\frac{\sigma_{PGS}}{R}i\left[ 1-\left( 1-q_{j}^{2} \right) \right]=\frac{\sigma_{PGS}}{R}iq_{j}^{2}=\left( \frac{{h_{j}\hat{\beta}}_{j}^{2}}{\sigma_{PGS}} \right)iR$$

Equation (1.10) also implies that the expected allele frequency $p_{j}^{\left( s \right)}=E\left[ X_{j} | \mathcal{l>}t \right]/2$ in the ascertained sample can be expressed as $\boldsymbol{p}_{\boldsymbol{j}}^{\left( \mathbf{s} \right)}\boldsymbol{=}\boldsymbol{p}_{\boldsymbol{j}}\boldsymbol{+}\left[ \frac{\boldsymbol{p}_{\boldsymbol{j}}\boldsymbol{(1-}\boldsymbol{p}_{\boldsymbol{j}}\boldsymbol{)}{\hat{\boldsymbol{\beta}}}_{\boldsymbol{j}}}{\boldsymbol{\sigma}_{\boldsymbol{PGS}}} \right]\boldsymbol{iR}$

$$\left( 1.11 \right) \boldsymbol{p}_{\boldsymbol{j}}^{\left( \mathbf{s} \right)}\boldsymbol{-}\boldsymbol{p}_{\boldsymbol{j}}\boldsymbol{=iR}\left[ \frac{\boldsymbol{p}_{\boldsymbol{j}}\boldsymbol{(1-}\boldsymbol{p}_{\boldsymbol{j}}\boldsymbol{)}{\hat{\boldsymbol{\beta}}}_{\boldsymbol{j}}}{\boldsymbol{\sigma}_{\boldsymbol{PGS}}} \right]$$

$$\left( \boldsymbol{1.12} \right) \mathbf{var.}\boldsymbol{=iR}\left[ \frac{\boldsymbol{p}_{\boldsymbol{j}}\boldsymbol{(1-}\boldsymbol{p}_{\boldsymbol{j}}\boldsymbol{)}{\hat{\boldsymbol{\beta}}}_{\boldsymbol{j}}}{\boldsymbol{\sigma}_{\boldsymbol{PGS}}} \right]$$

Finally,

$$\left( 1.12 \right) \mathrm{var}\left[ \delta_{j}\mathcal{|l>}t \right]=\hat{\beta}_{j}^{2} \mathrm{var}\left[ X_{j}\mathcal{|l>}t \right]=\mathrm{var}\left[ \frac{\sigma_{PGS}}{R}\left( q_{j}^{2}\mathcal{l+}e_{j}^{'} \right)\mathcal{|l>}t \right]$$

$$\left( 1.12 \right) \mathrm{var}\left[ \delta_{j}\mathcal{|l>}t \right]=\frac{\sigma_{PGS}^{2}}{R^{2}}\left[ q_{j}^{4}\mathrm{var}\left[ \mathcal{l} | \mathcal{l>}t \right]+q_{j}^{2}\left( 1-q_{j}^{2} \right) \right]$$

$$\left( 1.12 \right) \mathrm{var}\left[ \delta_{j}\mathcal{|l>}t \right]=\frac{\sigma_{PGS}^{2}}{R^{2}}q_{j}^{2}\left[ q_{j}^{2}[1-i(i-t)]+1-q_{j}^{2} \right]$$

$$\left( 1.12 \right) \mathrm{var}\left[ \delta_{j}\mathcal{|l>}t \right]=\frac{\sigma_{PGS}^{2}}{R^{2}}q_{j}^{2}\left[ 1-q_{j}^{2}i\left( i-t \right) \right]$$

$$\left( 1.12 \right) \mathrm{var}\left[ \delta_{j}\mathcal{|l>}t \right]={\boldsymbol{h}_{\boldsymbol{j}}\boldsymbol{\beta}}_{\boldsymbol{j}}^{\boldsymbol{2}}\left[ \boldsymbol{1-}\boldsymbol{q}_{\boldsymbol{j}}^{\boldsymbol{2}}\boldsymbol{i}\left( \boldsymbol{i-t} \right) \right]$$

Therefore,

$\left( 1.13 \right) \mathrm{var}\left[ X_{j}\mathcal{|l>}t \right]=h_{j}\left[ 1-q_{j}^{2}i\left( i-t \right) \right]=\boldsymbol{h}_{\boldsymbol{j}}\left[ \boldsymbol{1-}\left( \frac{\boldsymbol{h}_{\boldsymbol{j}}{\hat{\boldsymbol{\beta}}}_{\boldsymbol{j}}^{\boldsymbol{2}}}{\boldsymbol{\sigma}_{\boldsymbol{PGS}}^{\boldsymbol{2}}} \right)\boldsymbol{R}^{\boldsymbol{2}}\boldsymbol{i}\left( \boldsymbol{i-t} \right) \right]$.

We further expand our derivations to obtain an expression for $\mathrm{cov}[\delta_{j},\delta_{k}\mathcal{|l>}t]$.

We first derive $E[\delta_{j}\delta_{k}\mathcal{|l>}t]$:

$$\left( 1.14 \right) E\left[ \delta_{j}\delta_{k}\mathcal{|l>}t \right]=\frac{\sigma_{PGS}^{2}}{R^{2}}E\left[ (q_{j}^{2}\mathcal{l+}e_{j}^{'})(q_{k}^{2}\mathcal{l+}e_{k}^{'})\mathcal{|l>}t \right]$$

$$\left( 1.14 \right) E\left[ \delta_{j}\delta_{k}\mathcal{|l>}t \right]=\frac{\sigma_{PGS}^{2}}{R^{2}}\left( q_{j}^{2}q_{k}^{2}E\left[ \mathcal{l} | \mathcal{l>}t \right]+E[e_{j}^{'}e_{k}^{'}] \right)$$

$$\left( 1.14 \right) E\left[ \delta_{j}\delta_{k}\mathcal{|l>}t \right]=\frac{\sigma_{PGS}^{2}}{R^{2}}\left[ q_{j}^{2}q_{k}^{2}(1+it)+E[e_{j}^{'}e_{k}^{'}] \right]$$

We show in **Appendix A** that $E\left[ e_{j}^{'}e_{k}^{'} \right]=-q_{j}^{2}q_{k}^{2}$. Therefore,

$$\left( 1.15 \right) E\left[ \delta_{j}\delta_{k}\mathcal{|l>}t \right]=\frac{\sigma_{PGS}^{2}}{R^{2}}q_{j}^{2}q_{k}^{2}it$$

It then follows that

$$\left( 1.16 \right) \mathrm{cov}\left[ \delta_{j},\delta_{k} | \mathcal{l>}t \right]=E\left[ \delta_{j}\delta_{k}\mathcal{|l>}t \right]-E[\delta_{j}\mathcal{|l>}t]E[\delta_{k}\mathcal{|l>}t]$$

$$\left( 1.16 \right) \mathrm{cov}\left[ \delta_{j},\delta_{k} | \mathcal{l>}t \right]=\frac{\sigma_{PGS}^{2}}{R^{2}}q_{j}^{2}q_{k}^{2}it-\left( \frac{\sigma_{PGS}}{R}iq_{j}^{2} \right)\left( \frac{\sigma_{PGS}}{R}iq_{k}^{2} \right)$$

$$\left( 1.16 \right) \mathrm{cov}\left[ \delta_{j},\delta_{k} | \mathcal{l>}t \right]=-\frac{\sigma_{PGS}^{2}}{R^{2}}q_{j}^{2}q_{k}^{2}i(i-t)$$

$$\left( 1.16 \right) \mathrm{cov}\left[ \delta_{j},\delta_{k} | \mathcal{l>}t \right]=-\frac{\sigma_{PGS}^{2}}{R^{2}}\left( \frac{{h_{j}\hat{\beta}}_{j}^{2}R^{2}}{\sigma_{PGS}^{2}} \right)\left( \frac{{h_{k}\hat{\beta}}_{k}^{2}R^{2}}{\sigma_{PGS}^{2}} \right)i(i-t)$$

$$\left( 1.16 \right) \mathrm{cov}\left[ \delta_{j},\delta_{k} | \mathcal{l>}t \right]=\boldsymbol{-}\left( \frac{\boldsymbol{R}^{\boldsymbol{2}}}{\boldsymbol{\sigma}_{\boldsymbol{PGS}}^{\boldsymbol{2}}} \right)\boldsymbol{h}_{\boldsymbol{j}}\boldsymbol{h}_{\boldsymbol{k}}{\hat{\boldsymbol{\beta}}}_{\boldsymbol{j}}^{\boldsymbol{2}}{\hat{\boldsymbol{\beta}}}_{\boldsymbol{k}}^{\boldsymbol{2}}\boldsymbol{i(i-t)}$$

**Appendix A** also verifies that combining Eqn. (1.13) and Eqn. (1.16) leads to Eqn. (1.4).

##### 1.4. Target parameter and identifiability

From Eqn. (1.3) $E\left[ \delta| \mathcal{l>}t \right]=\boldsymbol{iR}\boldsymbol{\sigma}_{\boldsymbol{PGS}}$**,** where $i=\phi(t)/[1-\Phi(t)]$ with $\phi$ and $\Phi$ denoting the probability density function and cumulative distribution function of a standard Gaussian distribution, respectively. In practice, $R$ is not separately identifiable because different combinations of $R$ an $t$ (and therefore, $i$) can yield the same $E\left[ \delta| \mathcal{l>}t \right]$. We therefore focus on a new estimator:

$$\left( 1.17 \right) \theta=\frac{E\left[ \delta| \mathcal{l>}t \right]}{\sigma\left[ g \right]}=\frac{E\left[ \delta| \mathcal{l>}t \right]}{\boldsymbol{\sigma}_{\boldsymbol{PGS}}}\boldsymbol{=iR}$$

Given that $i$ is finite and strictly positive, $\theta$ significantly different from 0 implies that $R$ must also be distinct from 0. Moreover, both $\theta$ and $R$ have the same sign because $i>0$, such that knowing the sign $\theta$ informs the direction of ascertainment. Thus, $R$ itself requires an external information on $t$ (or $i$) to be recovered

##### 1.5. Bias in an association analysis between outcome and polygenic score in an ascertained sample

We further extend the liability model by adding an observed exposure $\varphi$. Selection still acts on the liability ($\mathcal{l}$), so conditioning on $\mathcal{l>}t$ will create a spurious association between $g$ and $Y$. Consider the model,

$\left( 1.22 \right) \varphi=ag+e_{Y}$,	$e_{Y}\sim N\left( 0, 1-a^{2} \right)$

$\left( 1.23 \right)\mathcal{l=}Rg+\gamma\varphi+\varepsilon_{\mathcal{l}}$,	$\varepsilon_{\mathcal{l}}\sim N\left( 0, 1-R^{2}-\gamma^{2} \right)$

where $a$ is the genetic effect on $g$ on $Y$ and $\gamma$ is the causal effect of $g$ on $\mathcal{l}$. The residual $e_{Y}$ is independent of $g$ and $\varepsilon_{\mathcal{l}}$ is independent of $g$ and $e_{Y}$. In the non-ascertained population, $\mathrm{cov}\left( g,\varphi\right)=a$, $\mathrm{cov}\left( g\mathcal{, l} \right)=R^{2}+\gamma a$, and $\mathrm{cov}\left( \varphi\mathcal{, l} \right)=aR+\gamma$

$\left( 1.24 \right) E\left[ g\mathcal{|l>}t \right]= \left( R+ \gamma a \right)i,$	$E\left[ \varphi\mathcal{|l>}t \right]= \left( aR+ \gamma\right)i,$

$$\left( 1.25 \right) \mathrm{var}\left[ g\mathcal{|l>}t \right]= 1-i\left( i-t \right)\left( R+ \gamma a \right)^{2}$$

$$\left( 1.26 \right) \mathrm{var}\left[ \varphi\mathcal{|l>}t \right]= 1-i\left( i-t \right)\left( aR+ \gamma\right)^{2}$$

$$\left( 1.27 \right) \mathrm{cov}\left[ g,\varphi\mathcal{|l>}t \right]= a-\left( R+ \gamma a \right)\left( aR+ \gamma\right)i\left( i-t \right)$$

The genetic effect of g on $\varphi$ among the ascertained individual becomes,

$$\left( 1.28 \right) \hat{a}=\frac{\mathrm{cov}\left[ g,\varphi\mathcal{|l>}t \right]}{\mathrm{var}\left[ g\mathcal{|l>}t \right]}=\frac{a-\left( R+ \gamma a \right)\left( aR+ \gamma\right)i\left( i-t \right)}{1-i\left( i-t \right)\left( R+ \gamma a \right)^{2}}$$

The bias term relative to the true effect a

$$\left( 1.29 \right) \mathrm{bias}=\frac{a-\left( R+ \gamma a \right)\left( aR+ \gamma\right)i\left( i-t \right)}{1-i\left( i-t \right)\left( R+ \gamma a \right)^{2}}-\frac{a\left[ 1-i\left( i-t \right)\left( aR+ \gamma\right)^{2} \right]}{1-i\left( i-t \right)\left( R+ \gamma a \right)^{2}}$$

$$= -\frac{\gamma\left( 1-a^{2} \right)\left[ R+ \gamma a \right]i\left( i-t \right)}{1-i\left( i-t \right)\left( R+ \gamma a \right)^{2}}$$

##### **Appendix A**

***Calculation of*** $\mathbf{E}\left[ \boldsymbol{e}_{\boldsymbol{j}}^{\boldsymbol{'}}\boldsymbol{e}_{\boldsymbol{k}}^{\boldsymbol{'}} \right]$

Eqn. (1.9) implies that

$$\left( A.1 \right) E\left[ e_{j}^{'}e_{k}^{'} \right]=E\left[ \left[ \mathcal{l}_{-j}-\left( 1-q_{j}^{2} \right)\mathcal{l} \right]\left[ \mathcal{l}_{-k}-\left( 1-q_{k}^{2} \right)\mathcal{l} \right] \right]$$

$$\left( 1.14 \right) E\left[ e_{j}^{'}e_{k}^{'} \right]=E\left[ \mathcal{l}_{-j}\mathcal{l}_{-k}-\left( 1-q_{j}^{2} \right)\mathcal{l}\mathcal{l}_{-k}-\left( 1-q_{k}^{2} \right)\mathcal{l}\mathcal{l}_{-j}+\left( 1-q_{j}^{2} \right)\left( 1-q_{l}^{2} \right)\mathcal{l}^{2} \right]$$

$$\left( 1.14 \right) E\left[ e_{j}^{'}e_{k}^{'} \right]=E\left[ \mathcal{l}_{-j}\mathcal{l}_{-k}]-\left( 1-q_{j}^{2} \right)E[\mathcal{l}\mathcal{l}_{-k}]-\left( 1-q_{k}^{2} \right)E[\mathcal{l}\mathcal{l}_{-j}]+\left( 1-q_{j}^{2} \right)\left( 1-q_{l}^{2} \right)E[\mathcal{l}^{2} \right]$$

$$\left( 1.14 \right) E\left[ e_{j}^{'}e_{k}^{'} \right]=E[\mathcal{l}_{-j}\mathcal{l}_{-k}]-\left( 1-q_{j}^{2} \right)cov[\mathcal{l,}\mathcal{l}_{-k}]-\left( 1-q_{k}^{2} \right)cov[\mathcal{l,}\mathcal{l}_{-j}]+\left( 1-q_{j}^{2} \right)\left( 1-q_{l}^{2} \right)var[\mathcal{l]}$$

$$\left( 1.14 \right) E\left[ e_{j}^{'}e_{k}^{'} \right]=E[\mathcal{l}_{-j}\mathcal{l}_{-k}]-\left( 1-q_{j}^{2} \right)\left( 1-q_{k}^{2} \right)-\left( 1-q_{k}^{2} \right)\left( 1-q_{j}^{2} \right)+\left( 1-q_{j}^{2} \right)\left( 1-q_{l}^{2} \right)$$

$$\left( 1.14 \right) E\left[ e_{j}^{'}e_{k}^{'} \right]=E[\mathcal{l}_{-j}\mathcal{l}_{-k}]-\left( 1-q_{j}^{2} \right)\left( 1-q_{k}^{2} \right)$$

$$\left( 1.14 \right) E\left[ e_{j}^{'}e_{k}^{'} \right]=E[\left( \mathcal{l-}\frac{R\hat{\beta}_{j}(X_{j}-2p_{j})}{\sigma_{PGS}} \right)\left( \mathcal{l-}\frac{R\hat{\beta}_{k}(X_{k}-2p_{k})}{\sigma_{PGS}} \right)]-\left( 1-q_{j}^{2}-q_{k}^{2}+q_{j}^{2}q_{k}^{2} \right)$$

$$\left( 1.14 \right) E\left[ e_{j}^{'}e_{k}^{'} \right]=\left( 1-\frac{R^{2}\hat{\beta}_{j}^{2}h_{j}}{\sigma_{PGS}^{2}}-\frac{R^{2}\hat{\beta}_{k}^{2}h_{k}}{\sigma_{PGS}^{2}} \right)-\left( 1-q_{j}^{2}-q_{k}^{2}+q_{j}^{2}q_{k}^{2} \right)$$

$$\left( 1.14 \right) E\left[ e_{j}^{'}e_{k}^{'} \right]=(1-q_{j}^{2}-q_{j}^{2})-\left( 1-q_{j}^{2}-q_{k}^{2}+q_{j}^{2}q_{k}^{2} \right)$$

$$\left( 1.14 \right) E\left[ e_{j}^{'}e_{k}^{'} \right]= \mathbf{-}\boldsymbol{q}_{\boldsymbol{j}}^{\boldsymbol{2}}\boldsymbol{q}_{\boldsymbol{k}}^{\boldsymbol{2}}$$

***Calculation of*** $\mathbf{var}\left[ \boldsymbol{\delta} | \mathcal{l>}\boldsymbol{t} \right]$ ***from Eqn. (1.13) and Eqn. (1.16)***

We have that $\delta=\sum_{j=1}^{M} \delta_{j} ,$ which implies that

$$\left( A.2 \right) \mathrm{var}\left[ \delta| \mathcal{l>}t \right]=\sum_{j=1}^{M} \mathrm{var}\left[ \delta_{j} | \mathcal{l>}t \right]+\sum_{j\neq k} \mathrm{cov}\left[ \delta_{j},\delta_{k} | \mathcal{l>}t \right]$$

$$\left( A.3 \right) \mathrm{var}\left[ \delta| \mathcal{l>}t \right]=\sum_{j=1}^{M} {h_{j}\hat{\beta}}_{j}^{2}\left[ 1-q_{j}^{2}i\left( i-t \right) \right]-\frac{R^{2}}{\sigma_{PGS}^{2}}i(i-t)\sum_{j\neq k} h_{j}h_{k}\hat{\beta}_{j}^{2}\hat{\beta}_{k}^{2}$$

$$\left( A.3 \right) \mathrm{var}\left[ \delta| \mathcal{l>}t \right]=\sum_{j=1}^{M} {h_{j}\hat{\beta}}_{j}^{2}-i\left( i-t \right)\sum_{j=1}^{M} {h_{j}\hat{\beta}}_{j}^{2}q_{j}^{2}-\frac{R^{2}}{\sigma_{PGS}^{2}}i(i-t)\sum_{j\neq k} h_{j}h_{k}\hat{\beta}_{j}^{2}\hat{\beta}_{k}^{2}$$

$$\left( A.3 \right) \mathrm{var}\left[ \delta| \mathcal{l>}t \right]=\sigma_{PGS}^{2}-i\left( i-t \right)\sum_{j=1}^{M} {h_{j}\hat{\beta}}_{j}^{2}\left( \frac{R^{2}\hat{\beta}_{j}^{2}h_{j}}{\sigma_{PGS}^{2}} \right)-\frac{R^{2}}{\sigma_{PGS}^{2}}i\left( i-t \right)\left[ \left( \sum_{j=1}^{M} h_{j}\hat{\beta}_{j}^{2} \right)^{2}-\sum_{j=1}^{M} h_{j}^{2}\hat{\beta}_{j}^{4} \right]$$

$$\left( A.3 \right) \mathrm{var}\left[ \delta| \mathcal{l>}t \right]=\sigma_{PGS}^{2}-i\left( i-t \right)\frac{R^{2}}{\sigma_{PGS}^{2}}\sum_{j=1}^{M} h_{j}^{2}\hat{\beta}_{j}^{4}-\frac{R^{2}}{\sigma_{PGS}^{2}}i\left( i-t \right)\left[ \left( \sum_{j=1}^{M} h_{j}\hat{\beta}_{j}^{2} \right)^{2}-\sum_{j=1}^{M} h_{j}^{2}\hat{\beta}_{j}^{4} \right]$$

$$\left( A.3 \right) \mathrm{var}\left[ \delta| \mathcal{l>}t \right]=\sigma_{PGS}^{2}-i\left( i-t \right)\frac{R^{2}}{\sigma_{PGS}^{2}}\sum_{j=1}^{M} h_{j}^{2}\hat{\beta}_{j}^{4}-\frac{R^{2}}{\sigma_{PGS}^{2}}i\left( i-t \right)\left[ \left( \sum_{j=1}^{M} h_{j}\hat{\beta}_{j}^{2} \right)^{2}-\sum_{j=1}^{M} h_{j}^{2}\hat{\beta}_{j}^{4} \right]$$

$$\left( A.3 \right) \mathrm{var}\left[ \delta| \mathcal{l>}t \right]=\sigma_{PGS}^{2}-\frac{R^{2}}{\sigma_{PGS}^{2}}i\left( i-t \right)\sigma_{PGS}^{4}$$

$\left( A.3 \right) \mathrm{var}\left[ \delta| \mathcal{l>}t \right]=\boldsymbol{\sigma}_{\boldsymbol{PGS}}^{\boldsymbol{2}}\left[ \boldsymbol{1-}\boldsymbol{i}\left( \boldsymbol{i-t} \right)\boldsymbol{R}^{\boldsymbol{2}} \right]$**.**

***Derivation of variance of*** ${\hat{\boldsymbol{\theta}}}_{\boldsymbol{1}}$

The estimator $\hat{\theta}_{1}$ can be expressed as $\hat{\theta}_{1}=u/\sqrt{v}$, with $u=\sum2\hat{\beta}_{j}\left( \hat{p}_{j}^{(s)}-\hat{p}_{j}^{(r)} \right)$ and $v=\sum2\hat{\beta}_{j}^{2}\hat{p}_{j}^{(r)}\left( 1-\hat{p}_{j}^{(r)} \right)$. Now consider the function $f$ of $u$ and $v$ such that $f\left( u,v \right)=uv^{{-1}/2}$. The partial derivatives of $f$ with respect to $u$ and $v$ can be expressed below as

$$\left( A.3 \right) \frac{\partial f}{\partial u}=v^{{-1}/2} \mathrm{and} \frac{\partial f}{\partial v}={-\frac{1}{2}uv}^{{-3}/2}$$

Therefore, using the delta method, we can be expressed the sampling variance of $\hat{\theta}_{1}$ as

$$\left( A.4 \right) \mathrm{var}\left[ \hat{\theta}_{1} \right]\boldsymbol{\approx}\left( \frac{1}{\sqrt{v}} \right)^{\boldsymbol{2}}\mathrm{var}\left( u \right)+\left( {-\frac{1}{2}uv}^{{-3}/2} \right)^{2}\mathrm{var}\left( v \right)+2\left( \frac{1}{\sqrt{v}} \right)\left( {-\frac{1}{2}uv}^{{-3}/2} \right)\mathrm{cov}\left( u,v \right)$$

$$\left( A.4 \right) \mathrm{var}\left[ \hat{\theta}_{1} \right]\boldsymbol{\approx}\frac{\mathrm{var}\left( u \right)}{v}+\frac{u^{2}}{4v^{3}}\mathrm{var}\left( v \right)-\frac{u}{v^{2}}\mathrm{cov}\left( u,v \right)$$

$$\left( A.4 \right) \mathrm{var}\left[ \hat{\theta}_{1} \right]\boldsymbol{\approx}\left[ \frac{1}{v} \right]\left[ \mathrm{var}\left( u \right)+\frac{u^{2}}{4v^{2}}\mathrm{var}\left( v \right)-\frac{u}{v}\mathrm{cov}\left( u,v \right) \right]$$

We now derive below expressions of $\mathrm{var}\left( u \right)$, $\mathrm{var}\left( v \right)$ and $\mathrm{cov}\left( u,v \right)$ as a function of the estimated allele frequency $\hat{p}_{j}$ and its corresponding sampling variance.

$$\mathrm{var}\left( u \right)=\sum{4\hat{\beta}}_{j}^{2}\left[ \frac{\hat{p}_{s}\left( 1-\hat{p}_{s} \right)}{2N_{s}}+\frac{\hat{p}_{r}\left( 1-\hat{p}_{r} \right)}{2N_{r}} \right]$$

$$\mathrm{var}\left( v \right)=\sum{4\hat{\beta}}_{j}^{4}\mathrm{var}\left[ \hat{p}_{j}-\hat{p}_{j}^{2} \right]=\sum{4\hat{\beta}}_{j}^{4}\left\{ \mathrm{var}\left( \hat{p}_{j} \right)+\mathrm{var}\left( \hat{p}_{j}^{2} \right)-2\mathrm{cov}\left( \hat{p}_{j},\hat{p}_{j}^{2} \right) \right\}$$

Note that

$$\mathrm{var}\left( \hat{p}_{j}^{2} \right)=E\left[ \hat{p}_{j}^{4} \right]-E\left[ \hat{p}_{j}^{2} \right]^{2}=E\left[ \hat{p}_{j}^{4} \right]-\left[ \mathrm{var}\left( \hat{p}_{j} \right)+{E\left[ \hat{p}_{j} \right]}^{2} \right]^{2}$$

Assuming that $\hat{p}_{j}$ is normally distributed, then the non-central moment of $\hat{p}_{j}^{n}$ is:

$E\left[ \hat{p}_{j}^{2} \right]=\hat{p}_{j}^{2}+\mathrm{var}\left( \hat{p}_{j} \right)$,

$E\left[ \hat{p}_{j}^{3} \right]=\hat{p}_{j}^{3}+3p_{j}\mathrm{var}\left( \hat{p}_{j} \right)$,

$$E\left[ \hat{p}_{j}^{4} \right]=\hat{p}+6\hat{p}_{j}^{2}\mathrm{var}\left( \hat{p}_{j} \right)+3{\mathrm{var}\left( \hat{p}_{j} \right)}^{2}$$

Thus,

$$\mathrm{var}\left( \hat{p}_{j}^{2} \right)=\hat{p}_{j}^{4}+6\hat{p}_{j}^{2}\mathrm{var}\left( \hat{p}_{j} \right)+3{\mathrm{var}\left( \hat{p}_{j} \right)}^{2}-\left[ \mathrm{var}\left( \hat{p}_{j} \right)+{E\left[ \hat{p}_{j} \right]}^{2} \right]^{2}$$

$$\mathrm{var}\left( \hat{p}_{j}^{2} \right)=4\hat{p}_{j}^{2}\mathrm{var}\left( p_{j} \right)+2{\mathrm{var}\left( \hat{p}_{j} \right)}^{2}$$

$$\mathrm{cov}\left( \hat{p}_{j},\hat{p}_{j}^{2} \right)=E\left[ \hat{p}_{j}^{3} \right]- E\left[ \hat{p}_{j}^{2} \right]E\left[ \hat{p}_{j} \right]=2\hat{p}_{j}\mathrm{var}\left( \hat{p}_{j} \right)$$

Therefore,

$$\left( A.5 \right) \mathrm{var}\left( v \right)=\sum{4\hat{\beta}}_{j}^{4}\left\{ \mathrm{var}\left( \hat{p}_{j} \right) \left[ 1-2\hat{p}_{j} \right]^{2}+2{\mathrm{var}\left( \hat{p}_{j} \right)}^{2} \right\}$$

Finally,

$$\mathrm{cov}\left( u,v \right)=c\mathrm{ov}\left( {2\hat{\beta}}_{j}\hat{p}_{j}^{(s)}-{2\hat{\beta}}_{j}\hat{p}_{j}^{(r)},{2\hat{\beta}}_{j}^{2}\hat{p}_{j}^{(r)}\left( 1-\hat{p}_{j}^{(r)} \right) \right)$$

$$\mathrm{cov}\left( u,v \right)=\sum-{4\hat{\beta}}_{j}^{3}\mathrm{cov}\left( \hat{p}_{j}, \hat{p}_{j}-\hat{p}_{j}^{2} \right)$$

$$\mathrm{cov}\left( u,v \right)=\sum-4\hat{\beta}_{j}^{3}\left[ \mathrm{var}\left( \hat{p}_{j} \right)-\mathrm{cov}\left( \hat{p}_{j},\hat{p}_{j}^{2} \right) \right]$$

Since $\mathrm{cov}\left( \hat{p}_{j},\hat{p}_{j}^{2} \right)=2\hat{p}_{j} \mathrm{var}\left( \hat{p}_{j} \right)$, then it follows that

$\left( A.6 \right) \mathrm{cov}\left( u,v \right)=\sum-{4\hat{\beta}}_{j}^{3}\left( 1-2\hat{p}_{j} \right)\mathrm{var}\left( \hat{p}_{j} \right)$.

#### **Supplementary Notes 2: Ancestry deconvolution to control for confounding due to population stratification**

Our approach to detect ascertainment focuses on comparing the mean PGS in the sample being analysed (hereafter referred to as target sample) with that of a reference population

$$\left( 2.1 \right) \delta= \sum_{j=1}^{M} {2\hat{\beta}}_{j}(p_{j}^{\left( s \right)}-p_{j}^{\left( r \right)})$$

where $p_{j}^{\left( s \right)}$ and $p_{j}^{\left( r \right)}$ are the allele frequencies of SNP $j$ in the sample and reference, respectively, and $\hat{\beta}_{j}$ is the estimated effect size of SNP $j$ on the focal trait.

In practice, we use multiple reference samples to account for population stratification within the target sample. Specifically, we used an ancestry deconvolution approach previously proposed by Arriaga-MacKenzie, Matesi (1) and Privé (2). This approach express allele frequencies in the target sample as a convex linear combination of allele frequencies from multiple populations:

$$\left( 2.2 \right) p_{j}^{\left( s \right)}=\sum_{k=1}^{K} \pi_{k}^{\left( s \right)}p_{j}^{\left( r_{k} \right)}$$

Note that the weights $\pi_{k}^{\left( s \right)}$ are sample-dependent but not SNP- nor trait-dependent. Therefore, we can estimate them for any target sample from a large set of independent background SNPs not necessarily associated with any trait. We denote $q_{i}^{\left( s \right)}$ and $q_{i}^{\left( r_{k} \right)}$ as the estimated allele frequency of the $i$-th background SNP in the target and $k$-th reference sample, respectively. Estimates of $\pi_{k}^{\left( s \right)}$ (denoted below as $\hat{\pi}_{k}^{\left( s \right)}$) are obtained by finding a non-negative solution to the following minimization problem

$$\left( 2.3 \right) {\hat{\boldsymbol{\pi}}}^{(s)}=\left( \hat{\pi}_{1}^{\left( s \right)},\ldots, \hat{\pi}_{K}^{\left( s \right)} \right)=\underset{\pi_{k}^{\left( s \right)}\geq0}{\mathrm{argmin}} \sum_{i=1}^{L} \left( q_{i}^{\left( s \right)}-\sum_{k=1}^{K} \pi_{k}^{\left( s \right)}q_{i}^{\left( r_{k} \right)} \right)^{2}$$

where $L$ is the number of randomly sampled background SNPs.

In our analyses we set $L=100,000$ and solve (2.3) using the R package nnls. Once ${\hat{\boldsymbol{\pi}}}^{(s)}$ is obtained, we recalculate reference allele frequencies at trait-associated alleles as

$$\left( 2.4 \right) \hat{p}_{j}^{\left( r \right)}=\sum_{k=1}^{K} \hat{\pi}_{k}^{\left( s \right)}p_{j}^{\left( r_{k} \right)}$$

and use them to estimates $\theta$. The sampling variance of $\hat{p}_{j}^{\left( r \right)}$ is calculated as

$$\left( 2.5 \right) \mathrm{var}\left( \hat{p}_{j}^{\left( r \right)} \right)=\sum_{k=1}^{K} \left[ \hat{\pi}_{k}^{\left( s \right)} \right]^{2}\frac{p_{j}^{\left( r_{k} \right)}\left[ 1-p_{j}^{\left( r_{k} \right)} \right]}{2N_{r_{k}}}$$

Where $N_{r_{k}}$ denotes the size of the $k$-th reference sample.

#### **Supplementary Note 3: Simulation to assess the robustness of** ${\hat{\boldsymbol{\theta}}}_{\boldsymbol{1}}$ **and** ${\hat{\boldsymbol{\theta}}}_{\boldsymbol{2}}$ **to missing reference and attenuation bias**

We ran additional simulations to address two practical questions following our findings with the deconvolution strategy: (1) how robust are the two estimators when ancestry deconvolution relies on a reference panel that lacks one of the ancestries present in the target sample? and (2) to what extent does sampling error in reference panel (i.e. predictor) bias the ancestral loadings and how does this attenuation bias propagate to $\hat{\theta}_{1}$ and $\hat{\theta}_{2}$?

Our first question was motivated by the possibility that reference panel may lack ancestries present in the target sample. Such incompleteness can induce bias that could scale with the missing ancestry fraction. To model this scenario, we assumed that the observed allele frequencies of the target sample is a mixture of two composite reference ancestries: $\hat{p}^{(s)}=\pi_{s}\hat{p}^{(r1)}+\left( 1-\pi_{s} \right)\hat{p}^{(r2)}$ with allele frequency differences determined by *F*_ST_ $\in\left\{ 0.001, 0.01 \right\}$. If both reference (i.e., $\hat{p}^{(r1)}$ and $\hat{p}^{(r2)}$) were available, ancestral loadings $\left( \hat{\pi}_{s} \right)$could be estimated from sets of independent background SNPs and the estimates can be used to reconstruct a weighted (synthetic) reference. However, to explicitly simulate a missing-reference scenario, we assume that only $\hat{p}^{(r1)}$ was available in the reference panel, thereby inducing a scenario with a missing reference. We set $\pi_{s}\in\left\{ 0.1, 0.3, 0.5, 0.7, 0.9 \right\}$ and re-estimated $\hat{\pi}_{s}$ by regressing the target sample allele frequencies on the available reference using an independent background SNPs set. Since only one reference is observed, the reconstructed reference reduced to $\tilde{p}^{(r)}=\hat{\pi}_{s}\hat{p}_{1}$. We extended this simulation to the same drift and extreme model conditions in previous experiments (**Methods**). Across both scenarios, missing a reference component biased$\hat{\theta}_{1}$ and inflates its type I error (**Supplementary Figure 8**). On the contrary, $\hat{\theta}_{2}$remained robust and unbiased to that misspecification (**Supplementary Figure 8**). Importantly, the intercept term $\hat{\theta}_{2}$ (denoted as $I_{2}$) effectively absorbed the bias that inflated $\hat{\theta}_{1}$, providing a diagnostic indicator for missing reference and allowing $\hat{\theta}_{2}$ to recover the true ascertainment signal. (**Supplementary Figure 9**).

For the second question, our intuition is that even when all ancestries are present and observed in the reference panel (i.e., $\hat{p}^{(r1)}$ and $\hat{p}^{(r2)}$), allele frequencies are still estimated from finite samples. This finite sampling introduces noise, which could attenuate $\hat{\pi}_{s}$ during deconvolution. Moreover, we have previously shown that the size of the reference panel is a major limiting factor in detecting ascertainment signals using our framework. This limitation is particularly relevant because most publicly available reference panels, such as the 1KGP, include ancestry groups represented by only ~100 individuals on average. Such limited sample sizes not only reduce the precision of $\hat{\pi}_{s}$ but can also propagate sampling noise into the downstream estimates of $\hat{\theta}_{1}$ and $\hat{\theta}_{2}$, potentially biasing the two estimators. To assess this impact, we again simulated the target samples $\left( \hat{p}^{(s)}=\pi_{s}\hat{p}^{(r1)}+\left( 1-\pi_{s} \right)\hat{p}^{(r2)} \right)$. Here, the reference panel consists of $\hat{p}^{(r1)}$ and $\hat{p}^{(r2)}$, and we introduce sampling noise into the second reference ($\hat{p}^{(r2)}$) ancestry by varying its sample size across three levels: 1000, 500, 100, while other parameters remained fixed (**Methods**). Next, we re-estimated $\hat{\pi}_{s}$ and reconstructed a new reference across the different levels. Finally, we quantified the resulting bias and type I error for $\hat{\theta}_{1}$ and $\hat{\theta}_{2}$ under the same drift and extreme model. Our simulations showed that sampling error in reference panel biases $\hat{\pi}_{s}$, which in turn misspecifies the reconstructed references allele frequencies, especially with small sample size and low *F*_ST_ (**Supplementary Figure 10-11**). The bias became more pronounced when the under-sampled ancestry contributed more strongly to the target composition, resulting in substantial inflation to the type I error rate for $\hat{\theta}_{1}$(**Supplementary Figure 10-11**). Applying an attenuation correction to the ancestral loadings correct for this bias, keeping the empirical type I error of $\hat{\theta}_{1}$close to 5% even when one reference ancestry was severely under-sampled (**Supplementary Figure 10-11**). In contrast, $\hat{\theta}_{2}$ remained robust to sampling error, maintaining well-calibrated type I error rates without any correction (**Supplementary Figure 10-11**).

#### **Supplementary Note 4: Sensitivity analyses using IPYSCH data**

First, we revisited the relationship between $\hat{\theta}_{1}$and $\hat{\theta}_{2}$ with real data. Using the same marginally associated SNPs ($P<5\times{10}^{-03}$) employed in the main analyses, we re-estimated our ascertainment signals using $\hat{\theta}_{1}$ across the 21 phenotypes and regressed $\hat{\theta}_{2}$ on $\hat{\theta}_{1}$ to quantify the systemic bias of the latter for each biobank. Consistent with our theoretical derivation and simulation results, $\hat{\theta}_{1}$and $\hat{\theta}_{2}$were almost perfectly linearly related across the 21 investigated traits in both iPSYCH cohort (**Supplementary Fig. 15**). Regression slopes were approximately equal to one, with near-zero intercepts, indicating that both estimators recovered essentially identical ascertainment signals when the reference population was appropriately specified. These results further support that differences between $\hat{\theta}_{1}$and $\hat{\theta}_{2}$ primarily arise under population stratification or reference misspecification, rather than from differences in the underlying quantity being estimated (**Supplementary Figure 8-11)**.

Second, we evaluated whether our findings were sensitive to SNP inclusion threshold. We repeated all analyses across increasingly stringent *P* value thresholds (from $5\times{10}^{-04}$ up to 5$\times{10}^{-08}$). For each threshold, we applied linkage disequilibrium (LD) clumping to ensure approximately independent SNPs with ancestry-specific reference LD panel based on the super-population from the 1KGP using PLINK (r² < 0.01, window size = 1 Mb). We observed strong consistency in the direction of ascertainment signalsacross thresholds (**Supplementary Fig. 16**), indicating that the inferred signal is not driven by specific subsets of SNPs. As expected, the magnitude of the point estimates generally decreased as the *P* value threshold became more stringent. This attenuation likely reflects the reduced number of SNPs and corresponding loss of polygenic signal captured at stricter thresholds, rather than changes in the underlying ascertainment pattern itself. Consistent with this interpretation, some traits no longer reached statistical significance at the most stringent thresholds (**Supplementary Fig. 17–18**), likely due to reduced predictive power rather than changes in the underlying expected value of $\theta.$

Lastly, we also evaluated the sensitivity of our estimators to the choice of GWAS effect sizes, which in this paper were often derived from studies predominantly composed of participants of European ancestry. To test this, we downloaded GWAS summary statistics from the non-European ancestry, specifically Japanese and Taiwanese population for four representative traits, namely height, SBP, DBP, and BMI. For each phenotype, we analysed sets of independently associated SNPs (*P*<$5\times{10}^{-03}$) using European-derived SNP sets as in the main analyses. Across all four traits and both study populations, estimates of $\hat{\theta}_{2}$ were broadly similar irrespective of whether discovery effect sizes were derived from European or East Asian GWAS. Although point estimates were somewhat attenuated when using Japanese or Taiwanese GWAS effect sizes, this likely reflects the substantially smaller number of overlapping SNPs available after restricting analyses to the European-derived independent SNP sets (**Supplementary Fig. 20–21**).

#### Supplementary Note 5: Implication for within-biobank analyses

Next, we asked whether the ascertainment that we have detected can be translated into an a-priori prediction of the bias that will arise when an exposure $\left( \varphi\right)$ is regressed on the $g$ in an ascertained sample. We model this situation by treating the liability $\mathcal{l}$as a collider that is jointly influenced by both $g$ and $\varphi$: $\varphi=ag+e_{\varphi}$, and $\mathcal{l=}Rg+\gamma\varphi+\varepsilon_{\mathcal{l}}$, where $a$ is the true genetic effect of 𝑔 on $\varphi$, $\gamma$ is the direct selection effect of $\varphi$ and $\mathcal{l}$, while $e_{\varphi}$ and$\varepsilon_{\mathcal{l}}$ are independent residuals, respectively (full derivation in **Supplementary Note 1**). Conditioning on $\mathcal{l}>t$ (i.e., selecting study participants) opens the collider pathway on $g\to\mathcal{l\leftarrow}\varphi$ creating a spurious correlation between 𝑔 and $\varphi$ even when none exists in the non-ascertained population (**Supplementary Fig. 24**). This path diagram (**Supplementary Fig. 24**) yields a closed-form expression for the expected slope $\left( \hat{a} \right)$ of $\varphi$ on 𝑔 in the ascertained sample (see **Supplementary Note 1.4**). To validate our model, we ran a simulation for each combination of the true genetic effect $\left( a\in\left\{ 0, 0.1, 0.2, 0.3, \right\} \right)$ and *R*, with the direct selection effect fixed at $\gamma=0.1$. Theory predicts that the observed genetic effect in the ascertained sample will be attenuated and can even flip sign as the as the strength of ascertainment, *R*, increases. We showed through simulation that genetic effect in the ascertainment sample decreases monotonically with *R* and is well captured by prediction equation (**Supplementary Fig. 25**).

### **SUPPLEMENTARY METHODS**

***Cohort recruitment, genotyping, and QC***

**The Integrative Psychiatric Research (iPSYCH) cohort**

The iPSYCH study is a large population-based case–control cohort from Denmark designed to investigate the genetic basis of psychiatric disorders (3, 4). Participants were drawn from nationwide registers, with cases identified based on psychiatric diagnoses and controls sampled from the general population. Recruitment procedures, cohort characteristics, genotyping, and quality control have been described in detail previously (3-5). Genotype data were generated from two genotyping waves (iPSYCH2012 and iPSYCH2015) and processed using the Ricopili pipeline. Standard quality control procedures were applied, including removal of related individuals, duplicates, and ancestry outliers based on principal components. Analyses were restricted to individuals of European ancestry. Variants were filtered based on imputation quality (INFO > 0.1), minor allele frequency (MAF > 0.005), and missingness (< 2%). For the present study, we derived allele frequency estimates from three datasets:

- iPSYCH2012: cases and controls from the 2012 genotyping wave
- iPSYCH2015: cases and controls from the 2015 genotyping wave, excluding individuals present in the 2012 wave
- iPSYCH controls (combined): a combined set of control individuals from both genotyping waves

Allele frequencies were computed separately for each dataset using PLINK v2.0 on a per-chromosome basis and subsequently merged across chromosomes. After quality control, 4,347,148 SNPs remained for iPSYCH2012, 3,723,020 for iPSYCH2015, and 4,439,439 for the combined control dataset.

**The Australian Genomics Depression Study (AGDS)**

The AGDS is a large depression cohort with comprehensive genetic and phenotypic data, comprising 15,792 participants of European ancestry, 92% of whom met diagnostic criteria for major depressive disorder (MDD) (6). The recruitment procedures and cohort characteristics of the AGDS have been described in detail previously (6, 7). Briefly, over 21,000 participants were recruited nationwide through a dual strategy combining invitations via government prescription records and a media campaign. Participants completed an online questionnaire that included a core diagnostic module assessing Major Depressive Disorder (MDD) based on DSM-5 criteria using the Composite International Diagnostic Interview-Short Form, alongside modules capturing comorbid psychiatric and behavioural phenotypes. Individuals meeting DSM-5 criteria for lifetime MDD and without self-reported schizophrenia, bipolar disorder, or ADHD ($N_{s}$ = 13,104) were further classified into non-mutually exclusive subtypes including early/late-onset, recurrent, atypical, comorbid anxiety, and seasonal affective disorder. The AGDS participants were genotyped using the Illumina Global Screening Array v2. Quality control followed standard procedures using GenomeStudio and PLINK 1.9, excluding SNPs with GenTrain score < 0.6, MAF < 0.01, call rate < 95%, or Hardy–Weinberg equilibrium (HWE) deviation (*P* value < 1×10⁻⁶). Sample-level QC excluded individuals with call rate < 95%, sex discrepancies, or evidence of non-European ancestry based on principal components. After QC, 14,252 AGDS participants (13,104 MDD cases) of European ancestry remained. Next, imputation was performed using the Haplotype Reference Consortium (HRC v1.1) reference panel and only variants with high imputation R^2^ > 0.6 were retained (Mitchell et al., 2022). Allele frequencies for the AGDS cohort were directly derived from post-QC imputed data using PLINK v1.9 with standard filtering thresholds (MAF< 0.05, geno 0.05, and HWE *P* value < 1×10⁻⁶). After QC, only 4,157,970 variants remained and were used in all downstream analyses for the AGDS population.

**BioBank Japan (BBJ)**

The BBJ project is a nation-wide hospital-based prospective biobank established in 2003 to investigate the genetic basis of common diseases in the Japanese population (8). The recruitment procedures, cohort characteristics, genotyping process and QC of the BBJ have been described in detail previously (8, 9). Allele frequencies for BBJ were obtained directly from the publicly released post-QC imputed dataset (8), Accession code: hum0197; <https://humandbs.biosciencedbc.jp/files/hum0197/hum0197.v3.BBJ.Hei.v1.zip>. Height was selected as the focal phenotype to represent BBJ, as it is uniformly available, and heritable, allowing robust estimation of cohort-specific allele frequency patterns. The summary statistics do not report per-variant sample sizes; therefore, we used the total post-QC sample size reported in Sakaue, Kanai (8), corresponding to $N_{s}$= 178,726 individuals. After removing SNPs with MAF<5%, only 3,808,287 variants remained and were used in all downstream analyses for the BBJ population.

**Estonian Biobank (EstBB)**

The EstBB is a population-based biobank established in 2000 to support genetic and epidemiological research in Estonia (10, 11). The recruitment procedures, cohort characteristics, genotyping process and QC have been described in detail previously (12). We derived allele frequencies for EstBB from the publicly available dataset (<https://ftp.ebi.ac.uk/pub/databases/gwas/summary_statistics/GCST90624001-GCST90625000/GCST90624701/GCST90624701.tsv.gz>). Body height was selected as the focal trait as it is uniformly available, and heritable, allowing robust estimation of cohort-specific allele frequency patterns. The summary statistics do not report per-variant sample sizes; therefore, we used the total post-QC sample size reported in Abner, Batool (12), corresponding to $N_{s}$= 204,747 individuals. After removing SNPs with MAF<5%, only 4,031,707 variants remained and were used in all downstream analyses for the EstBB population.

**The Mexican city prospective cohort study (MCPS)**

The MCPS is a large population-based cohort established to investigate chronic disease risk in adults residing in Mexico City (13). The recruitment procedures, cohort characteristics, and genotyping procedure have been described in detail previously (13). We obtained genome-wide allele frequencies for MCPS from the publicly available dataset (<https://ftp.ebi.ac.uk/pub/databases/gwas/summary_statistics/GCST90435001-GCST90436000/GCST90435342/> (13)). Note that the summary statistics files do not report per-variant sample sizes; therefore, we used the total post-QC sample size reported in Wen, Kuri-Morales (13), which is $N_{s}$= 136,401 individuals. Allele frequency estimates from these summary statistics were treated as the MCPS target population in all ascertainment analyses. After removing SNPs with MAF<5%, only 4,491,214 variants remained for all downstream analyses.

**Mexican biobank (MXB)**

The MXB project is based on the Encuesta Nacional de Salud 2000 (ENSA 2000), a nationally representative, probabilistic, multi-stage, stratified household survey conducted by the Mexican Secretariat of Health between November 1999 and June 2000 (14). The recruitment procedures, cohort characteristics, and genotyping procedure have been described in detail previously (14). In this study, allele frequencies were obtained directly from the publicly released post-QC imputed dataset (14), <https://doi.org/10.5281/zenodo.7420254>. Similar to BBJ, we used height as the focal trait for this biobank. The summary statistics report per-variant sample sizes; we therefore used the maximum reported sample size ($N_{s}$= 5,663) to represent the effective study size of the MXB cohort. After removing SNPs with MAF<5%, only 4,427,753 variants remained for all downstream analyses.

**The children cohort (age 8) from the Norwegian Mother and Child Cohort Study (MoBa_child8)**

The Norwegian Mother, Father and Child Cohort Study (MoBa) is a prospective, population-based pregnancy cohort that recruited pregnant women from 1999 to 2008 across 50 hospitals in Norway (15). Allele frequencies were derived from this QC-filtered subset of 14,747 children aged 8 (denoted as MoBa_child8; dataset available at: <https://www.fhi.no/en/ch/studies/moba/for-forskere-artikler/gwas-data-from-moba>, (15)). The summary statistics report per-variant sample sizes; we therefore used the maximum reported sample size ($N_{s}$= 3,862) to represent the effective study size of the MoBa_child8 cohort. After removing SNPs with MAF<5%, only 3,686, 529 variants remained for all downstream analyses.

**SweGen biobank**

The SweGen biobank represents a population-based whole-genome sequencing reference panel generated from Swedish national cohorts to capture genetic variation across the Swedish population (16). For the present study, allele frequencies were extracted directly from this curated QCed dataset from public domain ([https://swefreq.nbis.se](https://swefreq.nbis.se/); <https://doi.org/10.17044/NBIS/G000003>). This dataset contains whole-genome variant frequencies for 1,000 Swedish individuals generated within the SweGen project. After removing SNPs with MAF<5%, only 3,659,846 variants remained for all downstream analyses.

**Taiwan biobank (TWB)**

The TWB is a large, population-based cohort designed to characterize genetic and environmental determinants of health in the Taiwanese population (17). The recruitment procedures, cohort characteristics, and genotyping procedure have been described in detail previously (17). Allele frequencies were derived directly from this QC-filtered and ancestry-homogeneous TWB for height available in the public domain (<https://ftp.ebi.ac.uk/pub/databases/gwas/summary_statistics/GCST90278001-GCST90279000/GCST90278637/GCST90278637.tsv.gz>). The summary statistics report per-variant sample sizes; we therefore used the maximum reported sample size ($N_{s}$= 92,615) to represent the effective study size of the TWB cohort. After removing SNPs with MAF<5%, only 4,558,956 variants remained for all downstream analyses.

**The UK Biobank (UKB)**

The UKB (<http://www.ukbiobank.ac.uk/>) is a large, population-based prospective cohort that recruited over 500,000 individuals aged 40–69 years between 2006 and 2010 to study genetic, lifestyle, and environmental determinants of complex diseases (18). Participants were invited from National Health Service (NHS) patient registers within a 25-mile radius of 22 assessment centres across the United Kingdom, with approximately 9 million individuals contacted and a final response rate of ~5%. Baseline data collection included detailed questionnaires, physical measurements, and biological samples, with subsets of participants attending repeat assessment and imaging visits. Genome-wide genotyping was performed using either the UK BiLEVE Axiom array (~807,000 markers) or the UK Biobank Axiom array (~826,000 markers). For this study, we used the UK biobank imputed genotype data. These genotypes included ~16.6 million SNPs imputed to the Haplotype reference Consortium (19) imputation reference panel in 487,409 participants of the UKB. We restricted our analyses to individuals of predominant European ancestry, identified using principal component projection onto the 1KGP. After excluding participants with sex discordance, withdrawn consent, missing genotypes, MAF <0.05, missingness > 5%, about 6.5 million variant for 348,502 unrelated participants remained. Allele frequencies for UKB were derived from this QC-filtered, ancestry-homogeneous subset of participants. After QC, only 3,602,963 variants remained for all downstream analyses.

### **SUPPLEMENTARY TABLES**

| **S/N** | **Cohort** | **Study design** | **Sample type** | **Recruitment** | **Primary ascertainment axis** | **Expected ascertainment** |
| --- | --- | --- | --- | --- | --- | --- |
| 1 | AGDS | Case-enriched | MDD | Self-selection + prescription records | Depression | Strong enrichment for MDD |
| 2 | BBJ | Hospital-based | Disease | Clinical recruitment | Disease-specific | Enrichment for disease cases |
| 3 | EstBB | Population-based | General | National registry | General health | Mild selection bias |
| 4 | iPSYCH2012 | Case–control | Psychiatric | National registry-based | Psychiatric liability | Enrichment for psychiatric disorders |
| 5 | iPSYCH2015 | Case–control | Psychiatric | National registry-based | Psychiatric liability | Enrichment for psychiatric disorders |
| 6 | MCPS | Population-based | General | Prospective cohort | Cardiometabolic | Low ascertainment |
| 7 | MoBa_child8 | Birth cohort | Children | Pregnancy cohort | Early-life traits | Participation bias |
| 8 | MXB | Survey-based | General | National survey | General population | Low ascertainment |
| 9 | QGP | Population-based | General | National initiative | Regional ancestry | Population structure effects |
| 10 | SweGen | Reference panel | General | Sequencing panel | None | Minimal ascertainment |
| 11 | TWB | Population-based | General | National biobank | General health | Mild selection bias |
| 12 | UKB | Population-based | General | Volunteer-based | Socio-economic / health | Healthy and educated volunteer bias |

**Supplementary Table S1. Overview of cohort design and expected ascertainment patterns.** Summary of the 12 biobanks included in this study, describing study design, sample composition, recruitment strategy, and the primary axis of ascertainment. Expected ascertainment reflects the anticipated direction and source of enrichment based on cohort design (for example, disease-specific recruitment, volunteer participation, or population-based sampling). This information provides context for interpreting inferred ascertainment signals across cohorts.

**Supplementary Table S2. Number of SNPs analysed per trait using census representative reference in iPYSCH cohort. For each of the 21 traits, we report the number of SNPs included in the ascertainment analyses for the iPSYCH2012 and iPSYCH2015 cohorts using census-representative reference (control-derived allele frequencies). SNPs were selected based on association significance (P < 0.005) and minor allele frequency > 0.05 in each population. Differences in SNP counts between iPSYCH2012 and iPSYCH2015 cohort reflect variation in SNP availability and matching with the control reference sample.**

| S/N | Traits | Number of SNPs | |
| --- | --- | --- | --- |
|  |  | iPYSCH2012 | iPYSCH2015 |
| 1 | Height | 8361 | 6962 |
| 2 | Body mass index | 5134 | 4305 |
| 3 | Educational attainment | 5003 | 4176 |
| 4 | Attention deficit hyperactivity disorder | 2109 | 1765 |
| 5 | Anxiety | 1993 | 1673 |
| 6 | Bipolar disorder | 2672 | 2216 |
| 7 | Major depressive disorder | 3710 | 3152 |
| 8 | Migraine | 1852 | 1526 |
| 9 | Post-traumatic stress disorder | 2203 | 1859 |
| 10 | Age at menarche | 4371 | 3676 |
| 11 | Age at natural menopause | 2090 | 1724 |
| 12 | Diastolic blood pressure | 4185 | 3496 |
| 13 | Systolic blood pressure | 4379 | 3659 |
| 14 | Problematic alcohol use | 2992 | 2563 |
| 15 | Bone mineral density | 1254 | 1061 |
| 16 | Coronary artery disease | 2866 | 2426 |
| 17 | Type 2 diabetes | 5159 | 4311 |
| 18 | Schizophrenia | 3686 | 3044 |
| 19 | Asthma | 1557 | 1335 |
| 20 | Rheumatoid arthritis | 2030 | 1699 |
| 21 | Covid-19 | 1737 | 1468 |

**Supplementary Table S3. Number of independent SNPs analysed for each trait across within-ancestry biobank comparisons. SNP counts are summarised separately for European, American and East Asian based on the pairwise comparisons presented in Supplementary Figure 12a-e. For each ancestry group, the table reports the mean, minimum and maximum number of SNPs analysed, together with the number of pairwise comparisons. SNPs were selected based on association significance (P < 0.005), minor allele frequency > 0.05, and successful harmonisation with the reference dataset. Variation in SNP counts reflects differences in SNP availability and matching with the reference panel.**

| **S/N** | **Traits** | **No of SNPs per ancestry comparison** | | | | | | | | |
| --- | --- | --- | --- | --- | --- | --- | --- | --- | --- | --- |
|  |  | **European** | | | **Mexican** | | | **East-Asian** | | |
| **Traits** |  | **No of pairwise comparison = 21** | | | **No of pairwise comparison = 1** | | | **No of pairwise comparison = 1** | | |
|  |  | **Mean** | **Min** | **Max** | **Mean** | **Min** | **Max** | **Mean** | **Min** | **Max** |
| 1 | Height | 2981 | 2955 | 2983 | 2683 | 2683 | 2683 | 3831 | 3831 | 3831 |
| 2 | BMI | 1965 | 1947 | 1967 | 1861 | 1861 | 1861 | 2567 | 2567 | 2567 |
| 3 | EA | 1895 | 1877 | 1897 | 1799 | 1799 | 1799 | 2407 | 2407 | 2407 |
| 4 | ADHD | 756 | 750 | 757 | 758 | 758 | 758 | 947 | 947 | 947 |
| 5 | Anxiety | 795 | 790 | 795 | 786 | 786 | 786 | 950 | 950 | 950 |
| 6 | BIP | 922 | 915 | 923 | 887 | 887 | 887 | 1151 | 1151 | 1151 |
| 7 | MDD | 1417 | 1405 | 1418 | 1319 | 1319 | 1319 | 1714 | 1714 | 1714 |
| 8 | Migraine | 684 | 675 | 685 | 659 | 659 | 659 | 846 | 846 | 846 |
| 9 | PTSD | 810 | 804 | 810 | 765 | 765 | 765 | 959 | 959 | 959 |
| 10 | AAM | 1634 | 1618 | 1635 | 1521 | 1521 | 1521 | 2057 | 2057 | 2057 |
| 11 | ANM | 710 | 703 | 710 | 667 | 667 | 667 | 862 | 862 | 862 |
| 12 | DBP | 1562 | 1547 | 1563 | 1490 | 1490 | 1490 | 2028 | 2028 | 2028 |
| 13 | SBP | 1657 | 1638 | 1658 | 1605 | 1605 | 1605 | 2093 | 2093 | 2093 |
| 14 | PAU | 1092 | 1082 | 1093 | 972 | 972 | 972 | 1306 | 1306 | 1306 |
| 15 | BMD | 455 | 451 | 455 | 447 | 447 | 447 | 517 | 517 | 517 |
| 16 | CAD | 1080 | 1067 | 1081 | 966 | 966 | 966 | 1242 | 1242 | 1242 |
| 17 | T2D | 1866 | 1847 | 1867 | 1650 | 1650 | 1650 | 2159 | 2159 | 2159 |
| 18 | SCZ | 1353 | 1341 | 1354 | 1208 | 1208 | 1208 | 1605 | 1605 | 1605 |
| 19 | Asthma | 575 | 565 | 575 | 489 | 489 | 489 | 602 | 602 | 602 |
| 20 | RA | 737 | 731 | 738 | 649 | 649 | 649 | 807 | 807 | 807 |
| 21 | COVID-19 | 596 | 593 | 596 | 538 | 538 | 538 | 668 | 668 | 668 |

**Supplementary Table S4. Publicly available GWAS summary statistics analysed in this study.**
For each of the 21 traits, we report the source URL for downloading summary statistics (regression coefficients and P values) and the corresponding reference of the discovery GWAS

| **S/N** | **Traits** | **URL** | **Reference** |
| --- | --- | --- | --- |
| 1 | Height | <https://giant-consortium.web.broadinstitute.org/images/4/4e/GIANT_HEIGHT_YENGO_2022_GWAS_SUMMARY_STATS_ALL.gz> | (20) |
| 2 | Body mass index | <https://giant-consortium.web.broadinstitute.org/images/c/c8/Meta-analysis_Locke_et_al%2BUKBiobank_2018_UPDATED.txt.gz> | (21) |
| 3 | Educational attainment | <https://thessgac.com/> | (22) |
| 4 | Attention deficit hyperactivity disorder | <https://doi.org/10.6084/m9.figshare.22564390> | (23) |
| 5 | Anxiety | <https://zenodo.org/records/13135834/files/ANX_EUR.txt.gz?download=1> | (24) |
| 6 | Bipolar disorder | <https://doi.org/10.6084/m9.figshare.27216117> | (25) |
| 7 | Major depressive disorder | <https://doi.org/10.6084/m9.figshare.27061255> | (26) |
| 8 | Migraine | <https://doi.org/10.1038/s41588-021-00990-0> | (27) |
| 9 | Post-traumatic stress disorder | https://doi.org/10.6084/m9.figshare.26349322 | (28) |
| 10 | Age at menarche | <https://www.repository.cam.ac.uk/items/8c5f7afb-5fa2-45ea-b52d-4e643bc2a5b7> | (29) |
| 11 | Age at natural menopause | <https://www.reprogen.org/reprogen_ANM_201K_170621.txt.gz> | (30) |
| 12 | Diastolic blood pressure | <http://ftp.ebi.ac.uk/pub/databases/gwas/summary_statistics/GCST90310001-GCST90311000/GCST90310294> | (31) |
| 13 | Systolic blood pressure | <http://ftp.ebi.ac.uk/pub/databases/gwas/summary_statistics/GCST90310001-GCST90311000/GCST90310295> | (31) |
| 14 | Problematic alcohol use | https://medicine.yale.edu/lab/gelernter/stats/ | (32) |
| 15 | Bone mineral density | http://www.gefos.org/ | (33) |
| 16 | Coronary artery disease | https://cardiogramplusc4d.org/data-downloads/ | (34) |
| 17 | Type 2 diabetes | https://www.diagram-consortium.org/downloads.html | (35) |
| 18 | Schizophrenia | https://doi.org/10.6084/m9.figshare.19426775 | (36) |
| 19 | Asthma | https://zenodo.org/records/5513443 | (37) |
| 20 | Rheumatoid arthritis | http://ftp.ebi.ac.uk/pub/databases/gwas/summary_statistics/GCST90132001-GCST90133000/GCST90132222 | (38) |
| 21 | Covid-19 | <https://storage.googleapis.com/covid19-hg-public/freeze_7/results/20220403/main/sumstats/COVID19_HGI_A2_ALL_leave_23andme_20220403_GRCh37.tsv.gz> | (39) |

**Supplementary Table S5. Linear regression of** ${\hat{\boldsymbol{\theta}}}_{\boldsymbol{2}}$ **and** ${\hat{\boldsymbol{\theta}}}_{\boldsymbol{1}}$**.** Results are presented separately for within-ancestry and cross-ancestry analyses and include the regression intercept, slope, R², and number of comparisons to show their relationship.

| **Comparison set** | **N comparisons** | **Intercept** | **Slope** | **R^2^** |
| --- | --- | --- | --- | --- |
| Within ancestry | 529 | -0.0008476 | 1.005 | 0.997 |
| Cross ancestry | 736 | -0.019072 | 0.929 | 0.973 |

### **SUPPLEMENTARY FIGURES**

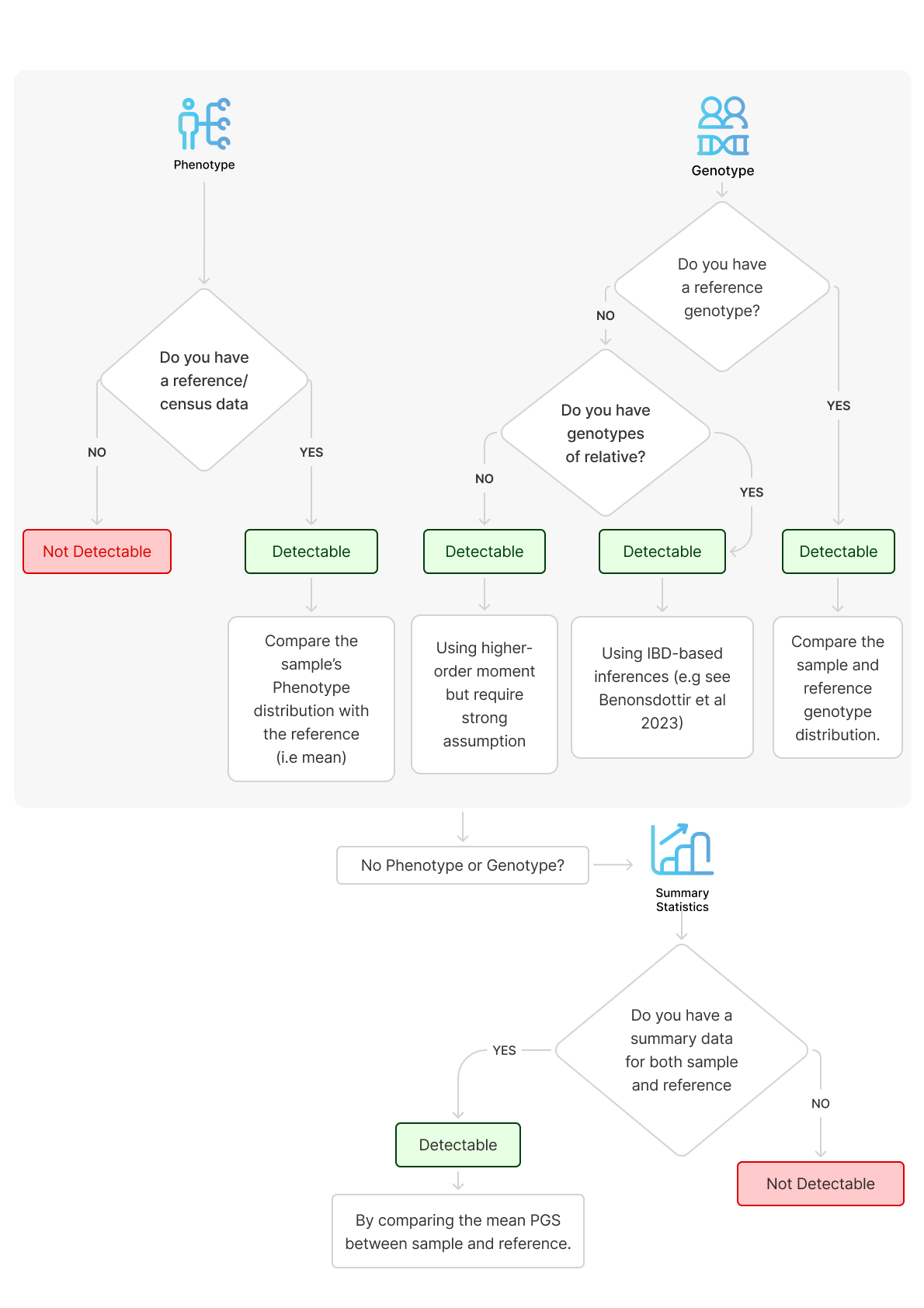

**Supplementary Fig. 1. Schematic overview of ascertainment bias detectability under different data availability scenarios.** Existing approaches can be broadly grouped into phenotype-based and genotype-based methods. When phenotype data are available for both the study sample and a representative reference population, ascertainment can be detected by comparing the distributions of the phenotype (e.g., mean). If individual-level genotypes are available, ascertainment can alternatively be inferred from genotype-based approaches, for example by comparing mean polygenic score between samples or leveraging identity-by-descent information. However, when neither phenotype nor genotype data are available at the individual level, ascertainment could be detected leveraging higher-order moment but requires strong assumption which in practice might not hold. In contrast, when only summary-level genotype data are available for both the target sample and a suitable reference population, ascertainment can still be assessed by comparing the mean polygenic score (PGS) between the two groups, an approach we formalize and evaluate in this study.

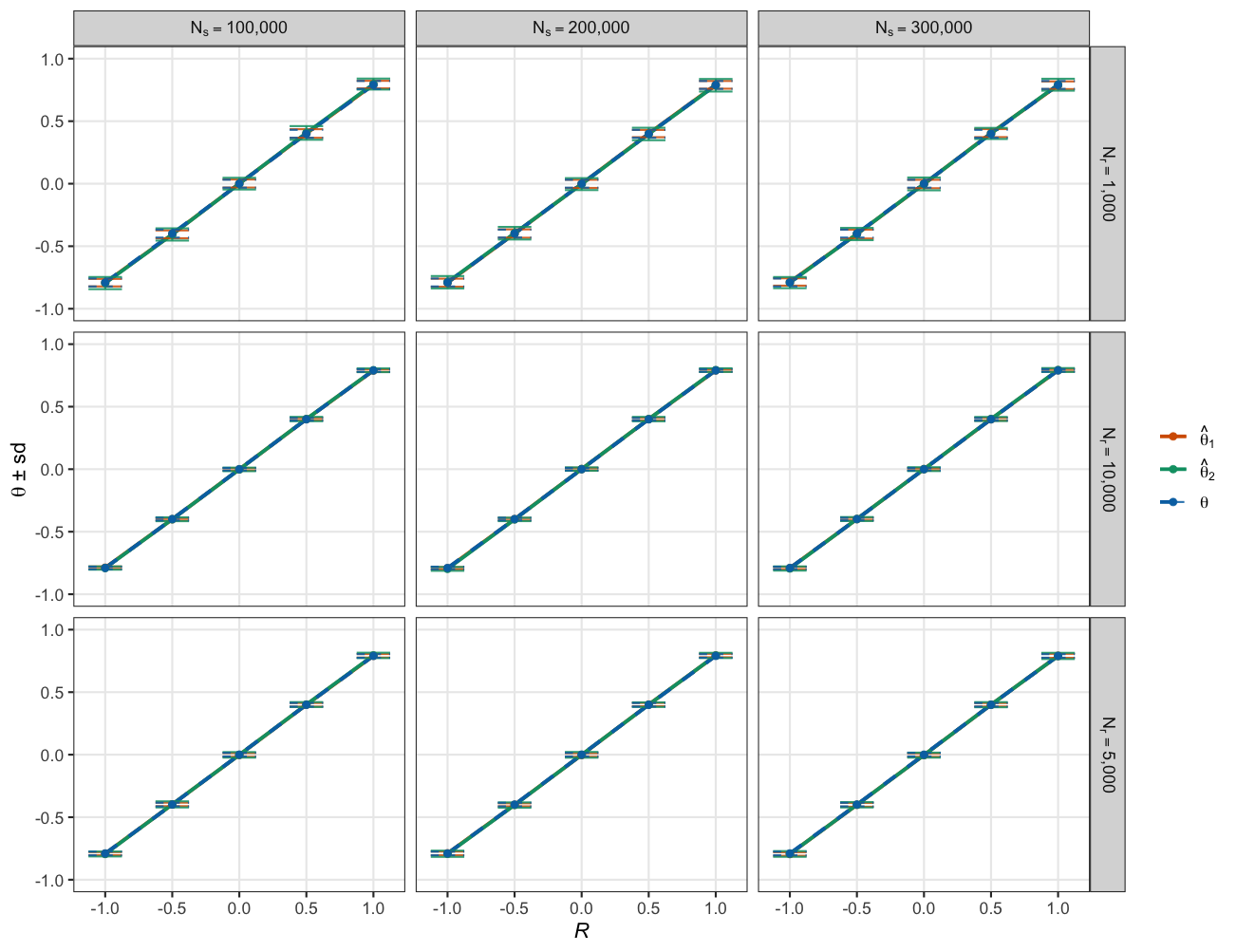

**Supplementary Fig. 2. Comparison of theoretical and empirical predictions of** $\boldsymbol{\theta}$ **across varying sample sizes and *R* (i.e., correlations between polygenic score and liability).** Each panel shows results for a specific combination of study sample size ($N_{s}$), and reference sample size ($N_{r}$). Empirical means and standard deviations of $\hat{\theta}_{1}$ (orange solid lines, ± s.d. error bars) and $\hat{\theta}_{2}$ (green solid lines, ± s.d. error bars) are compared with theoretical expectations ($\theta\pm s.d$; blue dash lines). Simulations were conducted across 1,000 replicates per parameter setting under a liability threshold model.

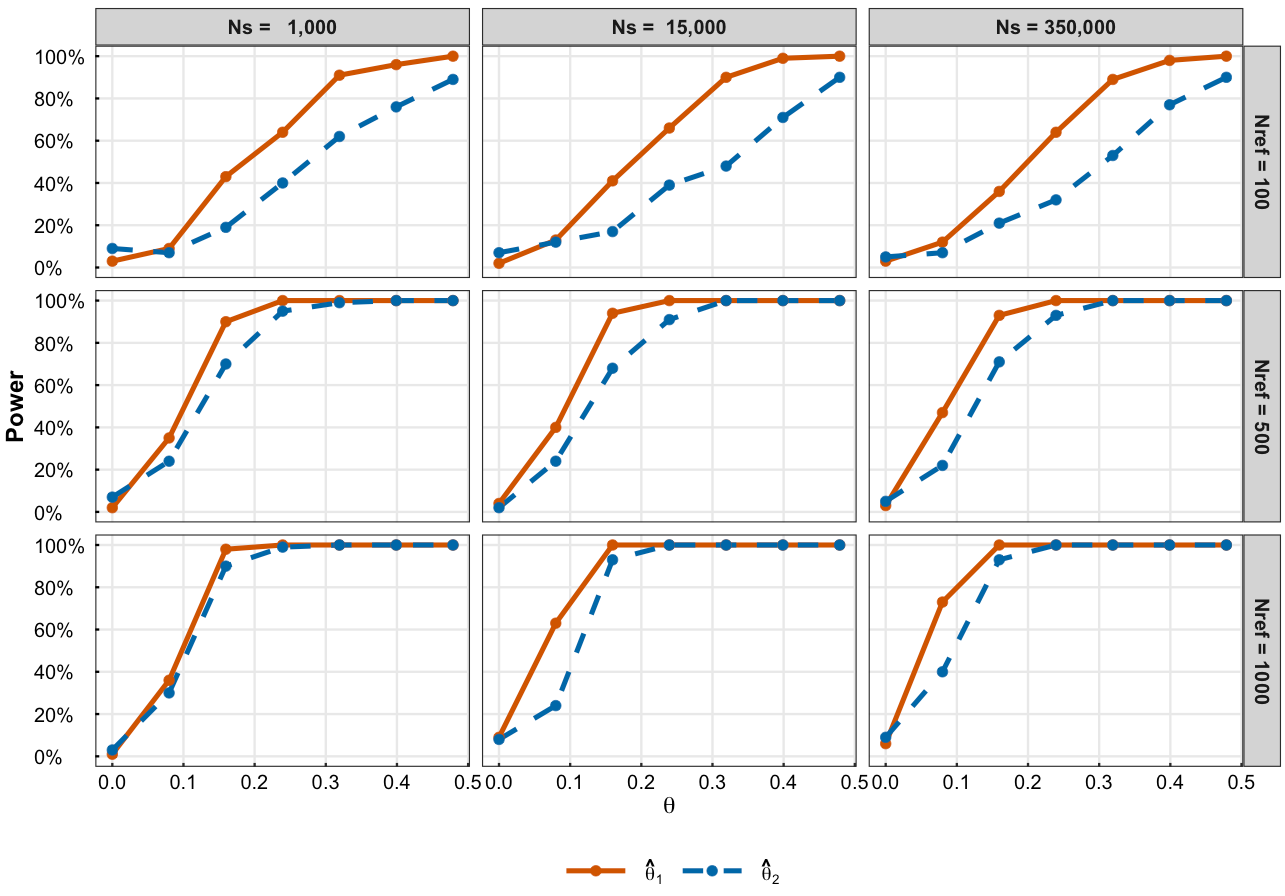

**Supplementary Fig. 3. Statistical power to detect ascertainment bias as a function of the ascertainment parameter** $\boldsymbol{\theta}$**, target sample size** $\boldsymbol{N}_{\boldsymbol{s}}$**, and reference sample size** $\boldsymbol{N}_{\boldsymbol{r}}$**.** Power was estimated from simulations under the liability threshold model for both $\hat{\theta}_{1}$(solid line) and $\hat{\theta}_{2}$(dashed line) at a significance level of $\alpha=0.05$. Power increased with the magnitude of $\theta$, but was strongly constrained by the size of the reference panel. Increasing the size of the target cohort did not improve power relative to increasing the size of the reference panel, indicating that **the precision of reference allele frequency estimates is the primary determinant of detectability.**

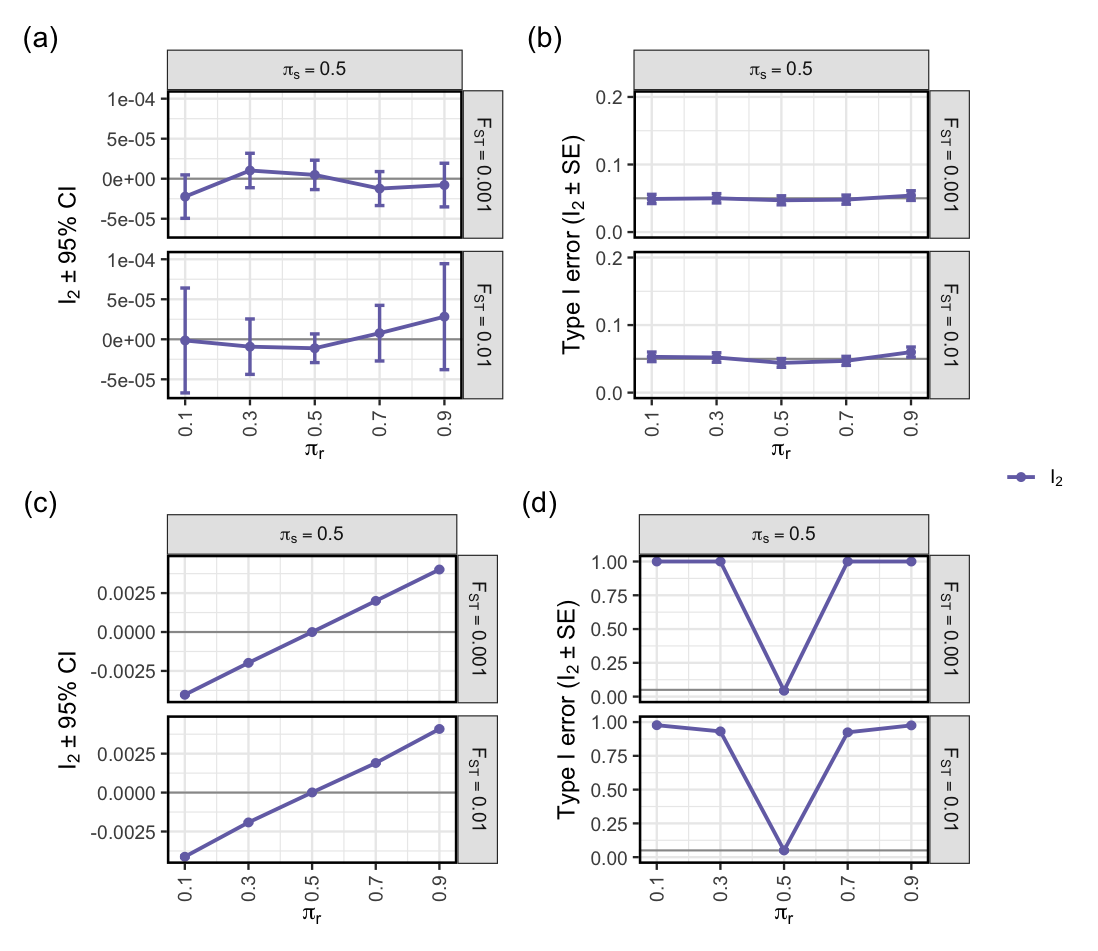

**Supplementary Fig. 4**. **Performance of** ${\hat{\boldsymbol{\theta}}}_{\boldsymbol{2}}$ **intercept (**$\boldsymbol{I}_{\boldsymbol{2}}$**) under drift (top row) and extreme (bottom row) model of population stratification across varying *F*_ST_.** Panels show results from simulations where both target and reference samples consist of two sub-populations with allele frequency differences determined by *F*_ST_ $\in\left\{ 0.001, 0.01 \right\}$ (**Methods**). The target sample was modelled with equal ancestral loadings ($\pi_{s}=0.5$), whereas the reference sample was simulated with either matched $\left( \pi_{r}=0.5 \right)$ or skewed ancestral loadings $\left( \pi_{r}\in\left\{ 0.1, 0.3, 0.7, 0.9 \right\} \right)$. **Panel (a, c)** shows the mean estimates of $I_{2}\pm95\% CI$ under drift (top) and extreme (bottom) model. **Panel (b, d)** shows the type I error rate of $I_{2}\pm SE$ under drift (top) and extreme (bottom) model. CI represent the confidence interval estimated as 1.96$\pm$s.e.m. Type I error was calculated as the fraction of *P* values below 0.05. SE represent standard error estimated as $\sqrt{\hat{p}\left( 1-p \right)}/n$, where $\hat{p}$ denotes the empirical type 1 error rate and $n$ represent the number of replicates. Simulation was performed across 10,000 replicates.

i) **Genetic drift**

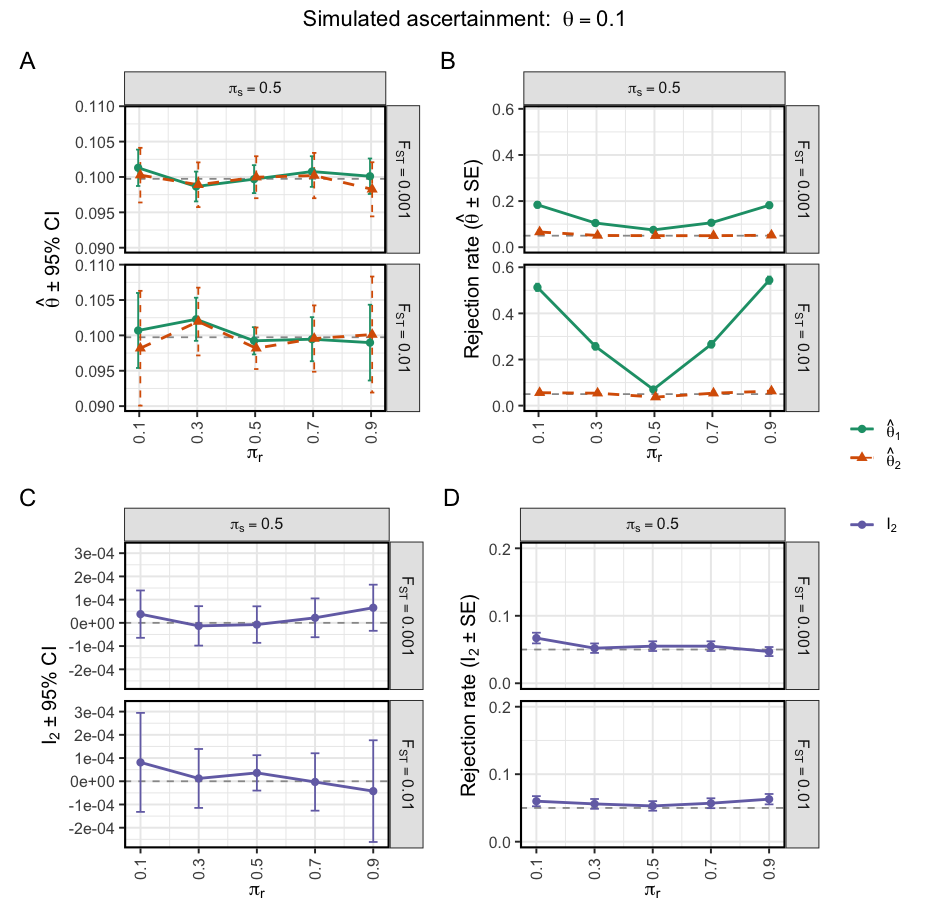

ii) **Extreme model**

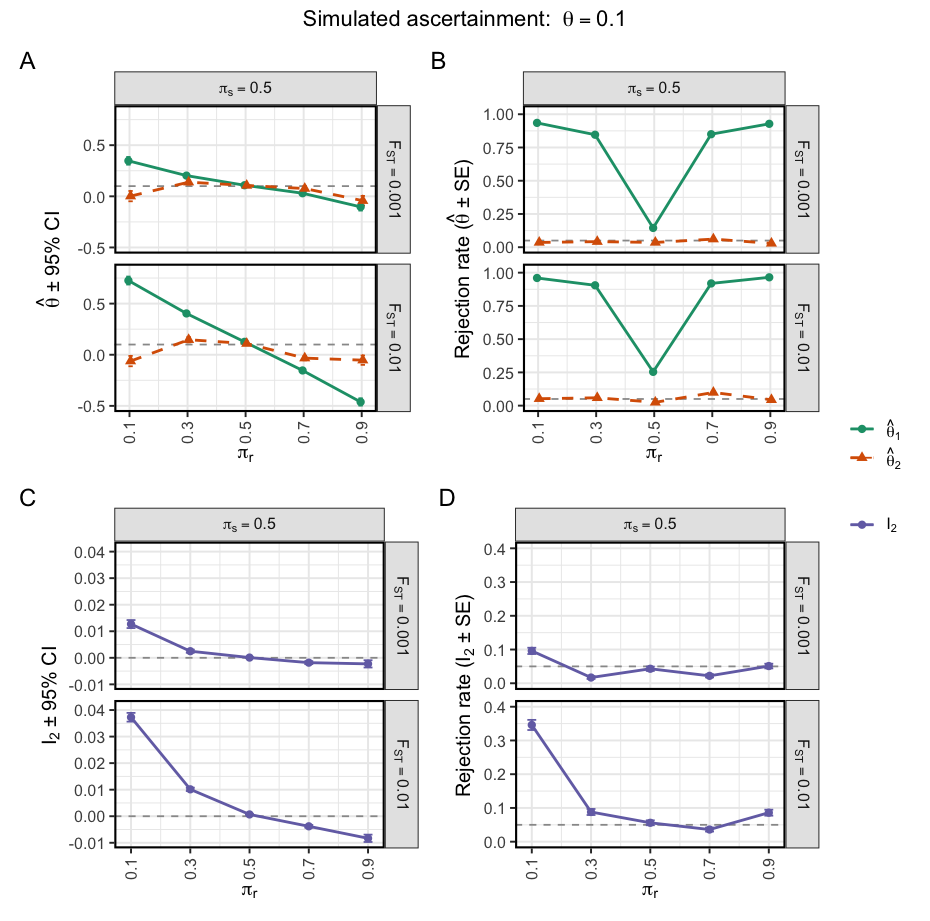

**Supplementary Fig. 5**. **Performance of** ${\hat{\boldsymbol{\theta}}}_{\boldsymbol{1}}$**,**${\hat{\boldsymbol{\theta}}}_{\boldsymbol{2}}$**and** $\boldsymbol{I}_{\boldsymbol{2}}$**under genuine ascertainment and population stratification.** Simulations were performed under a true ascertainment signal of $\theta_{true}=0.1$. Target and reference samples consisted of two sub-populations with allele frequencies generated under either a genetic drift model (i) or an extreme directional stratification model (ii) (Methods). Population differentiation was controlled by $F_{ST}\in\{0.001,0.01\}$. The target sample was simulated with equal ancestry proportions ($\pi_{s}=0.5$), whereas the reference sample was simulated with varying ancestry proportions ($\pi_{r}\in\{0.1,0.3,0.5,0.7,0.9\}$). Panels A and C show the mean estimates of $\hat{\theta}_{1}$, $\hat{\theta}_{2}$, and $I_{2}$across simulation replicates. Error bars represent 95% confidence intervals calculated as estimate $\pm1.96\times$s.e.m. Dashed horizontal lines indicate the true simulated value ($\theta_{true}=0.1$ in Panel A and $I_{2}=0$ in Panel C). Panels B and D show rejection rates at $\alpha=0.05$for the null hypotheses $H_{0}:\theta=\theta_{true}$and $H_{0}:I_{2}=0$, respectively. Error bars represent standard errors of the empirical rejection rate, calculated as $\sqrt{\hat{p}(1-\hat{p})/n}$, where $\hat{p}$is the empirical rejection rate and $n$is the number of simulation replicates. Results are based on 10,000 simulation replicates.

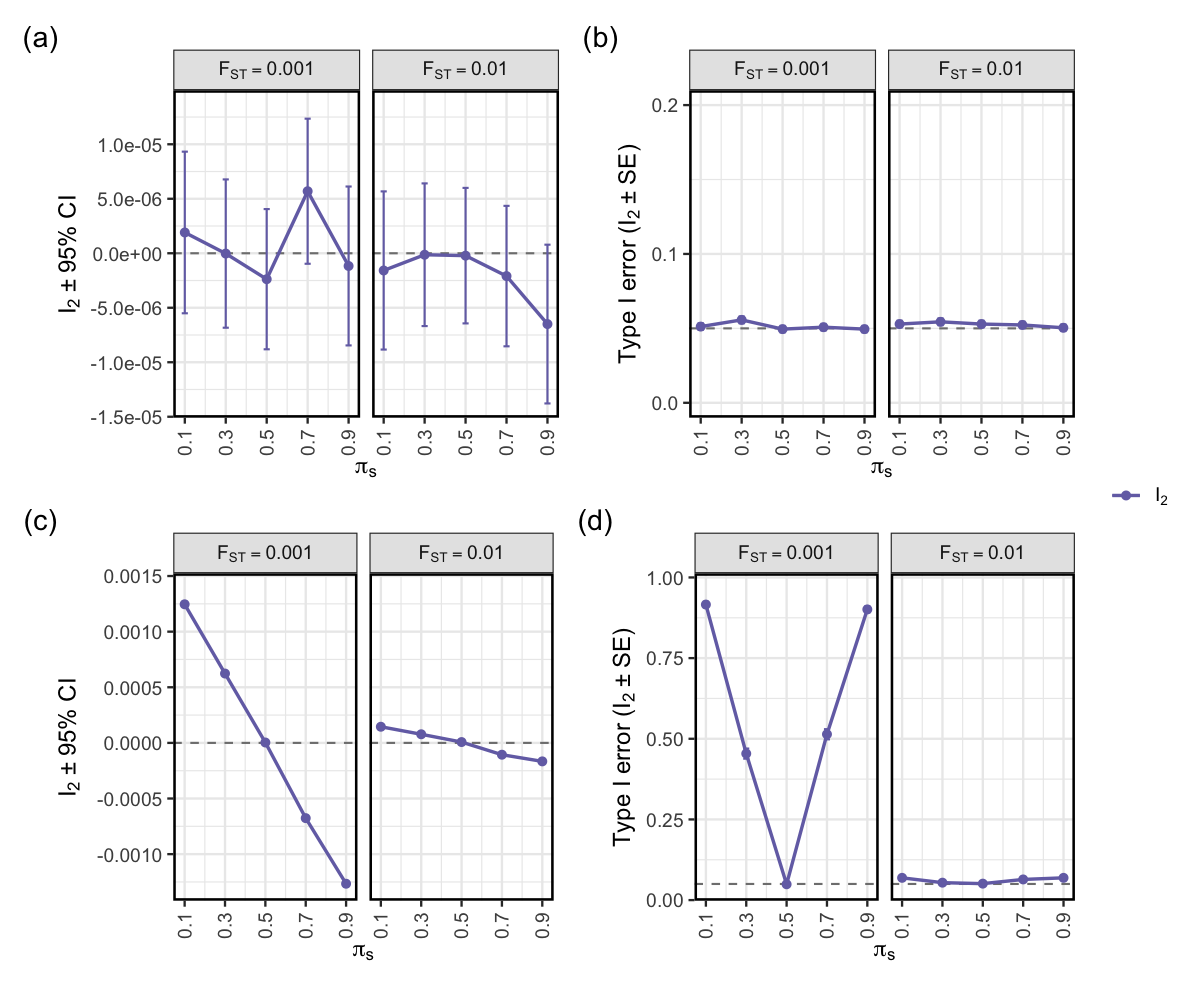

**Supplementary Fig. 6**. **Performance of** ${\hat{\boldsymbol{\theta}}}_{\boldsymbol{2}}$ **intercept (**$\boldsymbol{I}_{\boldsymbol{2}}$**) after applying the deconvolution approach under drift (top row) and extreme (bottom row) model of stratification.** Panels show results from simulations where the target sample was modelled as convex linear combination of allele frequencies from two regional reference ancestries $\left( \hat{p}^{(s)}=\pi_{s}\hat{p}^{(r1)}+\left( 1-\pi_{s} \right)\hat{p}^{(r2)} \right)$ with allele frequency differences determined by *F*_ST_ $\in\left\{ 0.001, 0.01 \right\}$. For each simulation settings, the ancestral loadings $\left( \hat{\pi}_{s} \right)$were re-estimated from sets of independent background SNPs and the estimates were used reconstruct a weighted (synthetic) reference panel (**Methods**). **Panel (a, c)** shows the mean estimates of $\hat{\theta}_{1}\pm95\% CI$ (green) and $\hat{\theta}_{2}\pm95\% CI$ (orange) under drift (top) and extreme (bottom) model. **Panel (b, d)** shows the type I error rate of $\hat{\theta}_{1}\pm SE$ (green) and $\hat{\theta}_{2}\pm SE$ (orange) under drift (top) and extreme (bottom) model. CI represent the confidence interval estimated as 1.96$\pm$s.e.m. Type I error was calculated as the fraction of *P* values below 0.05. SE represent standard error estimated as $\sqrt{\hat{p}\left( 1-p \right)}/n$, where $\hat{p}$ denotes the empirical type 1 error rate and $n$ represent the number of replicates. Simulation was performed across 10,000 replicates.

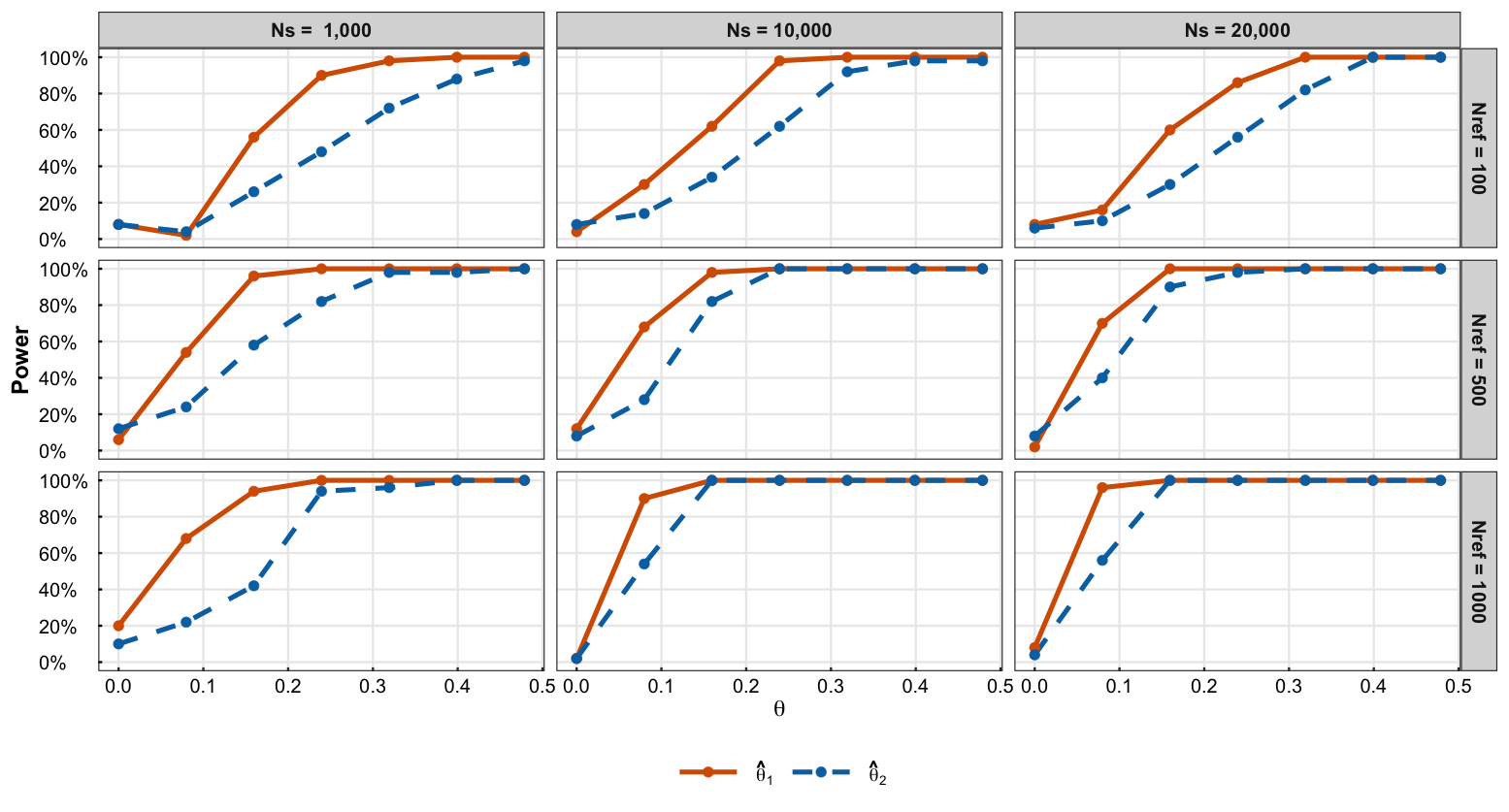

**Supplementary Fig. 7. Statistical power to detect ascertainment after deconvolution strategy.** Power was estimated from simulations under the liability threshold model for both $\hat{\theta}_{1}$(solid line) and $\hat{\theta}_{2}$(dashed line) at a significance level of $\alpha=0.05$. Power increased with the magnitude of $\theta$, but was strongly constrained by the size of the reference panel. Increasing the size of the target cohort did not improve power relative to increasing the size of the reference panel, indicating that **the precision of reference allele frequency estimates is the primary determinant of detectability.**

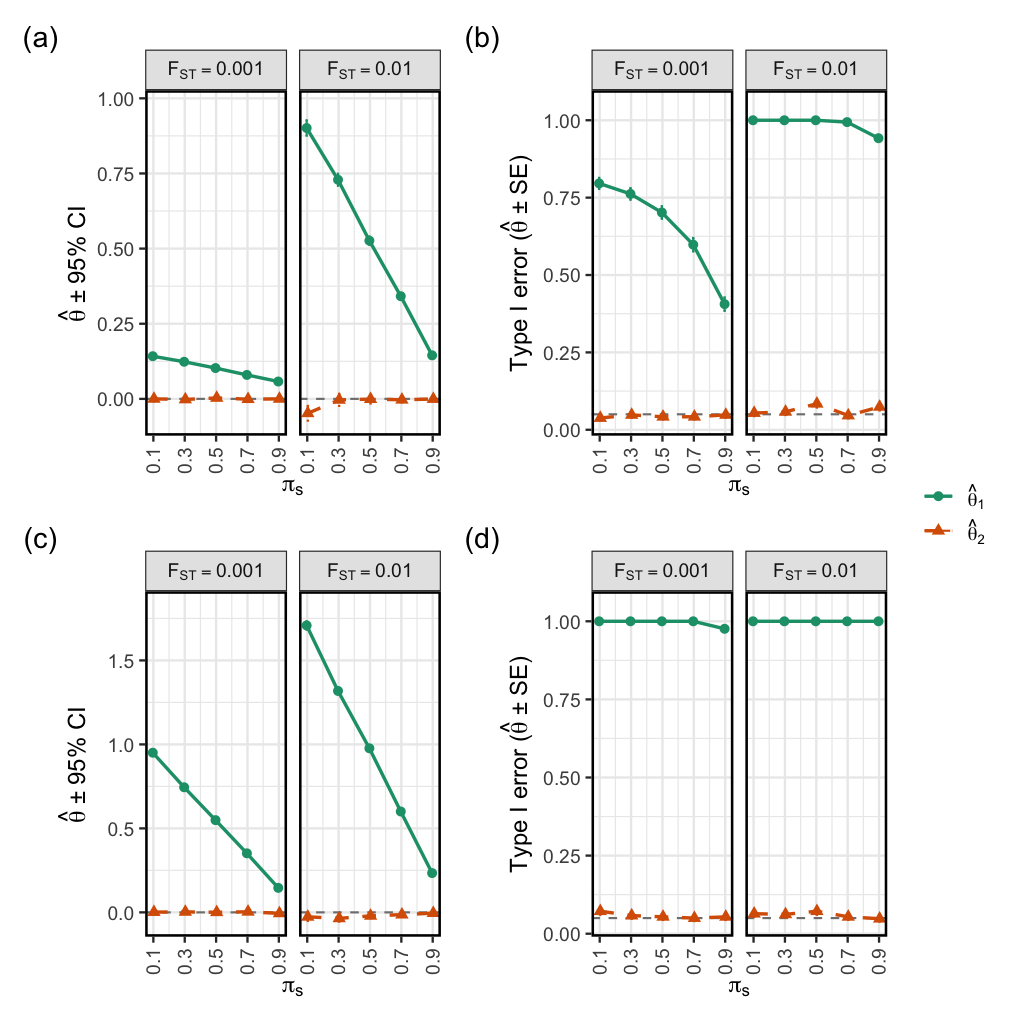

**Supplementary Fig. 8**. **Impact of missing ancestry in the reference panel on** ${\hat{\boldsymbol{\theta}}}_{\boldsymbol{1}}$ **and** ${\hat{\boldsymbol{\theta}}}_{\boldsymbol{2}}$ **under drift (top row) and extreme (bottom row) model**. Panels show simulation results assessing the robustness of $\hat{\theta}_{1}$ and $\hat{\theta}_{2}$ when the ancestry deconvolution was performed using an incomplete reference panel missing one ancestral component present in the target sample (**Methods**). Sub-populations differed by allele-frequency divergence of *F*_ST_ = 0.001 (**left**) or *F*_ST_ = 0.01 (**right**). **Panel (a, c)** shows the mean estimates of $\hat{\theta}_{1}\pm95\% CI$ (green) and $\hat{\theta}_{2}\pm95\% CI$ (orange) under drift (top) and extreme (bottom) model. **Panel (b, d)** shows the type I error rate of $\hat{\theta}_{1}\pm SE$ (green) and $\hat{\theta}_{2}\pm SE$ (orange) under drift (top) and extreme (bottom) model. CI represent the confidence interval estimated as 1.96$\pm$s.e.m. Type I error was calculated as the fraction of *P* values below 0.05. SE represent standard error estimated as $\sqrt{\hat{\alpha}\left( 1-\hat{\alpha} \right)/n}$, where $\hat{\alpha}$ denotes the empirical type 1 error rate and $n$ represent the number of replicates.

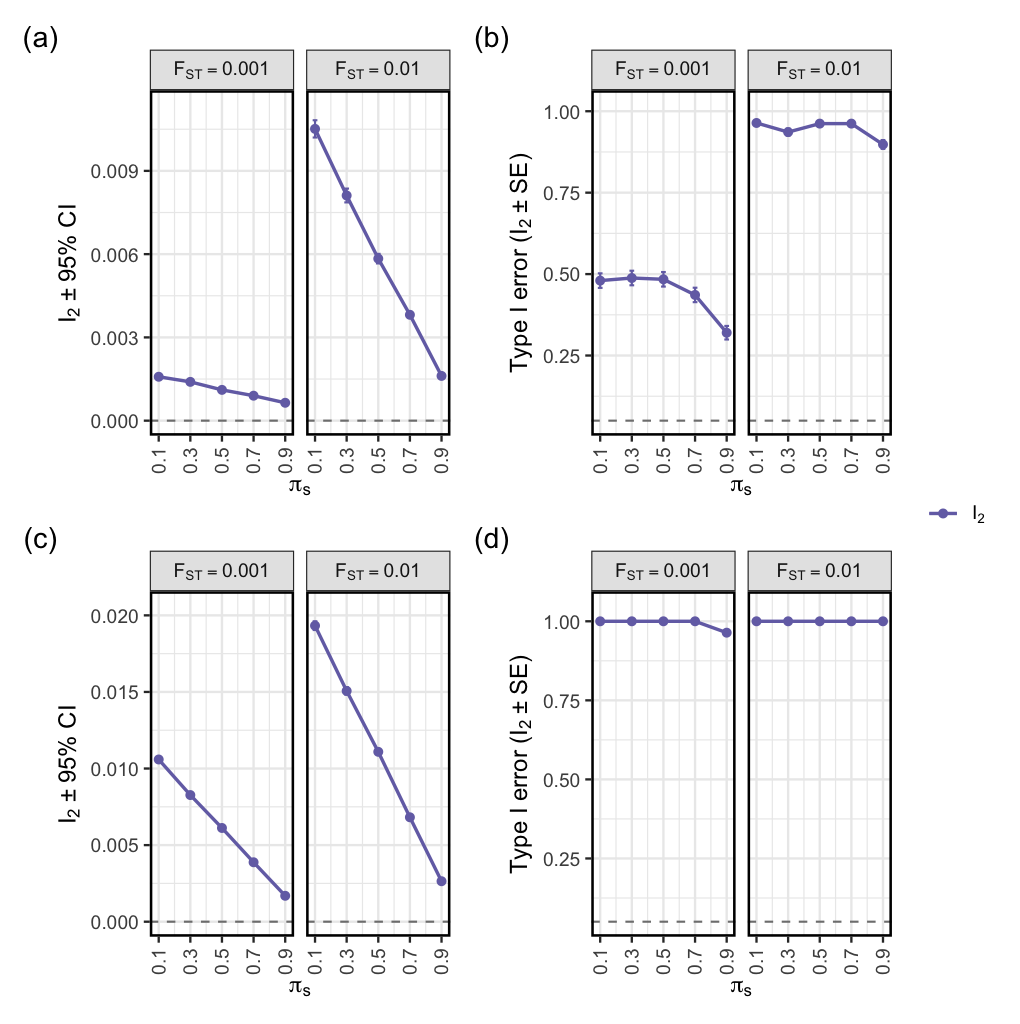

**Supplementary Fig. 9**. **Impact of missing ancestry in the reference panel on** ${\hat{\boldsymbol{\theta}}}_{\boldsymbol{2}}$ **intercept (**$\boldsymbol{I}_{\boldsymbol{2}}$**) under drift (top row) and extreme (bottom row) model**. Panels show simulation results assessing the performance of $I_{2}$ when the ancestry deconvolution was performed using an incomplete reference panel missing one ancestral component present in the target sample (**Methods**). Sub-populations differed by allele-frequency divergence of *F*_ST_ = 0.001 (**left**) or *F*_ST_ = 0.01 (**right**). **Panel (a, c)** shows the mean estimates of $I_{2}\pm95\% CI$ under drift (top) and extreme (bottom) model. **Panel (b, d)** shows the type I error rate of $I_{2}$under drift (top) and extreme (bottom) model. CI represent the confidence interval estimated as 1.96$\pm$s.e.m. Type I error was calculated as the fraction of *P* values below 0.05. SE represent standard error estimated as $\sqrt{\hat{\alpha}\left( 1-\hat{\alpha} \right)/n}$, where $\hat{\alpha}$ denotes the empirical type 1 error rate and $n$ represent the number of replicates.

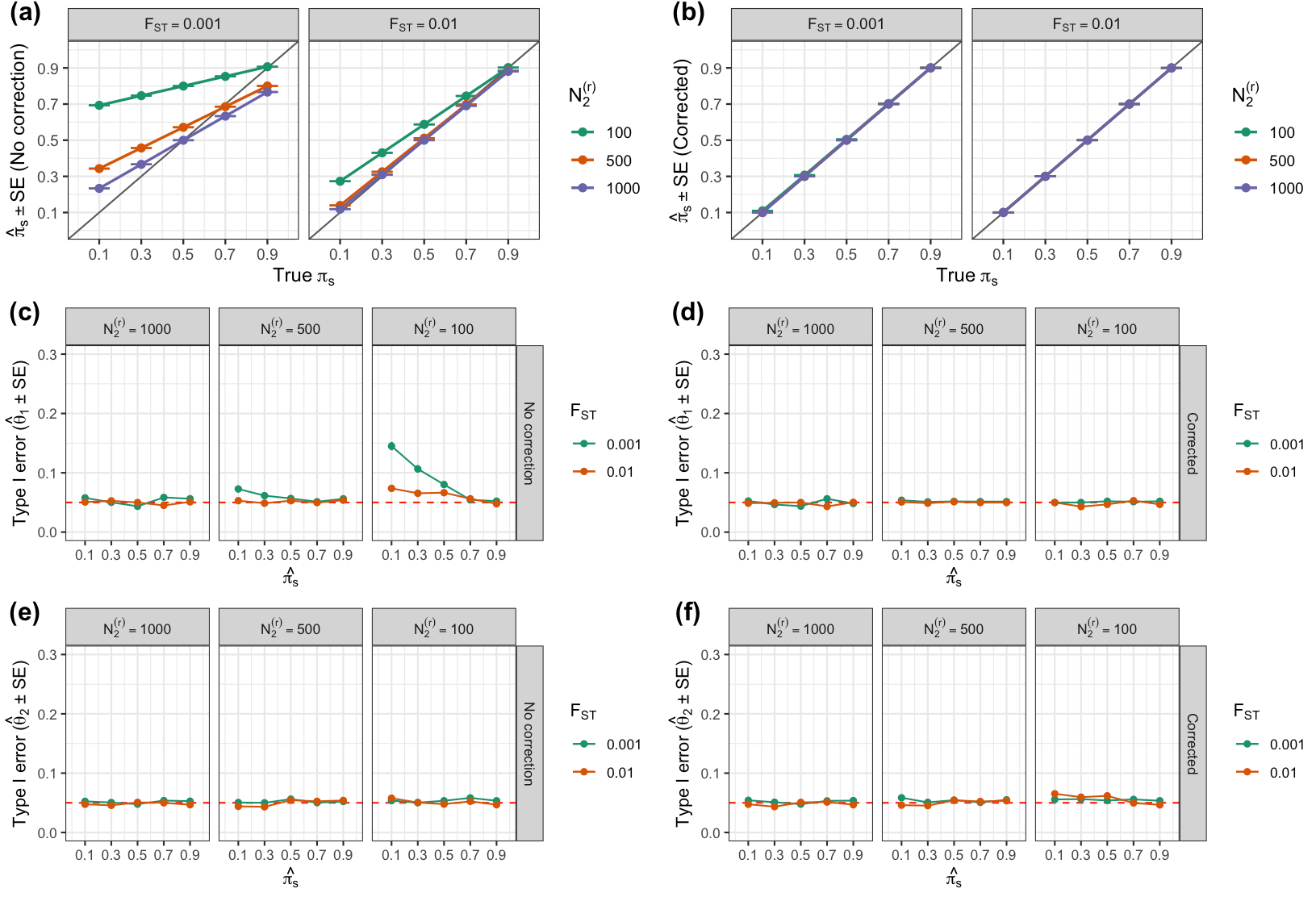

**Supplementary Fig. 10**. **Impact of sampling error in the reference panel on** ${\hat{\boldsymbol{\theta}}}_{\boldsymbol{1}}$ **and** ${\hat{\boldsymbol{\theta}}}_{\boldsymbol{2}}$ **under drift model with varying *F*_ST_**. Simulation results assessing the robustness of $\hat{\theta}_{1}$ and $\hat{\theta}_{2}$when ancestry deconvolution was performed using reference panels with unequal sample sizes across ancestries (**Methods**). The target sample was simulated as a two-way mixture of reference ancestries with $\pi_{s}\in\left\{ 0.1, 0.3, 0.5, 0.7, 0.9 \right\}$, while the sample size of the second ancestry $N_{2}^{(r)}$was systematically downsized from 1000 to 100 to introduce increasing sampling noise. Sub-populations differed by allele-frequency divergence of *F*_ST_ $\in\left\{ 0.001, 0.1 \right\}$ . **Panels (a–b)** show estimated ancestry loadings
$\left( \pi_{s}\pm sd \right)$against the true $\left( \pi_{s}\pm sd \right)$ before (**a**) and after correction (**b**). **Panel (c-d)** shows the type I error rate for $\hat{\theta}_{1}\pm SE$ before (**c**) and after correction (d). **Panel (e-f)** shows the type I error rate for $\hat{\theta}_{2}\pm SE$ before (**e**) and after correction (**f**). SE represents standard error. Simulation was performed across 10, 000 replicates. Type I error was calculated as the fraction of *P* values below 0.05.

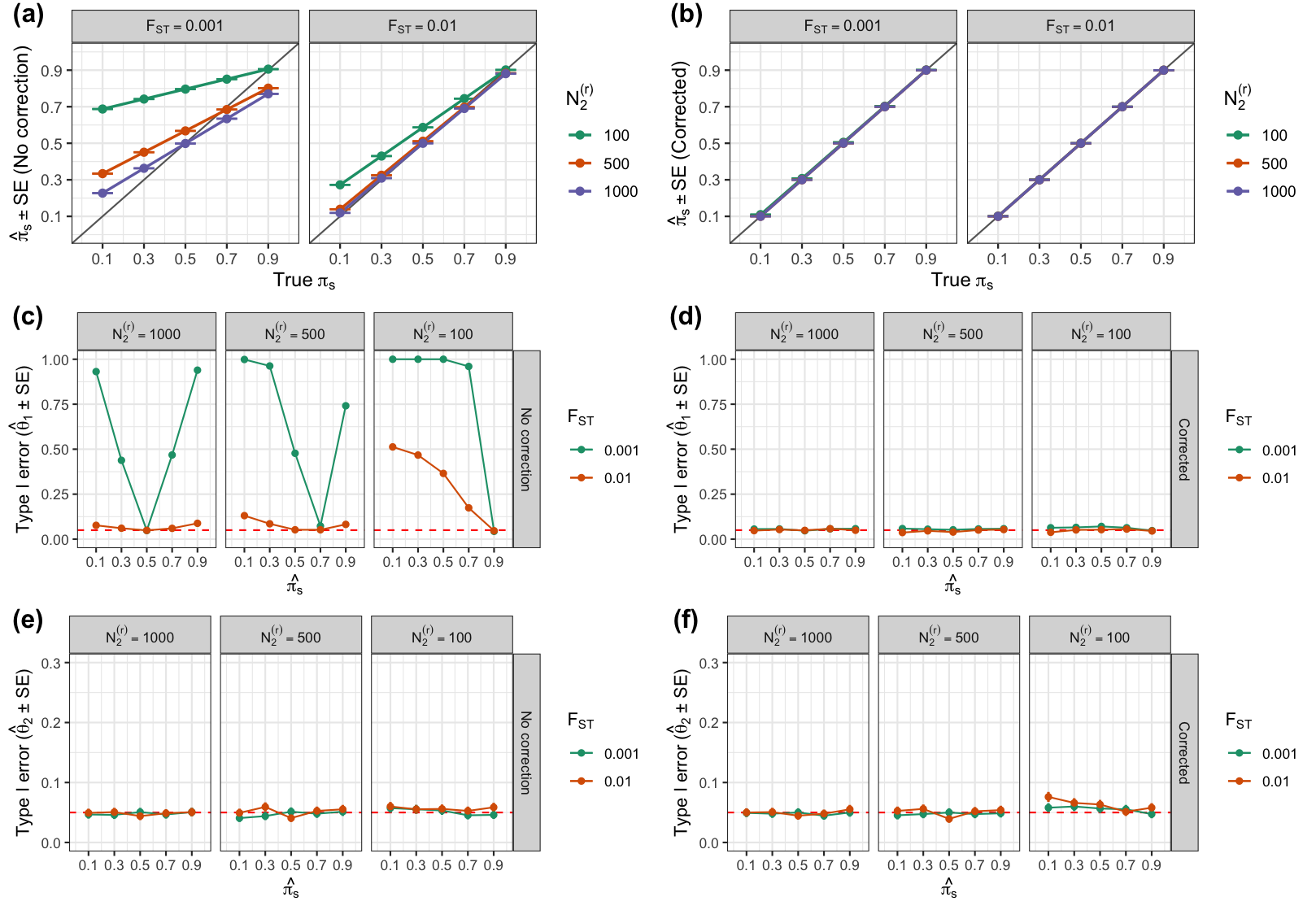

**Supplementary Fig. 11**. **Impact of sampling error in the reference panel on** ${\hat{\boldsymbol{\theta}}}_{\boldsymbol{1}}$ **and** ${\hat{\boldsymbol{\theta}}}_{\boldsymbol{2}}$ **under extreme model with varying *F*_ST_**. Simulation results assessing the robustness of $\hat{\theta}_{1}$ and $\hat{\theta}_{2}$when ancestry deconvolution was performed using reference panels with unequal sample sizes across ancestries (**Methods**). The target sample was simulated as a two-way mixture of reference ancestries with $\pi_{1}^{\left( s \right)}\in\left\{ 0.1, 0.3, 0.5, 0.7, 0.9 \right\}$, while the sample size of the second ancestry $N_{2}^{(r)}$was systematically downsized from 1000 to 100 to introduce increasing sampling noise. Sub-populations differed by allele-frequency divergence of *F*_ST_ $\in\left\{ 0.001, 0.1 \right\}$ . **Panels (A–B)** show estimated ancestry loadings
$\left( \hat{\pi}_{1}^{\left( s \right)}\pm sd \right)$against the true $\left( \pi_{1}^{\left( s \right)}\pm sd \right)$ before (**A**) and after correction (**B**). **Panel (C-D)** shows the type I error rate for $\hat{\theta}_{1}\pm SE$ before (**C**) and after correction (**D**). **Panel (E-F)** shows the type I error rate for $\hat{\theta}_{2}\pm SE$ before (**C**) and after correction (**D**). SE represents standard error. Simulation was performed across 10, 000 replicates. Type I error was calculated as the fraction of *P* values below 0.05.

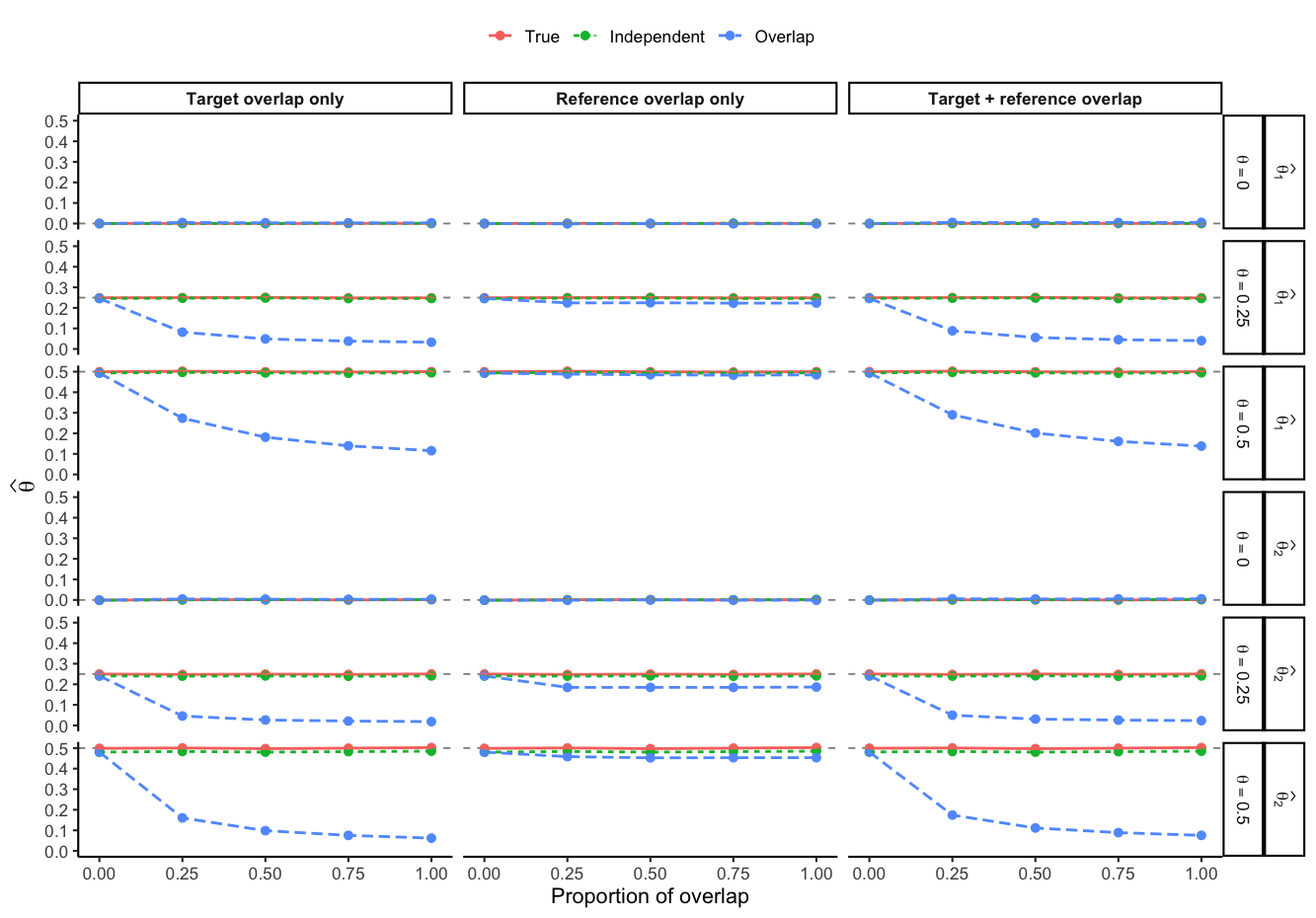

**Supplementary Fig. 12**. **Impact of sample overlap between the GWAS discovery cohort and target or reference samples on** $\hat{\theta}_{1}$**and** $\hat{\theta}_{2}$**.** Simulation results assessing the robustness of $\hat{\theta}_{1}$and $\hat{\theta}_{2}$when individuals from the target sample, reference panel, or both were included in the GWAS discovery cohort used to estimate SNP effects. Simulated ascertainment was varied across $\theta\in\{0,0.25,0.5\}$, and the proportion of sample overlap was varied from no overlap (0) to full overlap (1). Columns show target overlap only, reference overlap only, and target and reference overlap. Rows show estimates of $\hat{\theta}_{1}$and $\hat{\theta}_{2}$under each simulated ascertainment setting. Red, green, and blue lines denote estimates using true effects, an independent GWAS discovery sample, and a GWAS discovery sample with overlap, respectively. The grey dashed line denotes the simulated value of $\theta$. Points represent mean estimates across simulation replicates, with error bars denoting standard errors.

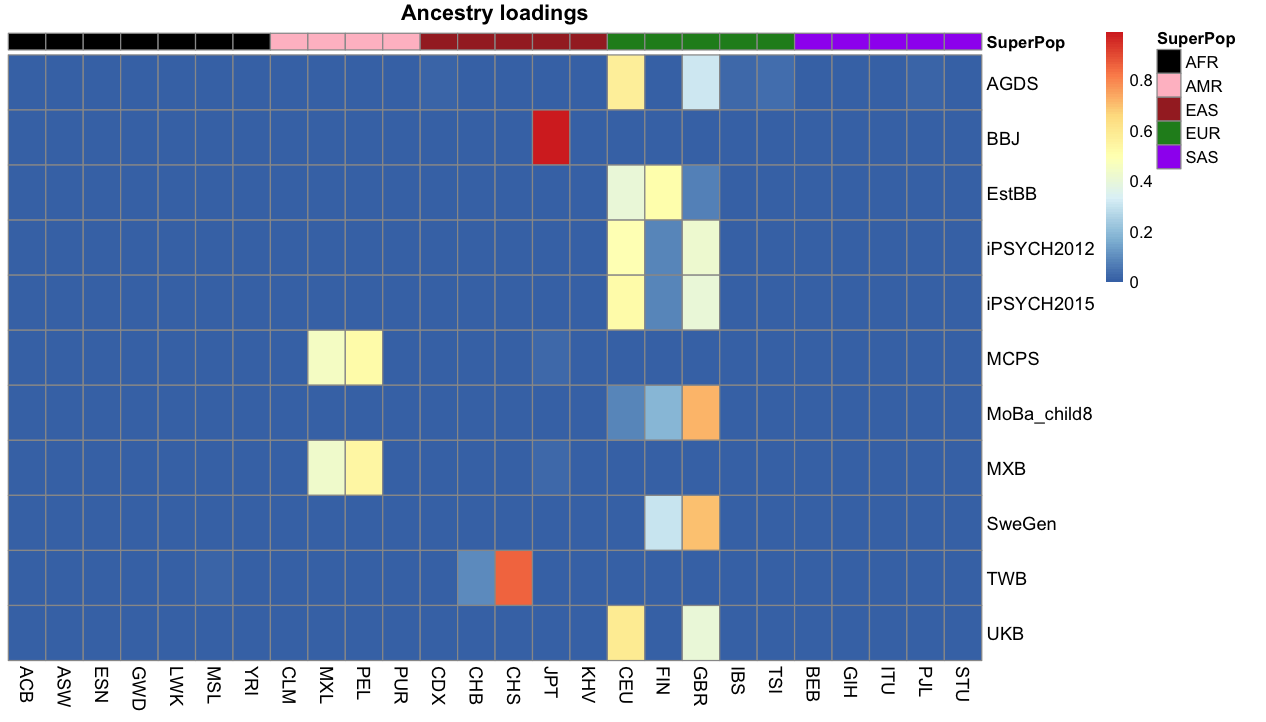

**Supplementary Fig. 13. Ancestral loadings and their corresponding superpopulations across biobanks.** Panel shows the ancestry loadings inferred across biobanks using 26 ancestral populations from the 1000 Genomes Project (1KGP) after correcting for attenuation-bias in the response variables (i.e., 1KGP). Each cell represents the estimated ancestry loading of a biobank (rows) across the 26 ancestry groups of the 1KGP reference populations (columns), with colour intensity reflecting the relative contribution of that ancestry. The colour bar above the columns indicates the corresponding 1KGP super-population (SuperPop; AFR = African, AMR = Admixed American, EAS = East Asian, EUR = European, SAS = South Asian).

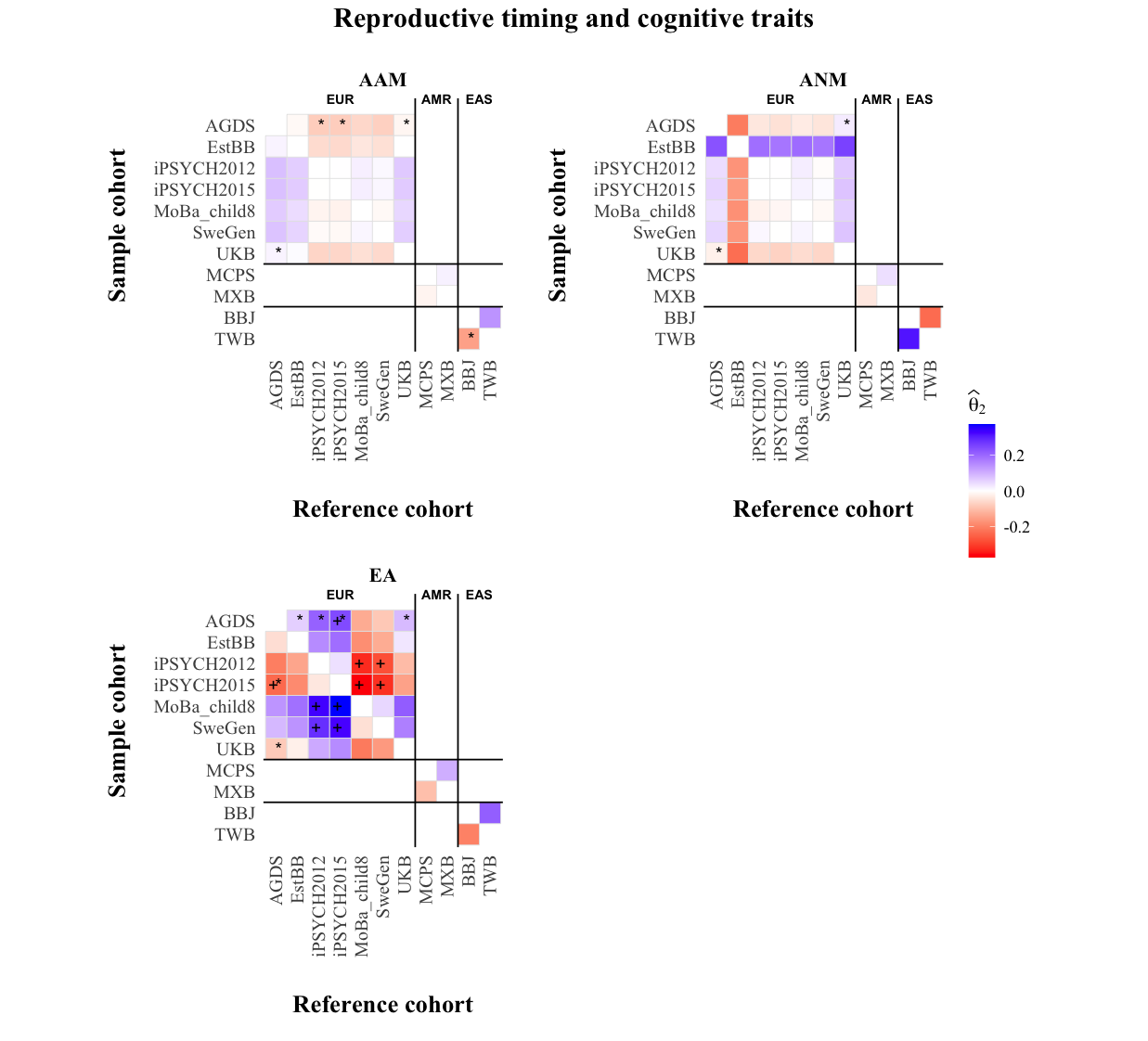

**Supplementary Fig. 14a**. **Relative ascertainment for reproductive timing and cognitive traits across within-ancestry biobank comparisons.** Heatmaps show estimates of $\hat{\theta}_{2}$ for age at menarche (AAM), age at natural menopause (ANM), and educational attainment (EA) across pairwise biobank comparisons within European (EUR), admixed American (AMR), and East Asian (EAS) ancestry groups. The biobanks includes the Australian Genomics Depression Study (AGDS), Biobank Japan (BBJ), the Estonian biobank (EstBB), the 2012 and 2015 waves of the Integrative Psychiatric Research biobank (iPSYCH2012 and iPSYCH2015), the Mexican biobank (MXB), the children cohort (age 8) from the Norwegian Mother and Child Cohort Study (MoBa_child8), the Mexican city prospective cohort study (MCPS), the Swedish population biobank (SweGen), the Taiwan biobank (TWB), and the UK Biobank (UKB)). Each tile represents the estimated relative ascertainment signal between a sample cohort (rows) and a reference cohort (columns), using ancestry-matched sets of independently associated SNPs selected through LD clumping based on the corresponding 1000 Genomes Project (1KGP) super-population reference panel. Positive (negative) and statistically significant value of $\hat{\theta}_{2}$implies that the sample cohort is enriched for individuals with higher (lower) genetic propensity or liability for the target trait compared to the reference. The Plus sign (+) indicates significant $\hat{\theta}_{2}$ estimates (*P* values < 0.0001) after Bonferroni correction across all within-ancestry pairwise comparisons and investigated traits. Asterisks (*) indicate estimates with significant intercept (*I₂*; *P* values < 0.0001) after Bonferroni correction. +* indicate when both are significant. Comparisons were restricted to cohorts belonging to the same broad ancestry group.

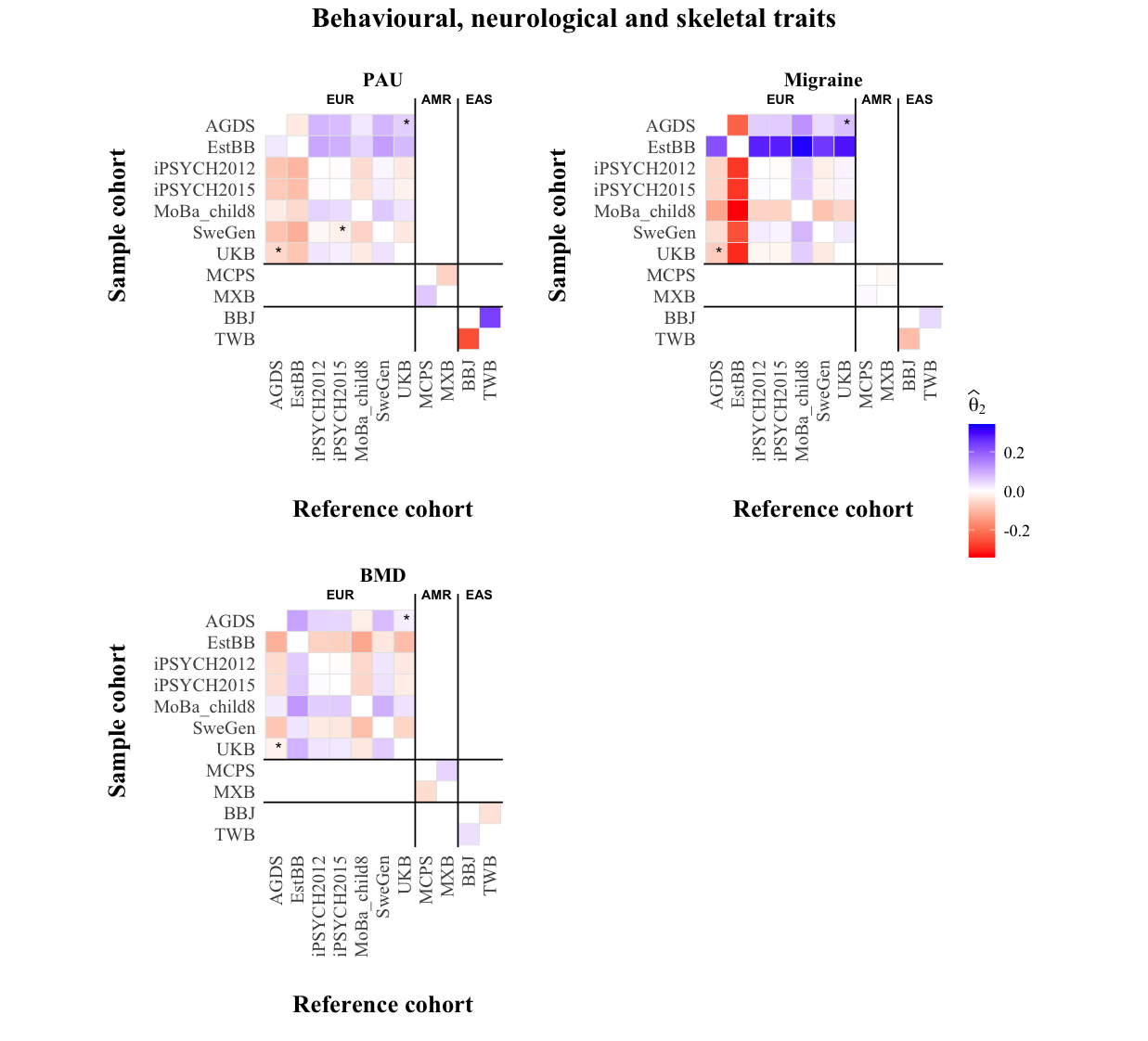

**Supplementary Fig. 14b**. **Relative ascertainment for behavioural, neurological, and skeletal traits across within-ancestry biobank comparisons**. Heatmaps show estimates of $\hat{\theta}_{2}$ for problematic alcohol use (PAU), migraine, and bone mineral density (BMD) across pairwise biobank comparisons within European (EUR), admixed American (AMR), and East Asian (EAS) ancestry groups. The biobanks includes the Australian Genomics Depression Study (AGDS), Biobank Japan (BBJ), the Estonian biobank (EstBB), the 2012 and 2015 waves of the Integrative Psychiatric Research biobank (iPSYCH2012 and iPSYCH2015), the Mexican biobank (MXB), the children cohort (age 8) from the Norwegian Mother and Child Cohort Study (MoBa_child8), the Mexican city prospective cohort study (MCPS), the Swedish population biobank (SweGen), the Taiwan biobank (TWB), and the UK Biobank (UKB)). Each tile represents the estimated relative ascertainment signal between a sample cohort (rows) and a reference cohort (columns), using ancestry-matched sets of independently associated SNPs selected through LD clumping based on the corresponding 1000 Genomes Project (1KGP) super-population reference panel. Positive (negative) and statistically significant value of $\hat{\theta}_{2}$implies that the sample cohort is enriched for individuals with higher (lower) genetic propensity or liability for the target trait compared to the reference. The Plus sign (+) indicates significant $\hat{\theta}_{2}$ estimates (*P* values < 0.0001) after Bonferroni correction across all within-ancestry pairwise comparisons and investigated traits. Asterisks (*) indicate estimates with significant intercept (*I₂*; *P* values < 0.0001) after Bonferroni correction. +* indicate when both are significant. Comparisons were restricted to cohorts belonging to the same broad ancestry group.

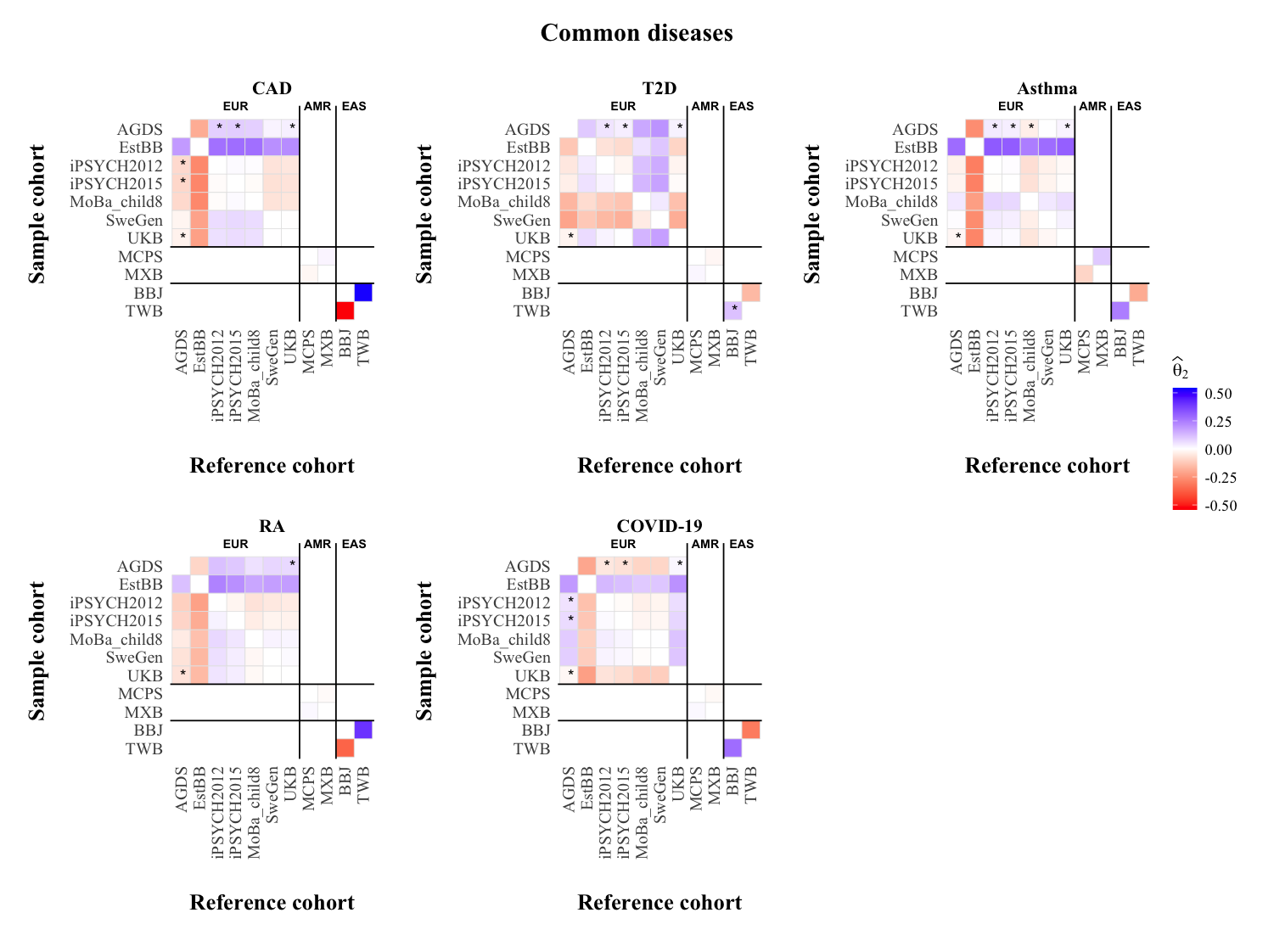

**Supplementary Fig. 14c. Relative ascertainment for common disease traits across within-ancestry biobank comparisons.** Heatmaps show estimates of $\hat{\theta}_{2}$ for coronary artery disease (CAD), type 2 diabetes (T2D), asthma, rheumatoid arthritis (RA), and COVID-19 across pairwise biobank comparisons within European (EUR), admixed American (AMR), and East Asian (EAS) ancestry groups. The biobanks includes the Australian Genomics Depression Study (AGDS), Biobank Japan (BBJ), the Estonian biobank (EstBB), the 2012 and 2015 waves of the Integrative Psychiatric Research biobank (iPSYCH2012 and iPSYCH2015), the Mexican biobank (MXB), the children cohort (age 8) from the Norwegian Mother and Child Cohort Study (MoBa_child8), the Mexican city prospective cohort study (MCPS), the Swedish population biobank (SweGen), the Taiwan biobank (TWB), and the UK Biobank (UKB)). Each tile represents the estimated relative ascertainment signal between a sample cohort (rows) and a reference cohort (columns), using ancestry-matched sets of independently associated SNPs selected through LD clumping based on the corresponding 1000 Genomes Project (1KGP) super-population reference panel. Positive (negative) and statistically significant value of $\hat{\theta}_{2}$implies that the sample cohort is enriched for individuals with higher (lower) genetic propensity or liability for the target trait compared to the reference. The Plus sign (+) indicates significant $\hat{\theta}_{2}$ estimates (*P* values < 0.0001) after Bonferroni correction across all within-ancestry pairwise comparisons and investigated traits. Asterisks (*) indicate estimates with significant intercept (*I₂*; *P* values < 0.0001) after Bonferroni correction. +* indicate when both are significant. Comparisons were restricted to cohorts belonging to the same broad ancestry group.

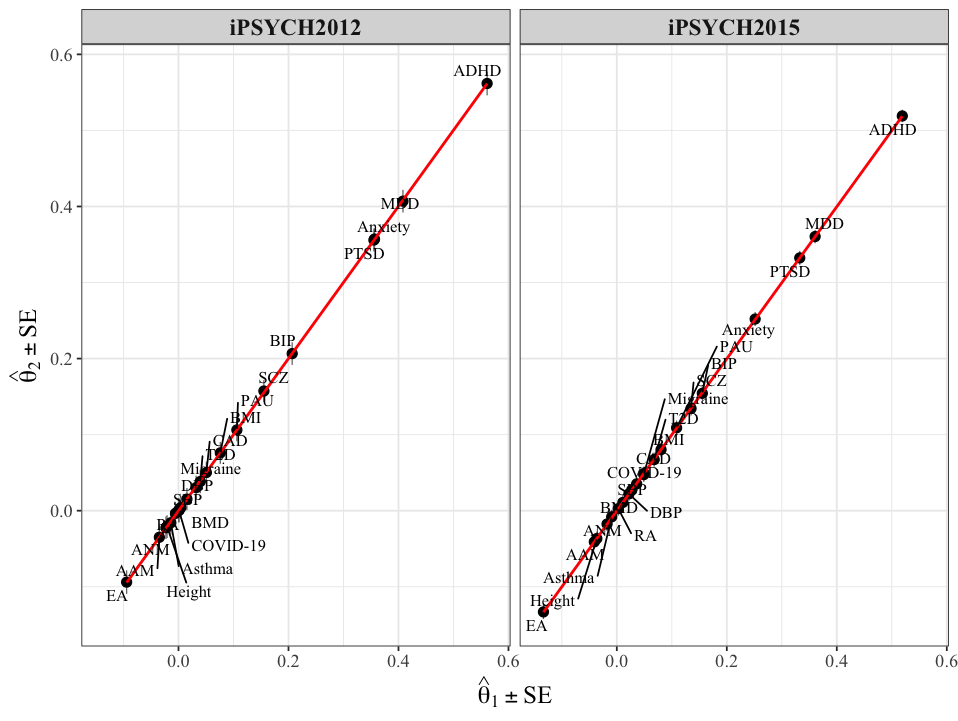

#### **Supplementary Fig. 15. Relationship between ascertainment estimates obtained using** $\hat{\theta}_{2}$**and** $\hat{\theta}_{1}$**in the iPSYCH cohorts** across 21 complex traits. Scatter plots comparing $\hat{\theta}_{2}$ (y-axis) and $\hat{\theta}_{1}$(x-axis) for iPSYCH2012 (left) and iPSYCH2015 (right). Each point represents one trait. Red lines indicate the fitted linear regression within each cohort. Estimates were largely in concordance across traits, with the largest ascertainment signals observed for psychiatric disorders, including ADHD, MDD, anxiety, and PTSD.

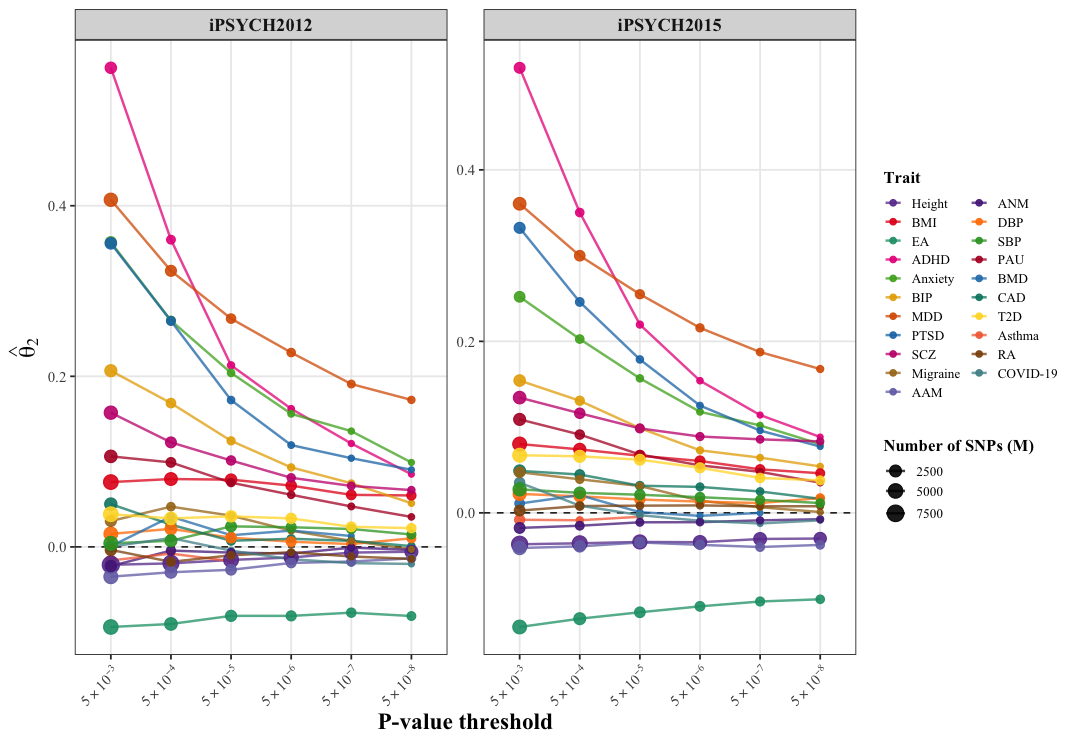

**Supplementary Fig. 16**. **Sensitivity of** ${\hat{\boldsymbol{\theta}}}_{\boldsymbol{2}}$ **to SNP inclusion thresholds.** Panel shows the estimates of $\hat{\theta}_{2}$ across 21 complex traits and diseases in the 2012 and 2015 waves of the Integrative Psychiatric Research (iPSYCH2012 and iPSYCH2015) biobank using a census-representative reference constructed from population controls.

| (a) | 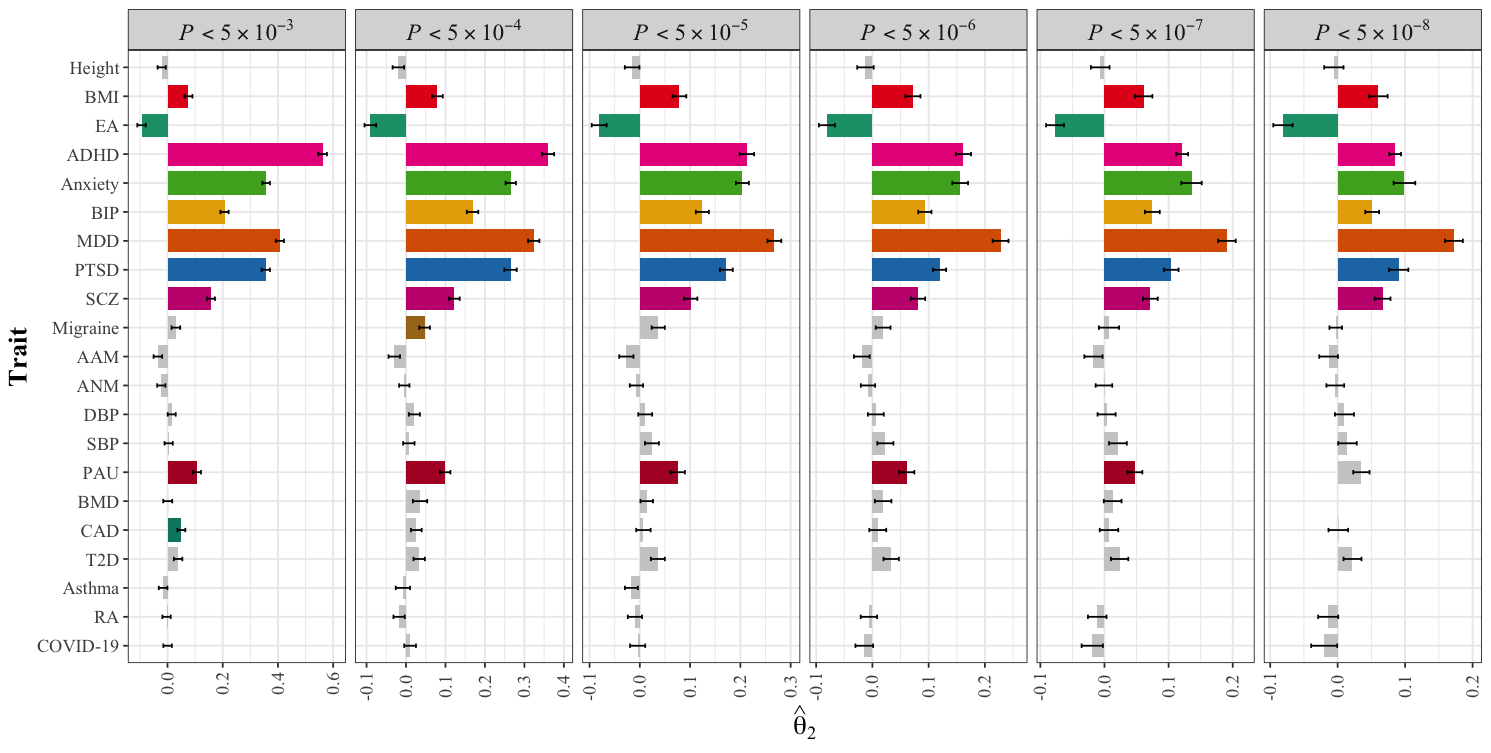 |
| --- | --- |
| (b) | 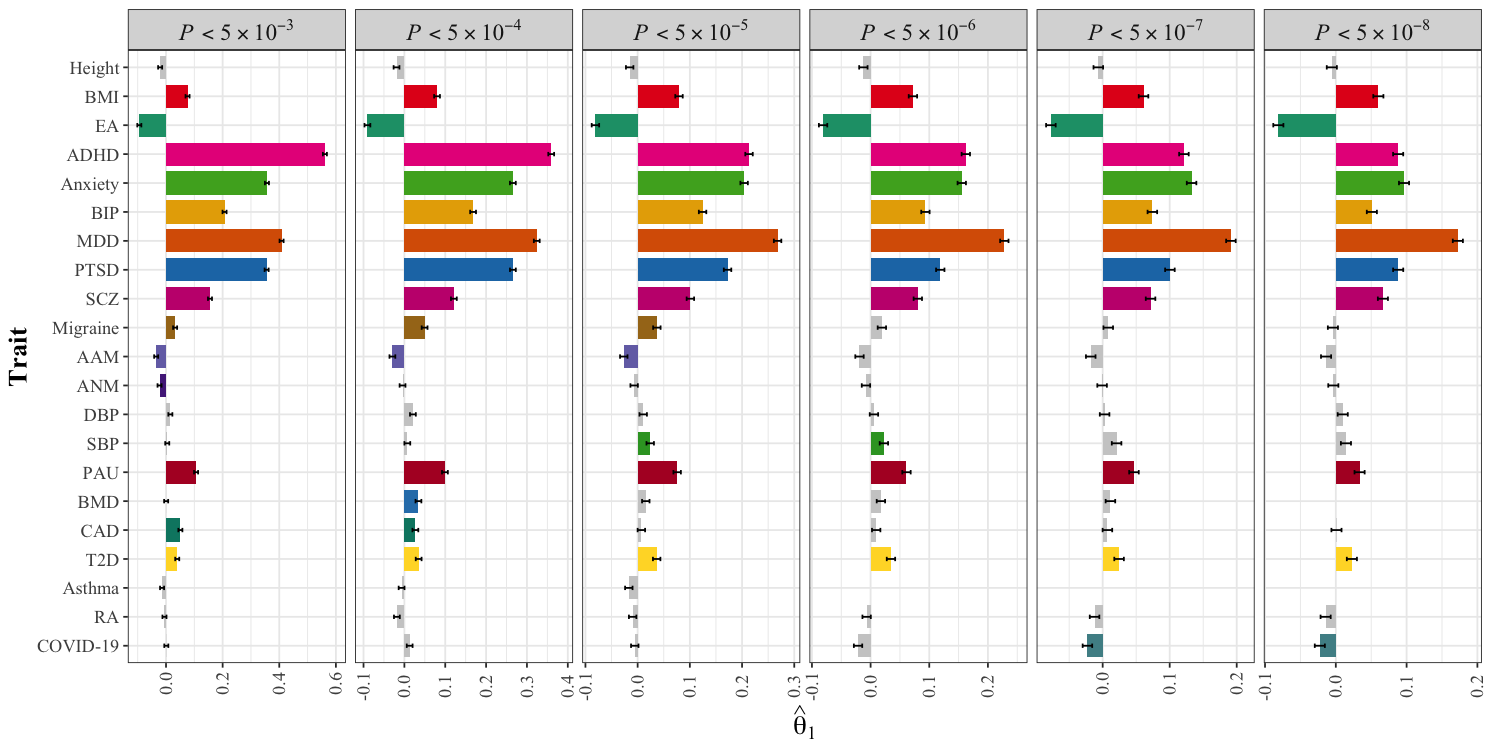 |

**Supplementary Fig. 17**. **Ascertainment estimates in iPYCH2012 across different SNPs with varying *P* values threshold. Panel (a)** shows estimates of $\hat{\theta}_{2}$ (top) and **(b)** the corresponding estimates of ${\hat{\boldsymbol{\theta}}}_{\boldsymbol{1}}$ (bottom) across 21 complex traits and diseases. Each panel corresponds to selected SNPs with *P* values threshold (from *P* < $5\times{10}^{-03}$ to *P* < $5\times{10}^{-08}$). Bars represent the point estimates $\pm$ standard error. Grey bars denote non-significant results, while colored bars highlight statistically significant estimates (*P* values < 0.0024). Asterisks indicate estimates with significant $I_{2}$(*P* values < 0.0024) and are shown for $\hat{\theta}_{2}$ only. Positive (negative) and statistically significant ascertainment values implies that the sample is enriched for individuals with higher (lower) genetic propensity for the target trait compared to the reference. Only traits with SNPs≥10 SNPs are shown.

| (a) | 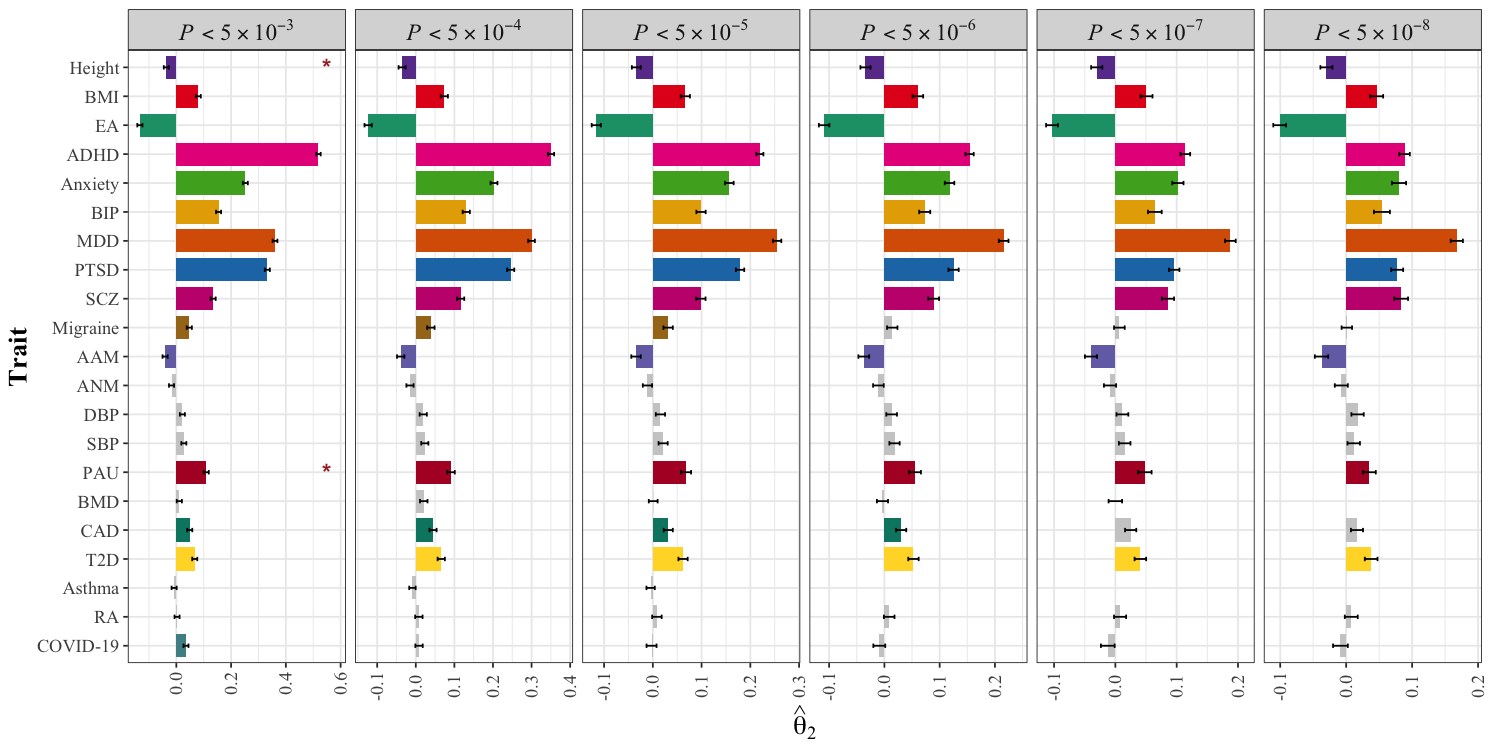 |
| --- | --- |
| (b) | 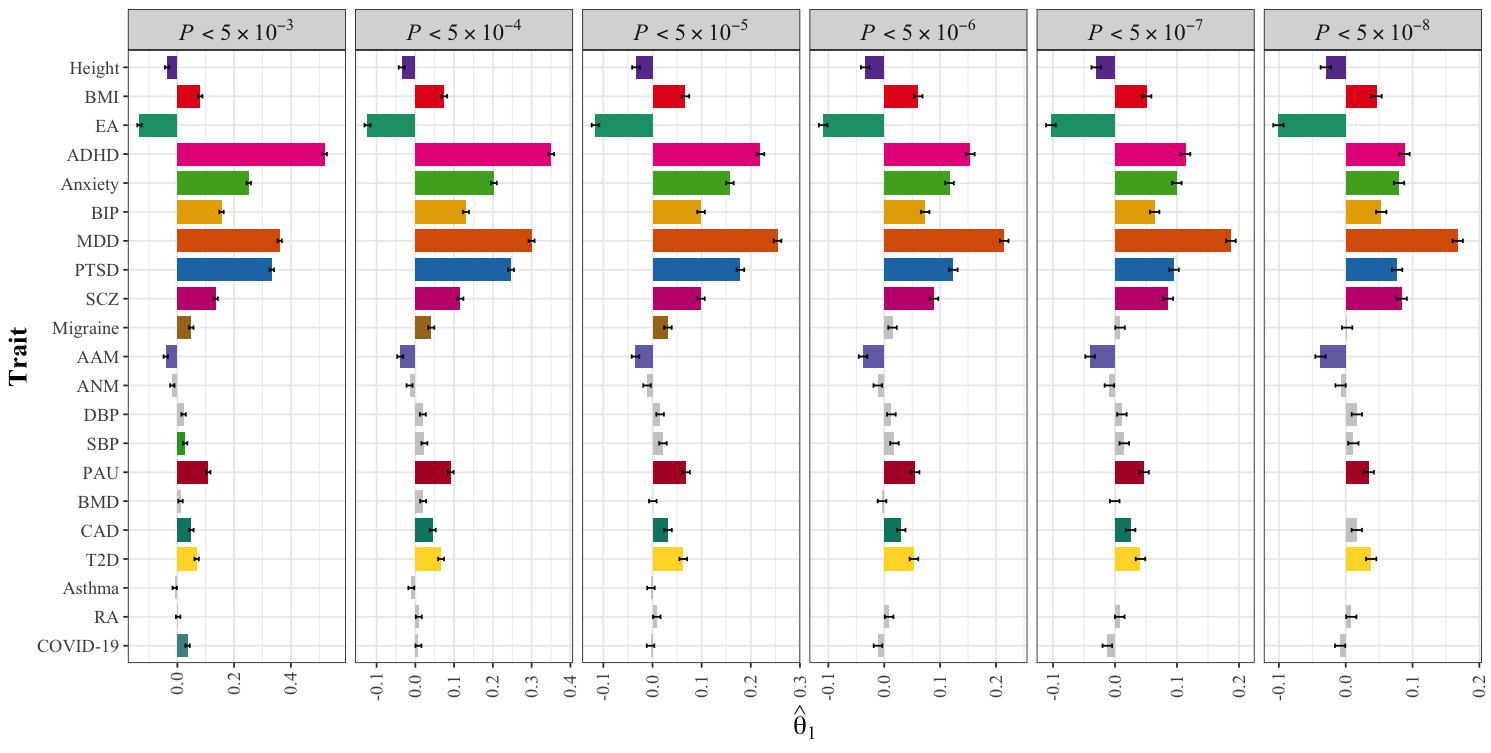 |

**Supplementary Fig. 18**. **Ascertainment estimates in iPYCH2015 across different SNPs with varying *P* values threshold. Panel (a)** shows estimates of $\hat{\theta}_{2}$ (top) and **(b)** the corresponding estimates of ${\hat{\boldsymbol{\theta}}}_{\boldsymbol{1}}$ (bottom) across 21 complex traits and diseases. Each panel corresponds to selected SNPs with *P* values threshold (from *P* < $5\times{10}^{-03}$ to *P* < $5\times{10}^{-08}$). Bars represent the point estimates $\pm$ standard error. Grey bars denote non-significant results, while colored bars highlight statistically significant estimates (*P* values < 0.0024). Asterisks indicate estimates with significant $I_{2}$(*P* values < 0.0024) and are shown for $\hat{\theta}_{2}$ only. Positive (negative) and statistically significant ascertainment values implies that the sample is enriched for individuals with higher (lower) genetic propensity for the target trait compared to the reference. Only traits with SNPs≥10 SNPs are shown.

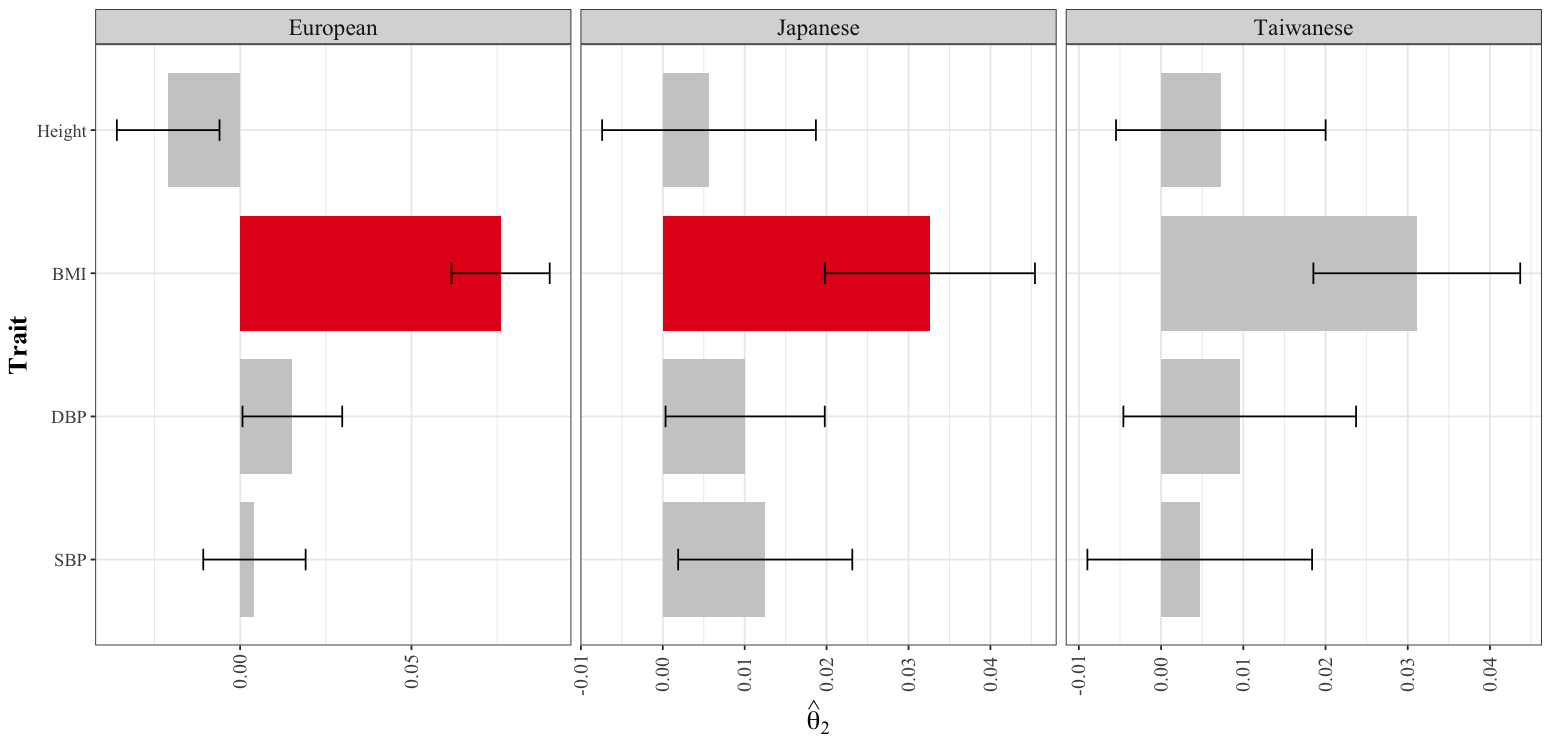

**Supplementary Fig. 19. Robustness of** ${\hat{\boldsymbol{\theta}}}_{\boldsymbol{2}}$ **to the choice of discovery GWAS in iPSYCH2012 cohort.** Each panel shows the result of four representative traits (height, systolic blood pressure (SBP), diastolic blood pressure (DBP) and body mass index (BMI)) when assessing sensitivity to the GWAS discovery effect sizes. Approximately independent SNPs (*P* < $5\times{10}^{-03}$) defined using the European LD panel from the 1KGP as in the main analyses, were used across all ancestry-specific GWAS comparisons. The numbers of approximately independent SNPs retained for European-, Japanese-, and Taiwanese-derived GWAS, respectively, were: $M_{Height}^{EUR}=8361, M_{Height}^{JPN}=1225,M_{Height}^{TWN}=794;$ $M_{BMI}^{EUR}=5134, M_{BMI}^{JPN}=401,M_{BMI}^{TWN}=293;M_{DBP}^{EUR}=4185, M_{DBP}^{JPN}=137,M_{DBP}^{TWN}=190;$ $M_{SBP}^{EUR}=4379, M_{SBP}^{JPN}=174,M_{SBP}^{TWN}=195$. The bars represent the points estimate for $\hat{\theta}_{2}\pm$ standard error. Grey bars denote non-significant results, while colored bars highlight significant estimates. Asterisks indicate estimates with significant $I_{2}$. Positive (negative) and statistically significant value of $\hat{\theta}_{2}$implies that the sample is enriched for individuals with higher (lower) genetic propensity for the target trait compared to the reference.

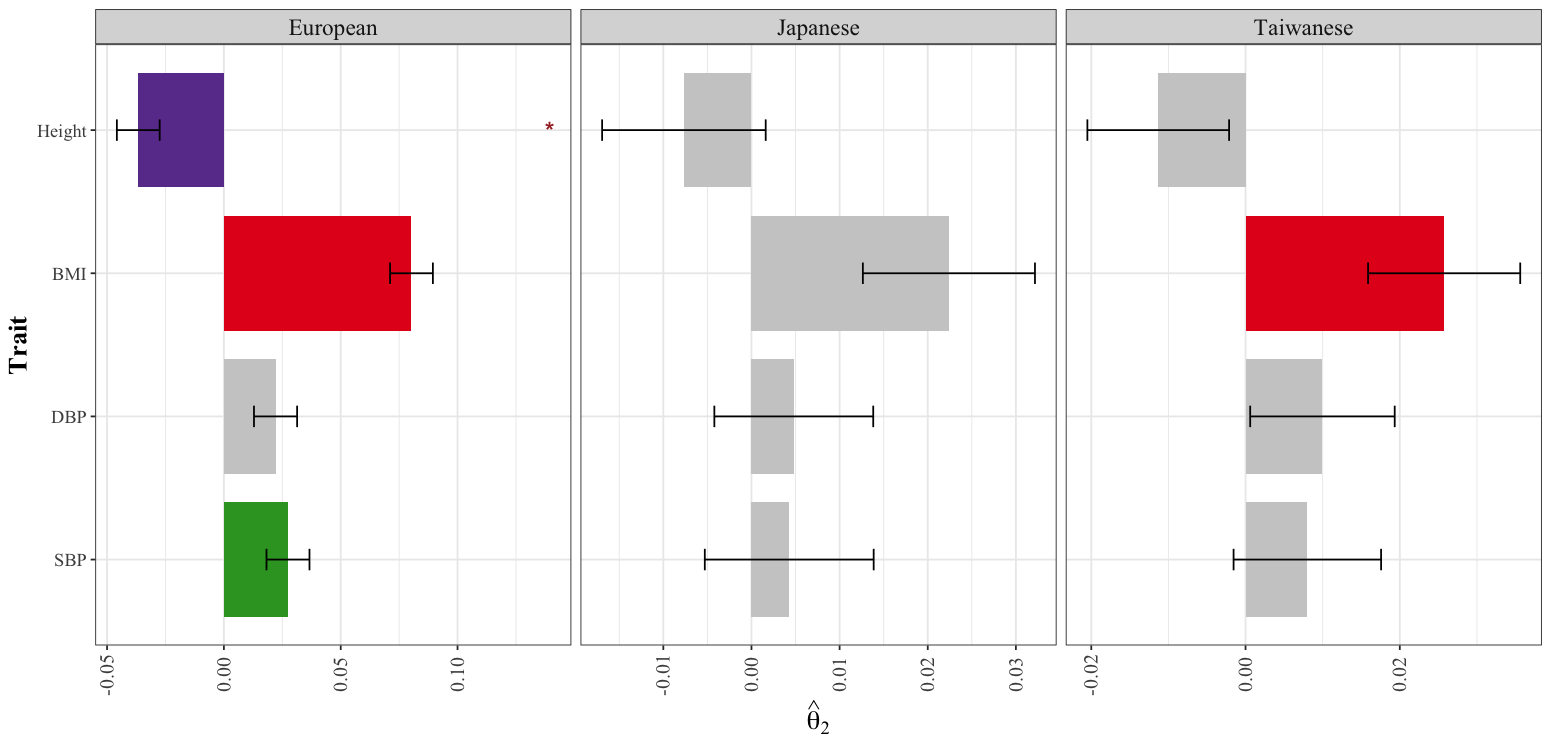

**Supplementary Fig. 20. Robustness of** ${\hat{\boldsymbol{\theta}}}_{\boldsymbol{2}}$ **to the choice of discovery GWAS in iPSYCH2015 cohort.** Each panel shows the result of four representative traits (height, systolic blood pressure (SBP), diastolic blood pressure (DBP) and body mass index (BMI)) when assessing sensitivity to the GWAS discovery effect sizes. Approximately independent SNPs (*P* < $5\times{10}^{-03}$) defined using the European LD panel from the 1KGP as in the main analyses, were used across all ancestry-specific GWAS comparisons. The numbers of approximately independent SNPs retained for European-, Japanese-, and Taiwanese-derived GWAS, respectively, were: $M_{Height}^{EUR}=8361, M_{Height}^{JPN}=1225,M_{Height}^{TWN}=794;$ $M_{BMI}^{EUR}=5134, M_{BMI}^{JPN}=401,M_{BMI}^{TWN}=293;M_{DBP}^{EUR}=4185, M_{DBP}^{JPN}=137,M_{DBP}^{TWN}=190;$ $M_{SBP}^{EUR}=4379, M_{SBP}^{JPN}=174,M_{SBP}^{TWN}=195$. The bars represent the points estimate for $\hat{\theta}_{2}\pm$ standard error. Grey bars denote non-significant results, while colored bars highlight significant estimates. Asterisks indicate estimates with significant $I_{2}$. Positive (negative) and statistically significant value of $\hat{\theta}_{2}$implies that the sample is enriched for individuals with higher (lower) genetic propensity for the target trait compared to the reference.

| (a) | 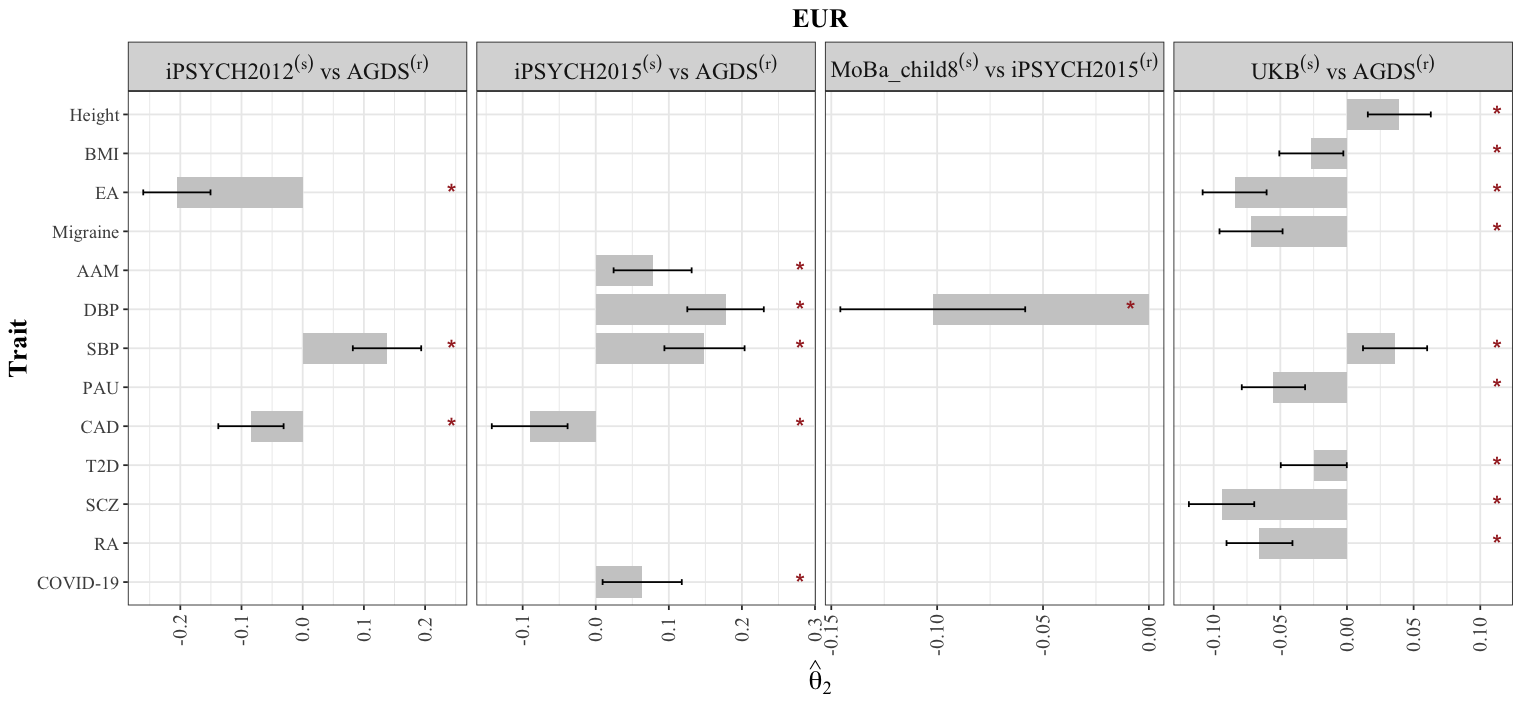 |
| --- | --- |
| (b) | 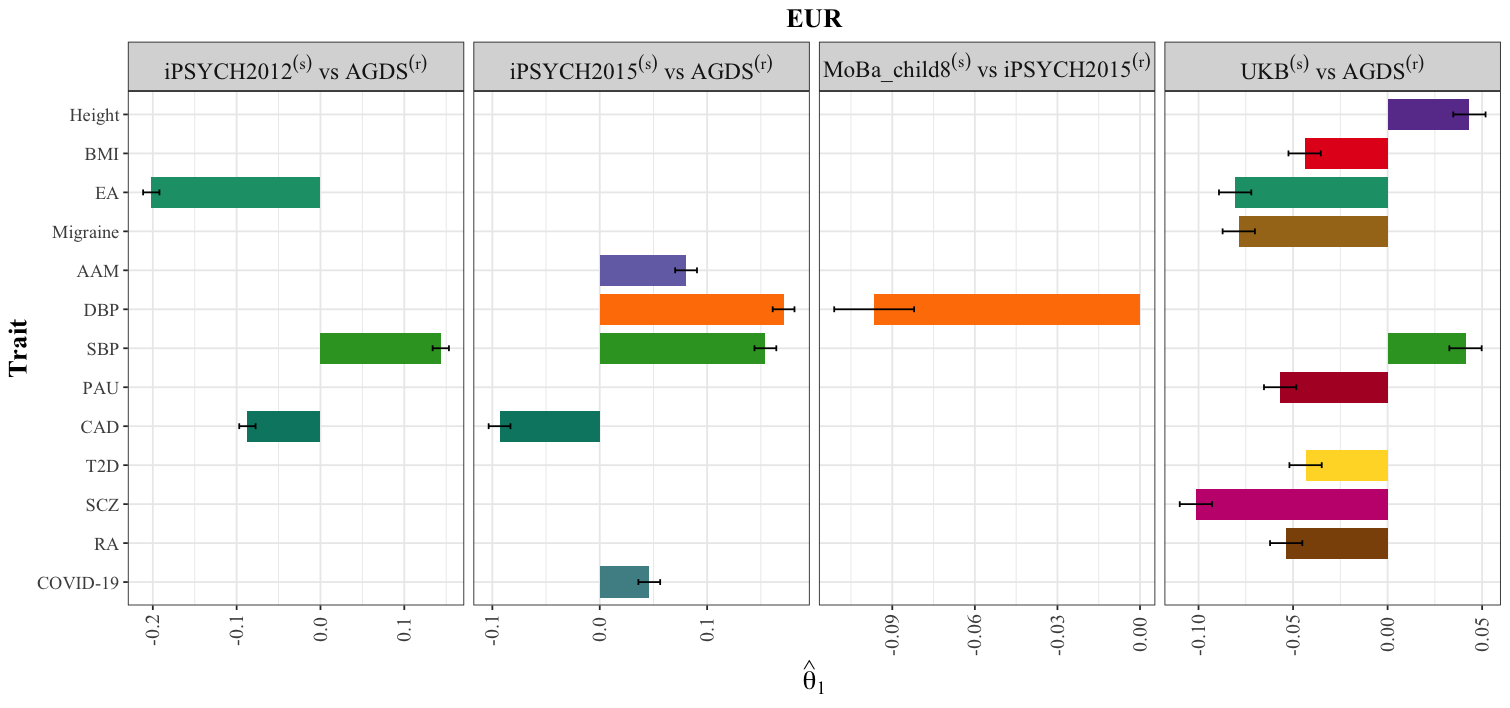 |

**Supplementary Fig. 21. Comparison of ascertainment signals obtained using** $\hat{\theta}_{2}$ **and** $\hat{\theta}_{1}$ **across within-European (EUR) ancestry biobank comparisons.** Panels show selected comparison for which $\hat{\theta}_{1}$ detected an ascertainment signal whereas $\hat{\theta}_{2}$ did not. **(a)** Relative ascertainment estimates obtained using the regression-based estimator $\hat{\theta}_{2}$ (top) and **(b)** the corresponding estimates obtained using mean PGS differences (${\hat{\boldsymbol{\theta}}}_{\boldsymbol{1}}$) (bottom). Bars represent point estimates of $\hat{\theta}_{2}$ ± standard error. Grey bars denote non-significant results, while coloured bars highlight statistically significant estimates after multiple-testing correction (*P* values < 0.0001). Asterisks indicate estimates with significant intercept (*I₂*; *P* values < 0.0001). Positive (negative) and statistically significant value of $\hat{\theta}_{2}$implies that the sample is enriched for individuals with higher (lower) genetic propensity or liability for the target trait compared to the reference cohort.

| (a) | 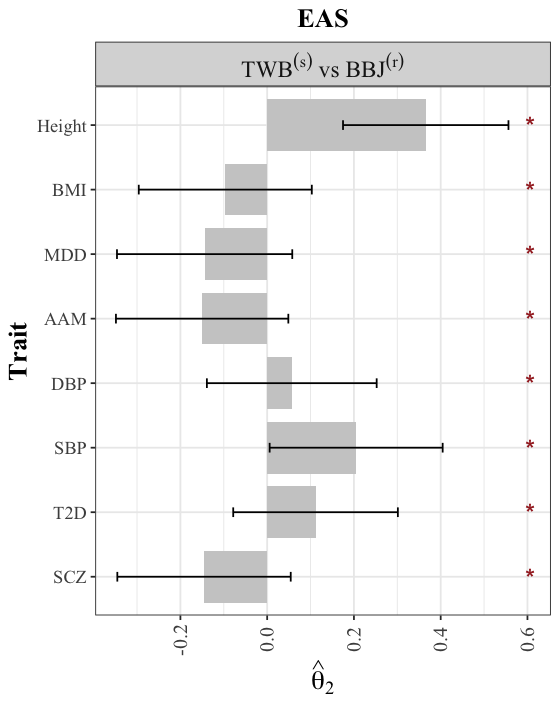 | (b) | 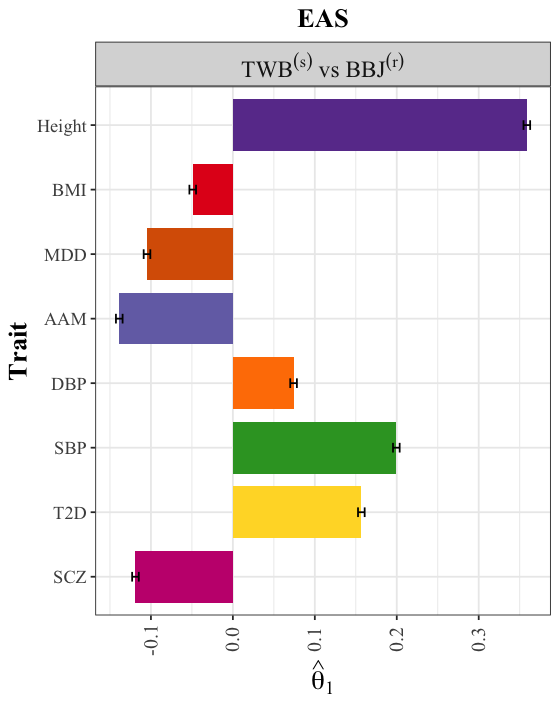 |
| --- | --- | --- | --- |

**Supplementary Fig. 22. Comparison of ascertainment signals obtained using** $\hat{\theta}_{2}$ **and** $\hat{\theta}_{1}$ **across within-East Asian (EAS) ancestry biobank comparisons.** Panels show selected comparison for which $\hat{\theta}_{1}$ detected an ascertainment signal whereas $\hat{\theta}_{2}$ did not. **(a)** Relative ascertainment estimates obtained using the regression-based estimator $\hat{\theta}_{2}$ (left) and **(b)** the corresponding estimates obtained using mean PGS differences (${\hat{\boldsymbol{\theta}}}_{\boldsymbol{1}}$) (right). Bars represent point estimates of $\hat{\theta}_{2}$ ± standard error. Grey bars denote non-significant results, while coloured bars highlight statistically significant estimates after multiple-testing correction (*P* values < 0.0001). Asterisks indicate estimates with significant intercept (*I₂*; *P* values < 0.0001). Positive (negative) and statistically significant value of $\hat{\theta}_{2}$implies that the sample is enriched for individuals with higher (lower) genetic propensity or liability for the target trait compared to the reference cohort.

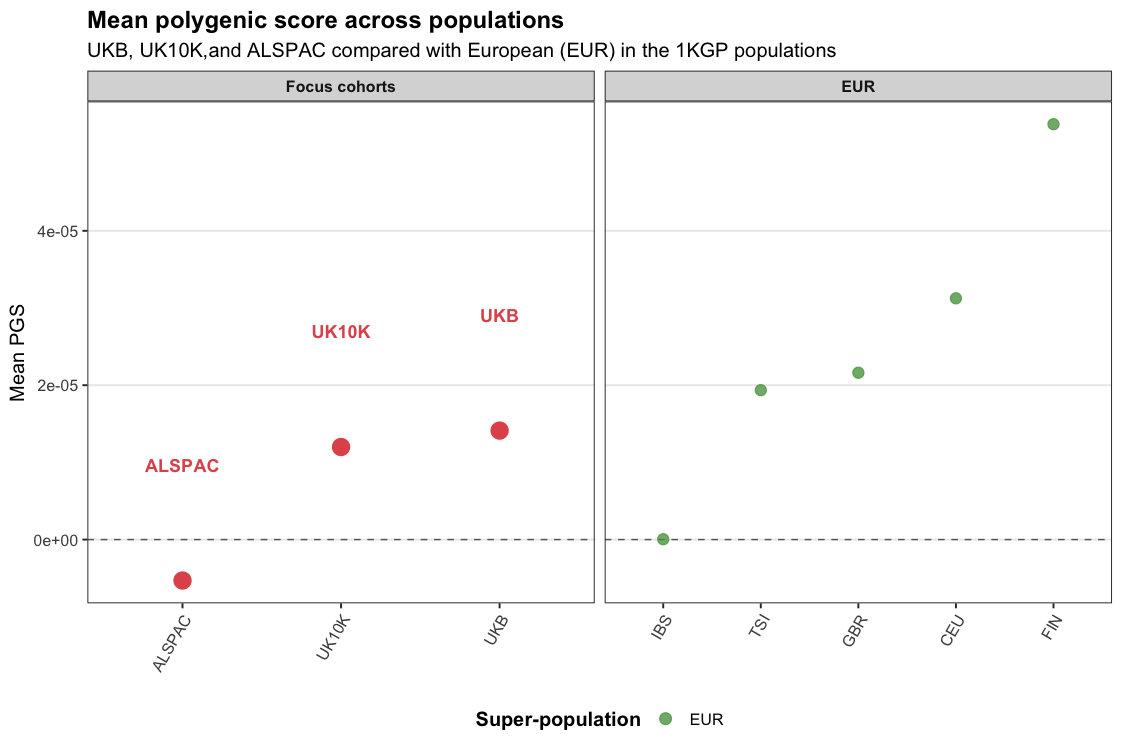

**Supplementary Fig. 23. Mean polygenic score for educational attainment (EA) across focus cohort and European ancestry in the 1000 Genomes Project.** Panel shows the mean polygenic scores (PGS) for educational attainment (EA) relative to the those from the European ancestry in the 1000 Genomes Project (1KGP). Each point represents the mean PGS within a population, with the dashed horizontal line indicating zero. Selected cohorts (UKB, UK10K and ALSPAC) are highlighted for comparison. Substantial variation in baseline EA PGS is observed across reference populations, with several groups exhibiting elevated mean PGS values.

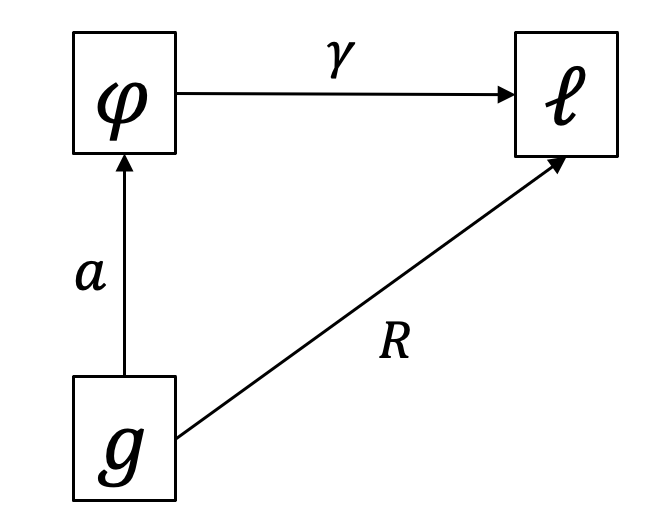

**Supplementary Fig. 24**. Path diagram of the liability threshold model linking the polygenic score (g), exposure ($\varphi$), and participation liability (ℓ). $a$ is the true genetic effect of 𝑔 on $\varphi$, $\gamma$ is the direct selection effect of $\varphi$ and $\mathcal{l}$ and R is the correlation between g and ℓ.

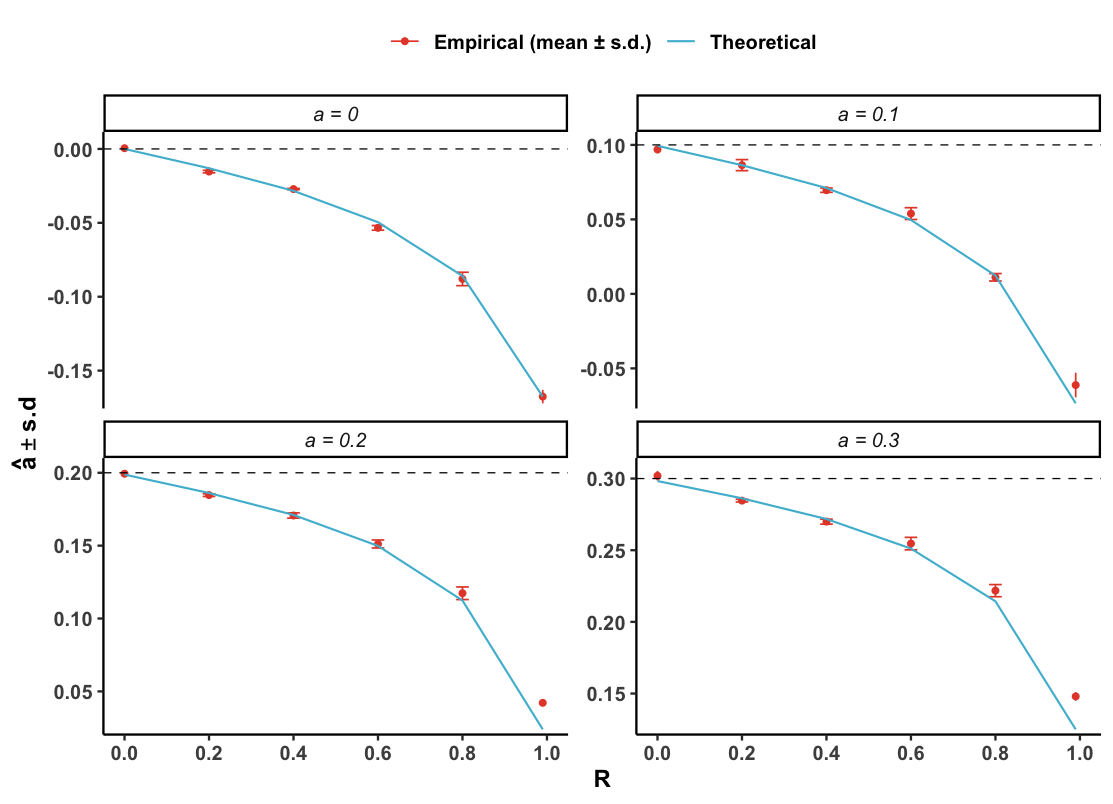

**Supplementary Fig. 25**. Theoretical and empirical bias in the regression of exposure ($\varphi$) on its polygenic score after selection. Panels show the predicted (blue line) and simulated (mean ± s.d.; with error bars) slope as a function of the correlation between polygenic score and the liability (R). The horizontal dashed line indicates the true genetic effect (a) in a non-ascertained sample. Parameters: $\varphi$ is fixed at 0.1.

### **SUPPLEMENTARY REFERENCES**

1.	Arriaga-MacKenzie IS, Matesi G, Chen S, Ronco A, Marker KM, Hall JR, et al. Summix: a method for detecting and adjusting for population structure in genetic summary data. The American Journal of Human Genetics. 2021;108(7):1270-82.

2.	Privé F. Using the UK Biobank as a global reference of worldwide populations: application to measuring ancestry diversity from GWAS summary statistics. Bioinformatics. 2022;38(13):3477-80.

3.	Bybjerg-Grauholm J, Bøcker Pedersen C, Bækvad-Hansen M, Giørtz Pedersen M, Adamsen D, Søholm Hansen C, et al. The iPSYCH2015 Case-Cohort sample: updated directions for unravelling genetic and environmental architectures of severe mental disorders. MedRxiv. 2020:2020.11. 30.20237768.

4.	Pedersen CB, Bybjerg-Grauholm J, Pedersen MG, Grove J, Agerbo E, Baekvad-Hansen M, et al. The iPSYCH2012 case–cohort sample: new directions for unravelling genetic and environmental architectures of severe mental disorders. Molecular psychiatry. 2018;23(1):6-14.

5.	Albiñana C, Zhu Z, Borbye-Lorenzen N, Boelt SG, Cohen AS, Skogstrand K, et al. Genetic correlates of vitamin D-binding protein and 25-hydroxyvitamin D in neonatal dried blood spots. Nature Communications. 2023;14(1):852.

6.	Mitchell BL, Campos AI, Whiteman DC, Olsen CM, Gordon SD, Walker AJ, et al. The Australian Genetics of Depression Study: new risk loci and dissecting heterogeneity between subtypes. Biological Psychiatry. 2022;92(3):227-35.

7.	Byrne EM, Kirk KM, Medland SE, McGrath JJ, Colodro-Conde L, Parker R, et al. Cohort profile: the Australian genetics of depression study. BMJ open. 2020;10(5):e032580.

8.	Sakaue S, Kanai M, Tanigawa Y, Karjalainen J, Kurki M, Koshiba S, et al. A cross-population atlas of genetic associations for 220 human phenotypes. Nature genetics. 2021;53(10):1415-24.

9.	Akiyama M, Ishigaki K, Sakaue S, Momozawa Y, Horikoshi M, Hirata M, et al. Characterizing rare and low-frequency height-associated variants in the Japanese population. Nature communications. 2019;10(1):4393.

10.	Leitsalu L, Haller T, Esko T, Tammesoo M-L, Alavere H, Snieder H, et al. Cohort profile: Estonian biobank of the Estonian genome center, university of Tartu. International journal of epidemiology. 2015;44(4):1137-47.

11.	Milani L, Alver M, Laur S, Reisberg S, Haller T, Aasmets O, et al. The Estonian Biobank’s journey from biobanking to personalized medicine. Nature communications. 2025;16(1):3270.

12.	Abner E, Batool K, Taba N, Nikopensius T, Läll K, Alekseienko A, et al. Characterization of prevalent genetic variants in the Estonian Biobank body-mass index GWAS. Nature Communications. 2025;16(1):8956.

13.	Wen S, Kuri-Morales P, Hu F, Nag A, Tachmazidou I, Deevi SV, et al. Comparative analysis of the Mexico City Prospective Study and the UK Biobank identifies ancestry-specific effects on clonal hematopoiesis. Nature Genetics. 2025;57(3):572-82.

14.	Sohail M, Palma-Martínez MJ, Chong AY, Quinto-Cortés CD, Barberena-Jonas C, Medina-Muñoz SG, et al. Mexican Biobank advances population and medical genomics of diverse ancestries. Nature. 2023;622(7984):775-83.

15.	Helgeland Ø, Vaudel M, Sole-Navais P, Flatley C, Juodakis J, Bacelis J, et al. Characterization of the genetic architecture of infant and early childhood body mass index. Nature Metabolism. 2022;4(3):344-58.

16.	Ameur A, Dahlberg J, Olason P, Vezzi F, Karlsson R, Martin M, et al. SweGen: a whole-genome data resource of genetic variability in a cross-section of the Swedish population. European Journal of Human Genetics. 2017;25(11):1253-60.

17.	Chen C-Y, Chen T-T, Feng Y-CA, Yu M, Lin S-C, Longchamps RJ, et al. Analysis across Taiwan Biobank, Biobank Japan, and UK Biobank identifies hundreds of novel loci for 36 quantitative traits. Cell genomics. 2023;3(12).

18.	Sudlow C, Gallacher J, Allen N, Beral V, Burton P, Danesh J, et al. UK biobank: an open access resource for identifying the causes of a wide range of complex diseases of middle and old age. PLoS medicine. 2015;12(3):e1001779.

19.	A reference panel of 64,976 haplotypes for genotype imputation. Nature genetics. 2016;48(10):1279-83.

20.	Yengo L, Vedantam S, Marouli E, Sidorenko J, Bartell E, Sakaue S, et al. A saturated map of common genetic variants associated with human height. Nature. 2022;610(7933):704-12.

21.	Yengo L, Sidorenko J, Kemper KE, Zheng Z, Wood AR, Weedon MN, et al. Meta-analysis of genome-wide association studies for height and body mass index in∼ 700000 individuals of European ancestry. Human molecular genetics. 2018;27(20):3641-9.

22.	Okbay A, Wu Y, Wang N, Jayashankar H, Bennett M, Nehzati SM, et al. Polygenic prediction of educational attainment within and between families from genome-wide association analyses in 3 million individuals. Nature genetics. 2022;54(4):437-49.

23.	Demontis D, Walters GB, Athanasiadis G, Walters R, Therrien K, Nielsen TT, et al. Genome-wide analyses of ADHD identify 27 risk loci, refine the genetic architecture and implicate several cognitive domains. Nature genetics. 2023;55(2):198-208.

24.	Friligkou E, Løkhammer S, Cabrera-Mendoza B, Shen J, He J, Deiana G, et al. Gene discovery and biological insights into anxiety disorders from a large-scale multi-ancestry genome-wide association study. Nature genetics. 2024;56(10):2036-45.

25.	O’Connell KS, Koromina M, Van Der Veen T, Boltz T, David FS, Yang JMK, et al. Genomics yields biological and phenotypic insights into bipolar disorder. Nature. 2025;639(8056):968-75.

26.	Adams MJ, Streit F, Meng X, Awasthi S, Adey BN, Choi KW, et al. Trans-ancestry genome-wide study of depression identifies 697 associations implicating cell types and pharmacotherapies. Cell. 2025;188(3):640-52. e9.

27.	Hautakangas H, Winsvold BS, Ruotsalainen SE, Bjornsdottir G, Harder AV, Kogelman LJ, et al. Genome-wide analysis of 102,084 migraine cases identifies 123 risk loci and subtype-specific risk alleles. Nature genetics. 2022;54(2):152-60.

28.	Nievergelt CM, Maihofer AX, Atkinson EG, Chen C-Y, Choi KW, Coleman JR, et al. Genome-wide association analyses identify 95 risk loci and provide insights into the neurobiology of post-traumatic stress disorder. Nature genetics. 2024;56(5):792-808.

29.	Kentistou KA, Kaisinger LR, Stankovic S, Vaudel M, Mendes de Oliveira E, Messina A, et al. Understanding the genetic complexity of puberty timing across the allele frequency spectrum. Nature genetics. 2024;56(7):1397-411.

30.	Ruth KS, Day FR, Hussain J, Martínez-Marchal A, Aiken CE, Azad A, et al. Genetic insights into biological mechanisms governing human ovarian ageing. Nature. 2021;596(7872):393-7.

31.	Keaton JM, Kamali Z, Xie T, Vaez A, Williams A, Goleva SB, et al. Genome-wide analysis in over 1 million individuals of European ancestry yields improved polygenic risk scores for blood pressure traits. Nature genetics. 2024;56(5):778-91.

32.	Zhou H, Kember RL, Deak JD, Xu H, Toikumo S, Yuan K, et al. Multi-ancestry study of the genetics of problematic alcohol use in over 1 million individuals. Nature Medicine. 2023;29(12):3184-92.

33.	Trajanoska K, Morris JA, Oei L, Zheng H-F, Evans DM, Kiel DP, et al. Assessment of the genetic and clinical determinants of fracture risk: genome wide association and mendelian randomisation study. bmj. 2018;362.

34.	Aragam KG, Jiang T, Goel A, Kanoni S, Wolford BN, Atri DS, et al. Discovery and systematic characterization of risk variants and genes for coronary artery disease in over a million participants. Nature genetics. 2022;54(12):1803-15.

35.	Suzuki K, Hatzikotoulas K, Southam L, Taylor HJ, Yin X, Lorenz KM, et al. Genetic drivers of heterogeneity in type 2 diabetes pathophysiology. Nature. 2024;627(8003):347-57.

36.	Trubetskoy V, Pardiñas AF, Qi T, Panagiotaropoulou G, Awasthi S, Bigdeli TB, et al. Mapping genomic loci implicates genes and synaptic biology in schizophrenia. Nature. 2022;604(7906):502-8.

37.	Herrera‐Luis E, Ortega VE, Ampleford EJ, Sio YY, Granell R, de Roos E, et al. Multi‐ancestry genome‐wide association study of asthma exacerbations. Pediatric allergy and immunology. 2022;33(6):e13802.

38.	Ishigaki K, Sakaue S, Terao C, Luo Y, Sonehara K, Yamaguchi K, et al. Multi-ancestry genome-wide association analyses identify novel genetic mechanisms in rheumatoid arthritis. Nature genetics. 2022;54(11):1640-51.

39.	Kanai M, Andrews SJ, Cordioli M, Stevens C, Neale BM, Daly M, et al. A second update on mapping the human genetic architecture of COVID-19. Nature. 2023;621(7977):E7-E26.
